# The Evolutionary History of Common Genetic Variants Influencing Human Cortical Surface Area

**DOI:** 10.1101/703793

**Authors:** Amanda K. Tilot, Ekaterina A. Khramtsova, Katrina Grasby, Neda Jahanshad, Jodie Painter, Lucía Colodro-Conde, Janita Bralten, Derrek P. Hibar, Penelope A. Lind, Siyao Liu, Sarah M. Brotman, Paul M. Thompson, Sarah E. Medland, Fabio Macciardi, Barbara E. Stranger, Lea K. Davis, Simon E. Fisher, Jason L. Stein

## Abstract

Structural brain changes along the lineage that led to modern *Homo sapiens* have contributed to our unique cognitive and social abilities. However, the evolutionarily relevant molecular variants impacting key aspects of neuroanatomy are largely unknown. Here, we integrate evolutionary annotations of the genome at diverse timescales with common variant associations from large-scale neuroimaging genetic screens in living humans, to reveal how selective pressures have shaped neocortical surface area. We show that variation within human gained enhancers active in the developing brain is associated with global surface area as well as that of specific regions. Moreover, we find evidence of recent polygenic selection over the past 2,000 years influencing surface area of multiple cortical regions, including those involved in spoken language and visual processing.

## Introduction

The distinctive size, shape, and neural architecture of the modern human brain reflects the cumulative effects of selective pressures over evolutionary history. Analyses of fossilized skulls indicate that endocranial volume has increased dramatically on the lineage that led to *Homo sapiens* in the ∼6 million years since our last common ancestor with chimpanzees^1–4^ (**Figure 1**). It is thought that these volumetric increases were driven in large part by expansions of neocortical surface area^5–7^, although changes in other brain structures, including the cerebellum, also likely played a significant role^8, 9^. Beyond overall size differences, skull endocasts of archaic hominins suggest that human-specific refinements to brain structure occurred during the last 300,000 years, most notably the shift toward a more globular shape^10, 11^. Changes in neuroanatomy in our ancestors were accompanied by increasingly sophisticated tool use, the emergence of proficient spoken language, world-wide migrations, and the development of agriculture, amongst other innovations^12^.

**Figure 1.**
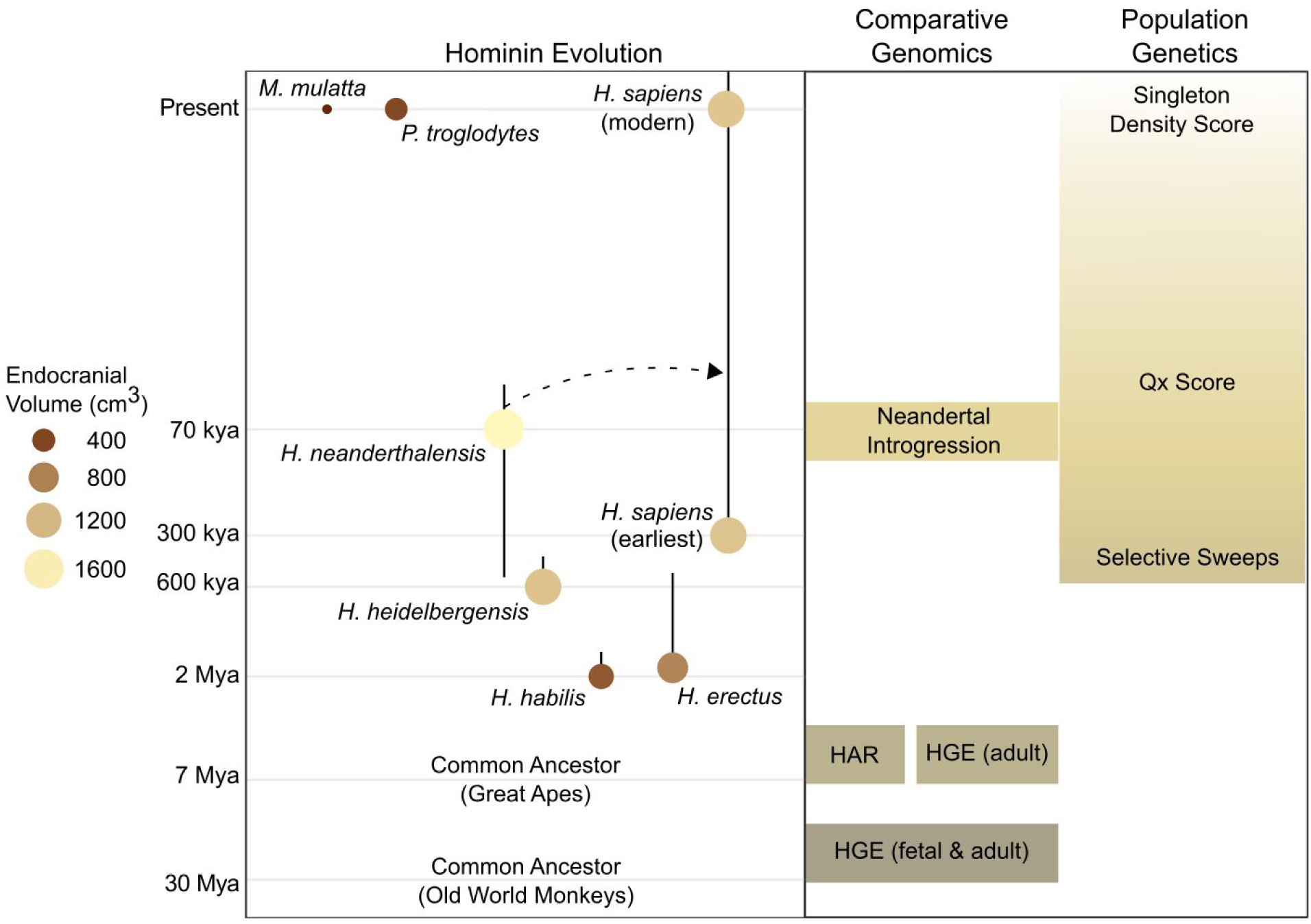
Overview of human evolution and timeframe captured by each set of analyses in this study. Size and shading of circles indicates average endocranial volume, vertical lines reflect the approximate timeframe of the hominin species (*left*)^59^. Evolutionary time is presented in log10 (Mya, million years ago; kya, thousand years ago). Different types of evolutionary annotations are indicated, identifying genomic loci that underwent changes over different time frames (*right*). HAR, human accelerated regions. HGE, human gained enhancers. See also Table S1.

A commonly held view is that differential expansion of distinct regions of the neocortex contributed to the evolution of the unique cognitive and social abilities of our species^5–7^. Consistent with this idea, comparative studies show that cortical surface area is greatly expanded in humans relative to extant non-human primates, in an areal-dependent manner^5, 13, 14^. Investigations of interindividual variation within current human populations have found substantial overlaps in the genomic architecture underlying endocranial volume and total neocortical surface area, while also supporting the existence of differential genetic effects on the surface area of specific cortical regions^15, 16^. A few notable studies have identified genomic differences that may have impacted aspects of brain structure along our lineage^13, 17, 18^, but the genetic variation that shaped the cortex across human evolution is still largely undetermined. In the present study, we adopt a novel strategy to uncover genetic variants that have contributed to anatomical features of the modern human brain. To do so, we identify loci of defined evolutionary relevance in the genome and assess the effects of those loci on cortical structure through large-scale neuroimaging genetics.

Two complementary approaches are typically used to assess evolutionary relevance of regions of the human genome: (1) *comparative genomics*, in which the sequence and/or regulatory functions of the human genome are compared to other extant primate species or extinct archaic hominins like Neanderthals and Denisovans, and (2) *population genetics*, which involves analyzing patterns of genetic variations in existing human populations to detect evidence of selection^19^. The methods available for each approach allow annotations of evolutionary relevance across different time scales of human evolution^19^ (**Figure 1**).

Here, we use comparative genomic annotations from three sources that may shed light on evolutionary changes in cortical structure. <u>Human Gained Enhancers</u> (HGEs) are gene regulatory elements that display stronger histone acetylation or methylation marks of promoters or enhancers in human cortical tissue compared to extant primates or mice^20, 21^. We can infer that these differences arose after our most recent common ancestor with the comparison species, e.g. Old World monkeys at about 30 million years ago (Mya). <u>Human Accelerated Regions</u> (HARs) are sections of the genome that are highly conserved in extant vertebrate species, but show an acceleration of substitutions along the human lineage^22^, after our most recent common ancestor with chimpanzees, around 6 Mya. <u>Neanderthal introgressed fragments</u> are rare stretches of Neanderthal DNA that can be detected in the genomes of non-African individuals in current populations. These are remnants of interbreeding events that occurred at various stages, most recently ∼50-65 kya (thousand years ago)^23^, in populations that migrated out of Africa. Neanderthal introgression has been associated with variation in several traits in modern humans, including risk of depression^24, 25^.

We additionally investigate three types of population genetic annotations that may be relevant to changes in cortical structure. <u>Selective sweep regions</u> are exceptionally long genomic regions of low diversity resulting from nearby selected alleles quickly rising in frequency to become fixed (i.e., non-varying) in the population^26^. One method for identifying such sweeps, Extended Lineage Sorting, can be used to detect putative selective events occurring over the last ∼300-600 kya^26^. Moving to more recent periods of evolutionary history, the <u>Qx score</u> determines if alleles at multiple loci associated with a multifactorial trait also display consistent differences in allele frequencies across existing human populations, which is taken as a signature of polygenic adaptation occurring following the separation of those populations (∼50 kya)^27^. Finally, the <u>Singleton Density Score</u> (SDS) uses genome sequencing data to identify haplotypes (linked clusters of alleles at neighbouring markers) that show a decreased accumulation of singleton mutations in the population being studied, providing evidence for natural polygenic selection acting over the past ∼2,000-3,000 years^28^.

By themselves, these indices can point to loci of likely evolutionary significance in the human genome, but they are not informative with respect to which loci (if any) influence the structure of the human brain. We hypothesize that, for evolutionarily relevant genetic variants that have not reached fixation in current human populations, data from genome-wide association studies (GWAS) of cortical structure can help determine their potential functional impact on brain structure. Moreover, we reason that GWAS data may shed light on the evolution of cortical structure by: (1) revealing if interindividual variation in defined genomic regions of evolutionary significance are associated with variation in neural anatomy, and (2) determining if alleles under selective pressure are associated with variation in neural anatomy. Crucially, this novel approach for studying human brain evolution depends on the availability of large datasets of many thousands of individuals in which measures of brain structure (characterized through non-invasive neuroimaging, for example) have been coupled to genome-wide genotyping. In this regard, we take advantage of recent large-scale GWAS work from the Enhancing NeuroImaging Genetics through Meta Analysis (ENIGMA) consortium^16^ and the UK Biobank^29^, which have identified hundreds of genetic loci associated with interindividual variability in human cortical structure in living populations. Thus, in the present study, we integrate multiple genomic annotations spanning 30 million years of our evolutionary history with data from a GWAS meta-analysis of cortical surface area (SA) in over 35,000 modern humans^16^ to (1) assess the aggregate impact of each annotation on modern variation in cortical SA and (2) identify specific genetic variants within these annotations with effects on human neural development.

## Results

### Genome-wide association to human cortical surface area

The ENIGMA consortium recently conducted a GWAS meta-analysis to identify common variants associated with variability in SA in European populations^16^. A similar GWAS meta-analysis was also conducted for cortical thickness, but given the massive expansion of SA in modern humans and only subtle increases in cortical thickness as compared to extant mammalian species^5^, we chose to focus on SA for this study. Association was assessed in 35,660 individuals from 55 cohorts across the lifespan, including data from the UK Biobank^16^ (UKBB, N = 9,923). From standard structural magnetic resonance imaging (MRI) scans of the brain, the ENIGMA consortium automatically measured total SA across the cortex, as well as average bilateral SA for each of 34 cortical regions defined by gyral anatomy^30^. Genetic associations were conducted within each cohort, separately. When conducting association tests of gyrally defined regions, the global measure of SA was included as a covariate, to test for genetic influences specific to each region. The association models also included four multi-dimensional scaling components to control for ancestry, linear and non-linear corrections for age and sex, diagnostic status and scanner (if applicable; see Online Methods). Summary statistics were then combined across cohorts using a fixed effects meta-analysis framework^31^.

### Reducing the impact of subtle population stratification

Population stratification is the existence of systematic differences in allele frequencies between populations. Unbalanced representations of multiple populations in genetic association studies can lead to false-positive findings that are driven by allele frequency differences between populations rather than true association with a trait^32^. The effects of subtle population stratification on GWAS statistics can inflate the assessment of polygenic selection impacting a trait^33–36^. We first tested whether subtle population stratification was influencing meta-analysis effect sizes in the cortical GWAS data, even after standard correction for multi-dimensional scaling components of ancestry prior to meta-analysis^16^. Principal component (PC) analysis enabled us to identify major axes of variation in allele frequency across current human populations using unrelated individuals of all ancestries from the 1000 Genomes Phase 3 data. Then, we tested the association of each SNP to the top 20 PCs (each treated as a separate trait) within the 1000 Genomes population, yielding an estimate of the degree to which each SNP contributes to population frequency differences along each principal axis of variation (Beta_PCs). Finally, using Pearson’s correlation the Beta_PCs were correlated with the effect sizes from the GWAS meta-analysis for each trait, which may be impacted by population stratification (Beta_Strat). To assess the significance of the correlation in the context of linkage disequilibrium (LD), a block jackknife approach was employed to calculate the standard errors for the correlation^37, 38^. Significant correlations between Beta_PCs (consistent allele frequency differences differentiating human populations) and Beta_Strat (effect sizes of variants on human brain structure from GWAS) are indicative of subtle, uncorrected population stratification^35, 36^. As shown in **Figure 2A**, we detected significant relationships between Beta_Strat and PCs 6 and 18 for global SA, indicating subtle residual population stratification affecting the GWAS summary statistics. This analysis also showed subtle population stratification affecting summary statistics for each of the regional SAs, to varying degrees (**Figure S1**). We note that another measure of population stratification, the LD-score regression intercept^39^, gave values that were uniformly less than 1.05 (a commonly used threshold for ruling out stratification) for global SA and all regional SAs (**Figure 2C**).

**Figure 2.**
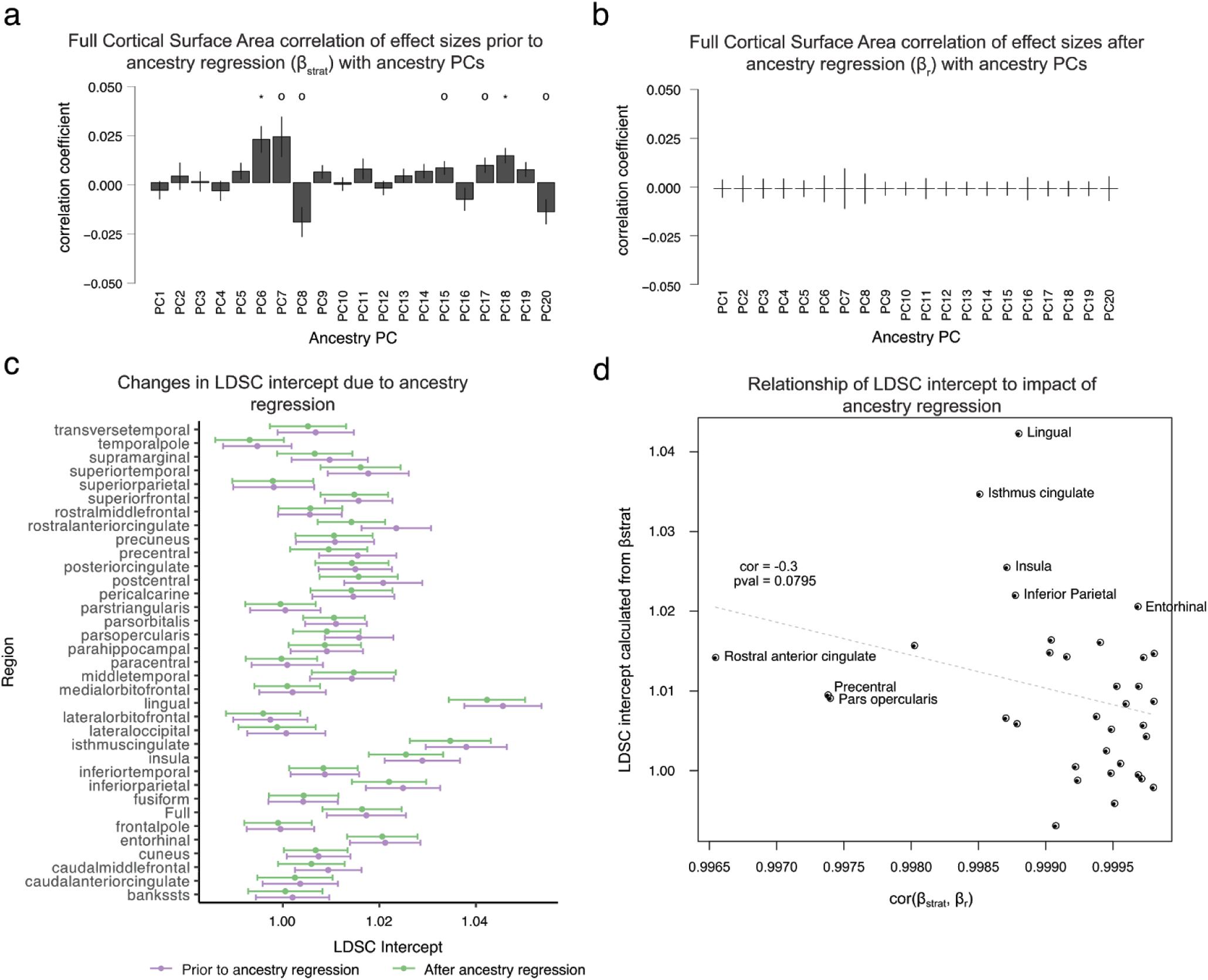
Identifying and correcting the effects of subtle population stratification on GWAS results. (a) Correlations between SNP loadings on ancestry PCs (Beta_PCs) and GWAS effect sizes for full surface area (Beta_Strat) demonstrate evidence for subtle population stratification (* indicates Bonferroni corrected significant correlation *p*-value < 0.0025, and o indicates a nominally significant correlation, *p*-value < 0.05). **(b)** Subtle population stratification is reduced after ancestry regression. **(c)** LD-score regression (LDSC) intercepts, standard measures of population stratification, are generally decreased after ancestry regression. **(d)** There is an inverse relationship between the degree of subtle population stratification (LDSC intercept prior to ancestry regression) and the amount of change caused by ancestry regression (cor(Beta_strat, Beta_r)). Error bars represent standard errors. See also Figures S1-2.

To correct for this subtle population stratification, we implemented an ancestry regression procedure based on GWAS summary statistics^40^. The residuals (Beta_r) of a model fitting GWAS effect sizes (Beta_Strat) with the first 20 PC weightings (Beta_PC) were used as ancestry-corrected estimates of effect sizes. As expected, these ancestry-corrected estimates (Beta_r) showed much reduced correlations with PC weights (**Figure 2B**, **Figure S2**). Additionally, LD-score regression intercepts for each phenotype after ancestry regression were slightly decreased, consistent with diminished effects of subtle population stratification on common variant associations to SA (**Figure 2C**). Furthermore, correlations between effect size measurements after ancestry regression (Beta_r) and effect size measurements prior to ancestry regression (Beta_Strat) were all extremely high, indicating that ancestry regression did not strongly change the association statistics (correlations all > 0.995; **Figure 2D**). There were 287 genome-wide significant loci (*p*-values < 5×10^−8^; clumping r^2^ < 0.2) prior to ancestry regression impacting global SA or any of the regional SAs and 284 genome-wide significant loci after ancestry regression. Finally, those brain regions that showed the highest LD-score regression intercepts, indicative of being most affected by subtle residual population stratification, were also those that showed the largest changes in GWAS effect sizes following ancestry regression (r = −0.30; *p*-value = 0.0795; **Figure 2D**). For all subsequent analyses, we used the ancestry-corrected effect size estimates, standard errors, and *p*-values, thereby minimizing the impact of population stratification on the results of our evolutionary assessments.

### Significant heritability enrichment within Human Gained Enhancers (30 Mya)

For our first analyses of evolutionarily relevant loci, we targeted human fetal brain enhancer elements that emerged since our last common ancestor with macaques, termed *human gained enhancers* (HGEs)^20^. These elements were detected by comparing post-translational modifications of histone tails indicative of enhancers and promoters (H3K27ac and H3K4me2) across humans, macaques, and mice. Using brain tissue from similar developmental time points across the three species, regulatory elements (peaks in the histone modification signals) were identified that were present in human fetal brain at 7 post-conception weeks (PCW), but to a significantly lesser degree in developing macaque or mouse brain tissue^20^. To understand how these HGEs influence cortical SA in modern humans, we measured their relative contribution to total *SNP heritability*. A trait’s SNP-based heritability is the total amount of variance in the trait (e.g., global surface area) that can be attributed to common variation across the genome, and it can be estimated from GWAS summary statistics. This genome-wide SNP heritability can be partitioned into functional or structural categories to measure how specific genomic regions of interest (in this case, evolutionary annotations) contribute to the heritability of the trait.

We assessed how common variants within HGEs contribute to the SNP-based heritability of cortical SAs, testing for enrichment using LD-score regression partitioned heritability^41^, with false-discovery-rate (FDR) correction for the 35 GWAS (34 regions plus global SA). Further, because SNPs within regulatory elements that are active during fetal development are known to make significant impacts on both intracranial volume and cortical SA^16, 42^, we controlled for a global category of fetal brain active regulatory elements (derived from the Epigenomics Roadmap^43^) in the analysis. SNPs within HGE elements made significantly enriched contributions to cortical SA heritability, for global SA (Enrichment = 9.94, FDR corrected *p*-value = 0.012), and also for 26 out of 34 gyrally-defined regions after controlling for global SA (**Figure 3a**). The enrichment signal was strongest for the *pars orbitalis*, part of the inferior frontal gyrus (Enrichment = 43.29, FDR corrected *p*-value = 5.6 x 10^−4^). As the regional GWAS results were controlled for global SA, the heritability enrichment signals detected in each region are independent of global SA. Our findings indicate that SNPs within these HGEs have effects beyond those of general fetal enhancers. Together, the data suggest that a key set of neural enhancer regions that became functional since our split from Old World monkeys contribute an unusually large amount to the heritability of both regional and global cortical SA in adult humans. This influence on SA in the adult brain may be realized through common genetic variation within these HGEs impacting gene regulation during fetal brain development.

**Figure 3.**
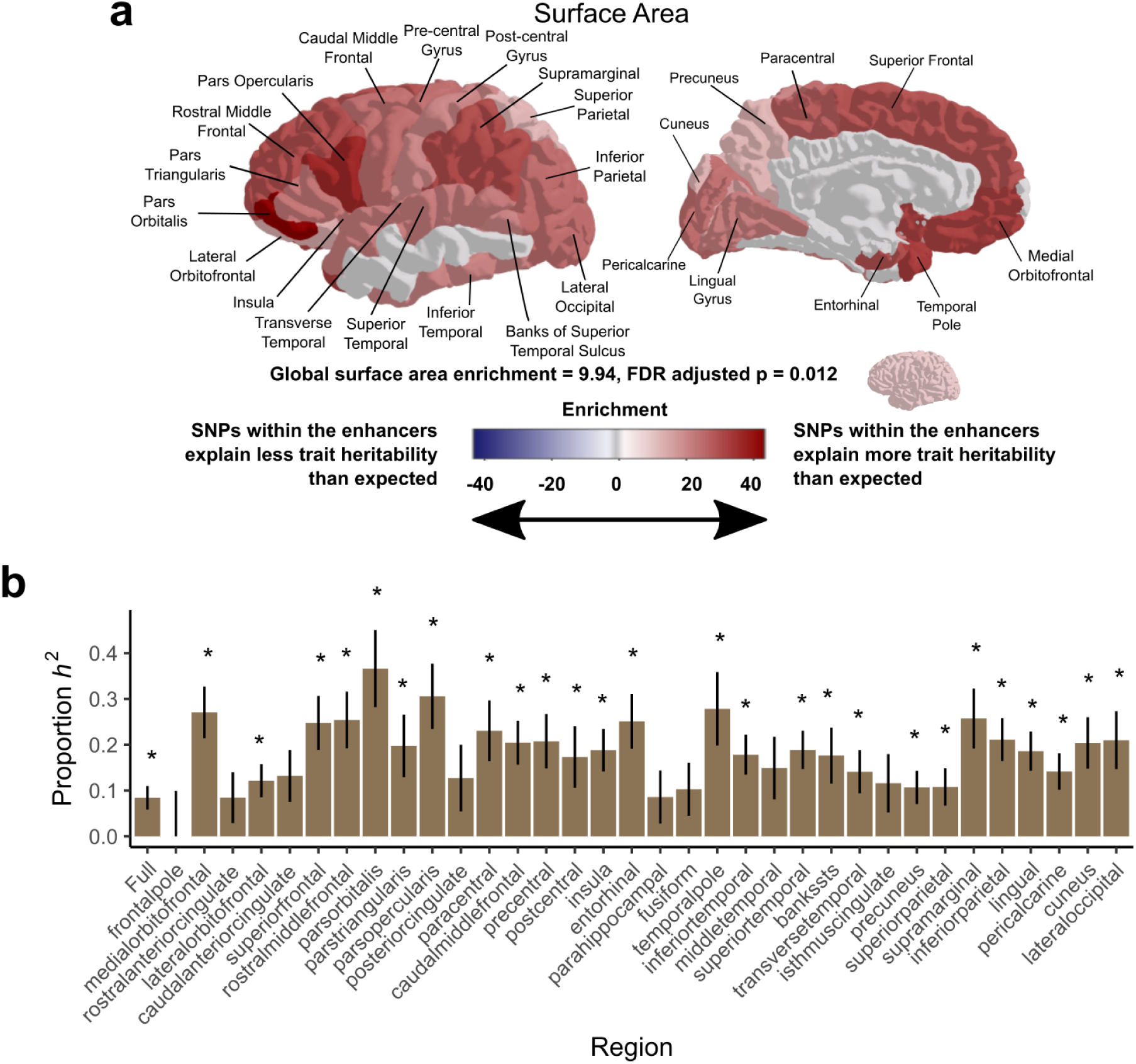
SNPs within human gained enhancers active at 7 weeks post conception explain a significant proportion of the heritability of surface area for several brain regions. (a) Shading indicates the enrichment values of SNP-heritability explained by human gained enhancers active at 7 weeks post conception for the surface area of each region, non-significant values are shaded in grey. **(b)** Proportion of SNP-heritability explained by 7 PCW HGEs for surface area of each cortical region. Asterisks label regions with FDR-corrected *p*-values < 0.05, error bars represent standard errors. See also Figure S4.

### Derived alleles within HGEs are associated with cumulative increases in SA

Partitioning heritability via LD-score regression^41^ identifies categories of SNPs that explain variation in a given quantitative trait. However, this method cannot determine directionality - whether alleles of interest are associated with increases or decreases in the trait. In this case, we were specifically interested in the potential roles of alleles that are *derived*, meaning that they arose on the human lineage after divergence from other primate species, asking whether they are associated with increases or decreases of SA measures. To determine how derived alleles within fetal brain HGEs impact cortical SA in aggregate, we generated a *polygenic score* (PGS) from the GWAS meta-analysis data. GWAS effects (already corrected for subtle population stratification, as above) were aligned to the Ensembl-defined derived alleles, which are based on alignment of six primate species (human, chimpanzee, gorilla, orangutan, macaque, and marmoset)^44^. After scaling the derived allele-aligned effects by their standard errors, we generated a PGS for the fetal brain HGEs by summing the scaled effects of all LD-independent SNPs within each element that met a *p*-value threshold of 0.005 in the GWAS data, and dividing by the total number of SNPs passing the threshold (**Figure 4A**). To evaluate the robustness of the primary PGS results, we tested additional *p*-value thresholds ranging from 0.0005 to 0.5. While the strength of the signals degraded with more lenient thresholds, the direction of effects generally remained the same (**Figure S3**).

**Figure 4.**
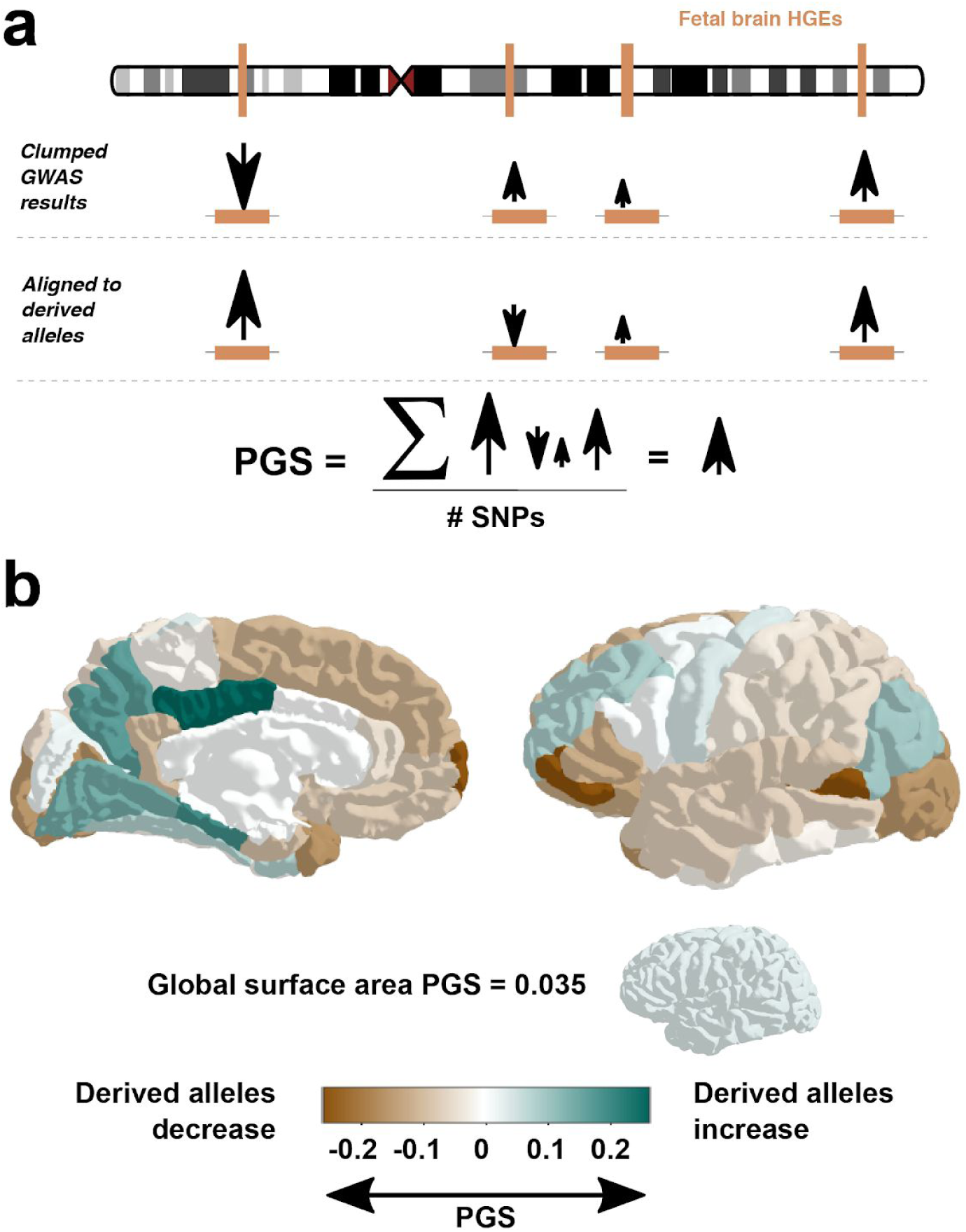
Directionality of effect of derived alleles at human gained enhancers on cortical surface area. (a) Schematic describing the construction of polygenic scores based on averaging effect sizes of LD-independent derived alleles within regions of interest. **(b)** Polygenic scores for cortical surface area, constructed from derived alleles within human gained enhancer elements (7 PCW HGEs). PGS = polygenic score. See also Figure S3.

The PGS for global SA (*p*-value threshold = 0.005 yielding 266 LD-independent SNPs) was 0.035, indicating that derived alleles within the fetal brain HGEs are associated with increases in global SA in GWAS data, when considered in aggregate. This finding is consistent with the known expansion of human global SA compared to macaque^14^ and further along our lineage^4^. Across the 34 gyrally-defined regions, there was considerable variability in whether the derived alleles, in aggregate, showed association with increases or decreases in regional SA, controlling for global SA (**Figure 4B**). To help interpret this heterogeneity, we looked to regional differences in cortical surface areas (*arealization*) between humans and macaques, based on registration of structural MRI scans between the species^14^. Beyond the global increase in cortical SA, regions where derived alleles within HGEs associate with increased SA in humans (shown in green in **Figure 4B**) were not obviously aligned with regions that have the highest levels of expansion compared to macaque, namely the dorsolateral prefrontal cortex and temporo-parietal junction^14^. However, in the reverse direction, the lateral occipital cortex has very low relative expansion in humans compared to macaques, and the derived alleles within HGEs collectively pointed toward decreased SA. It is important to note that arealization differences between human and macaque are built upon genetic variation accumulated since our very distant last common ancestor. Further, unlike the GWAS data used here, the comparative neuroanatomical maps do not control for the difference in total cortical SA^14^.

### Other classes of evolutionary annotations are not enriched for cortical SA heritability

We examined the contributions of several other evolution-focused annotations (e.g. adult HGEs, HARs, selective sweeps, and Neanderthal introgressed regions) to the heritability of cortical SA, finding no positive enrichment (**Figure S4**). This suggests that these particular sets of genomic regions do not contribute more to the heritability of cortical SA than expected, given their size.

### Linking GWAS results, genes, and evolutionary history

To further understand genes regulated impacted by common variation within HGEs, we established which of the genome-wide significant SA loci (*p*-value < 5×10^−8^ including SNPs in LD at *r^2^* > 0.6) fall within HGEs and also impact gene expression in adult cortical tissue^45^ (expression quantitative trait loci - eQTLs - at FDR < 0.05). Five of 22 genome-wide significant global SA loci overlapped (directly or with an LD-associated SNP) with a 7 PCW HGE. Three of those 5 loci also have a significant eQTL impacting 18 protein-coding *eGenes*, defined as the genes whose expression is associated with the genetic variation. These eGenes included developmentally relevant genes *FOXO3*, *ERBB3* and *WNT3* (a full list is found in **Table S2**).

Two SNPs in LD with rs2802295 (rs7772662, *r^2^* = 0.227; rs9400239, *r^2^* = 0.715) associated with global SA map to a 7 PCW fetal brain HGE as well as a Neanderthal introgressed region, and are located within an intron of the *FOXO3* gene on chromosome 6q21 (**Figure 5a**). rs2802295 has also been associated with interindividual variation in general intelligence^46^ (marked by rs2490272 index SNP, *r^2^* = 1.0 with rs2802295), with the SA increasing allele also associated with higher scores on tests of intelligence. The SNP also functions as a cortical eQTL for *FOXO3* (FDR adjusted *p*-value=0.0051, derived from the adult brain PsychENCODE dataset^45^), a transcription factor that regulates neuronal stem cell homeostasis^47^, amongst other roles.

**Figure 5.**
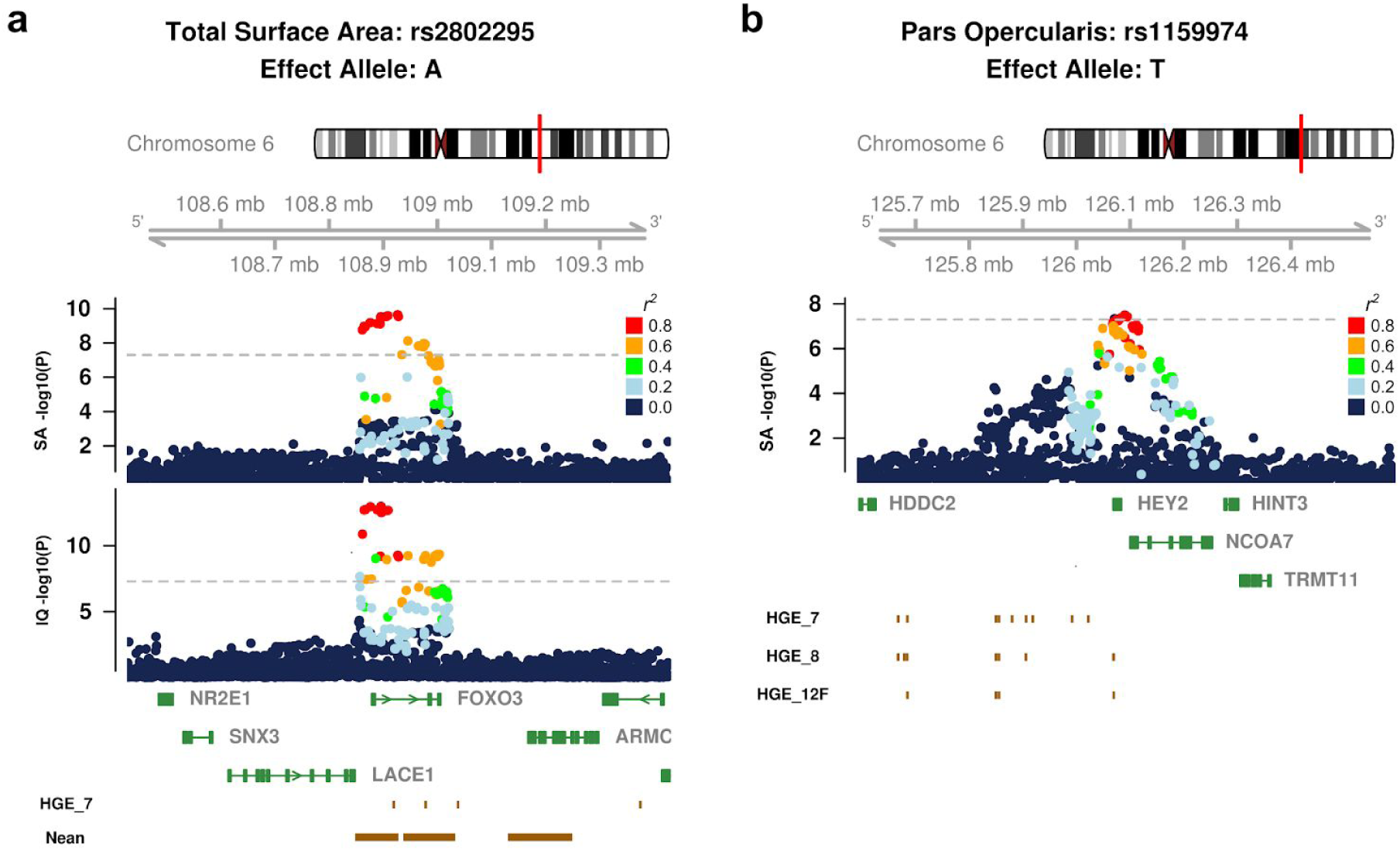
Loci associated with cortical surface area overlap evolutionary annotations and are associated with diverse biological functions. (a) Regional plots showing associations with rs2802295 and global surface area (upper) and IQ^46^ (lower) and **(b)** rs1159974 with *pars opercularis* surface area. Tracks depict the location of nearby genes and overlapping evolution-focused annotations. See also Figure S5.

Considering the 228 genome-wide significant regional SA loci, there were 31 that overlapped (directly or with an LD-associated SNP) with a 7 PCW HGE. Out of those 31 loci, 23 also have a significant eQTL, impacting a total of 42 protein-coding eGenes. These eGenes include known genes involved in areal identity including *LMO4*^48^ (a full list is found in **Table S2**). Another gene impacted by genetic variation within HGEs is *HEY2*. Six SNPs in LD with rs1159974 associated with *pars opercularis* SA map to a 7 PCW fetal brain HGE (rs12527010, *r^2^* = 0.284; rs9491471, *r^2^* = 0.244; rs926854, *r^2^* = 0.244; rs926855, *r^2^* = 0.244; rs1739364, *r^2^* = 0.274; rs1777220, *r^2^* = 0.338), with the locus centered on the promoter of the *HEY2* gene on chromosome 6q22 (**Figure 5b**). This SNP also functions as a cortical eQTL for *HEY2* (FDR adjusted *p*-value = 1.47×10^−41^ derived from the adult brain PsychENCODE dataset^45^), which regulates neural progenitor proliferation during neurogenesis^49^.

Likely due to the limited number of eGenes identified, no significant (FDR < 0.05) gene ontology terms with greater than 5 intersections with HGE regulated genes were identified. Nevertheless, this analysis points to specific developmentally interesting genes regulated by HGEs which have shaped both the overall surface area of the cortex and specific regions. Plots of all genome-wide significant loci that overlapped with one or more of the evolutionary annotations considered in this study are provided in **Figure S5**.

### Evidence for Polygenic selection over recent history impacting brain structure

To capture the effects of allele frequency changes over more recent timescales, we applied two measures that are taken to reflect selective pressures in humans over either ∼50 kya (Qx^27^) or ∼2-3 kya (singleton density scores, SDS^28^). The Qx score is used to infer polygenic selection over the past ∼50,000 years, testing whether polymorphic markers associated with a trait display more variation in allele frequencies across current human populations than would be expected based on drift. Using LD-clumped ancestry regressed GWAS summary statistics thresholded at a significance value of *p* < 1×10^−4^ for each human cortical region, as well as global SA, we did not detect significant variation in allele frequencies for the associated SNPs across current human populations. Thus, our analyses did not find evidence for polygenic adaptation to local selective pressures acting on cortical SA traits on this ∼50,000 year timescale (**Table S3**).

Finally, we assessed how alleles that show evidence of very recent selective pressure impact cortical SA. The singleton density score (SDS) reveals haplotypes under recent positive/negative selection in the human genome by identifying those that harbor fewer/greater singleton variants (presumed to have arisen recently) near any given SNP^28^. This metric, together with data from a suitable GWAS, can be used to infer whether a trait of interest has been subject to highly polygenic selection on a recent timescale, over the past ∼2,000-3,000 years. As with all other analyses in this study, we worked with GWAS data that had undergone an ancestry regression procedure. We found that alleles with evidence of positive selection over the relevant timescale have a small but detectable influence on increasing global SA in the GWAS datasets (block jackknife correlation = 0.0091, FDR adjusted *p*-value = 0.0090; **Figure 6A**). In addition, our results showed that alleles undergoing polygenic selection over the past ∼2,000-3,000 years are associated with variation in cortical SA of individual gyrally-defined brain regions (**Figure 6A**). Notably, based on the cortical region-specific GWASs, there is a detectable relationship between alleles under positive polygenic selective pressure (increasing in allele frequency over time) and increased cortical SA in regions known to be important for speech/language functions (*pars opercularis*) and visual processing (lateral occipital cortex). Conversely, alleles under negative polygenic selective pressure (decreasing allele frequency over time) are associated with increased cortical SA in the pre- and post-central gyrus, regions involved in somatosensation and movement.

**Figure 6.**
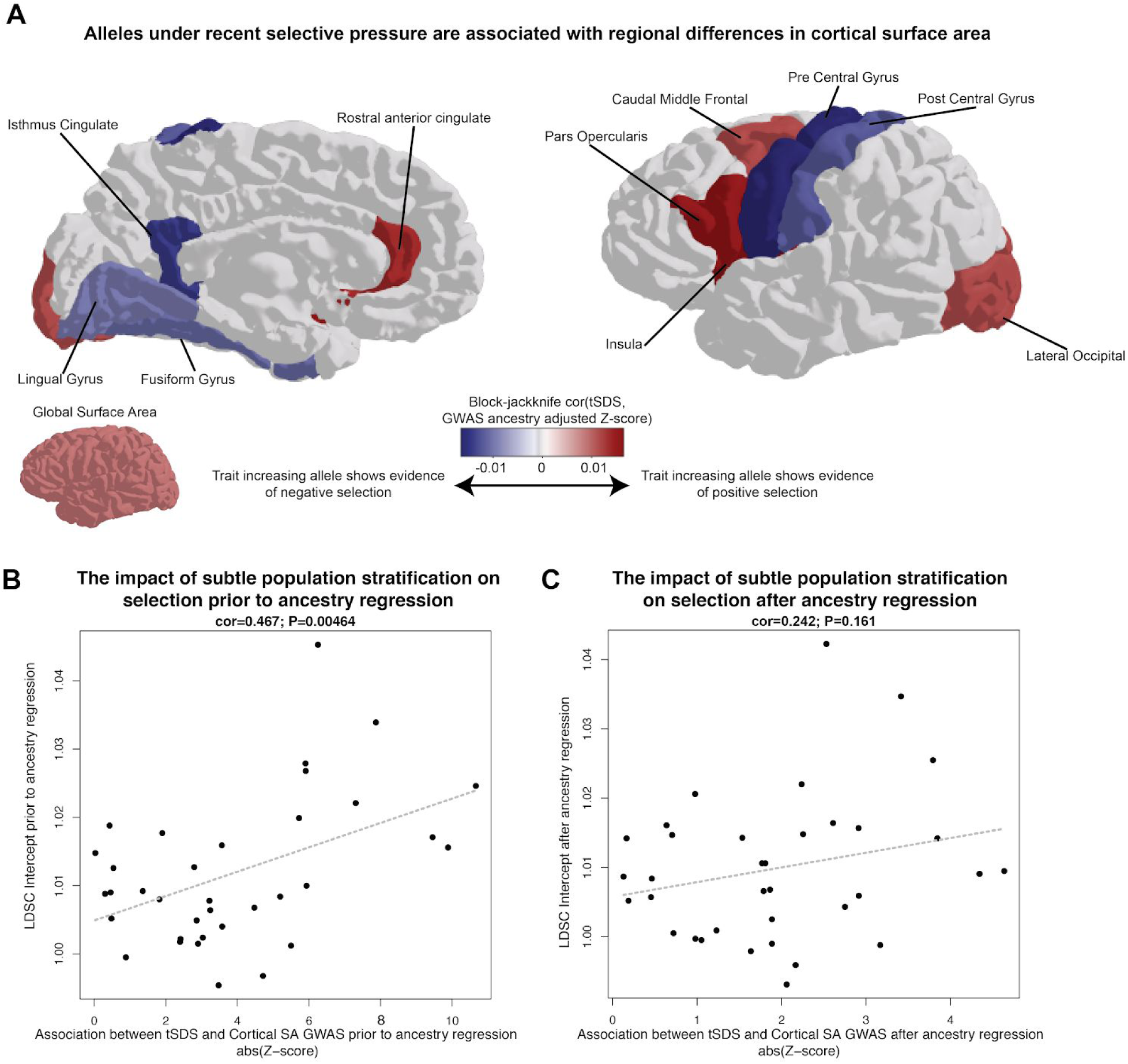
Evidence for haplotypes under very recent selection (∼2,000-3,000 years) impacting cortical structure. (a) A block-jackknife correlation of ancestry regressed effect sizes from GWAS (Z-scores) with scores of recent selection (tSDS) demonstrates evidence for polygenic alleles under selective pressure also influencing both global and regional surface areas (colored regions indicate FDR < 0.05). Colder colors indicate that the trait increasing alleles (associated with increased surface area) are generally associated with negative selection (decreasing allele frequencies in the population), whereas warmer colors indicate that trait increasing alleles are associated with positive selection. **(b)** Subtle population stratification, measured via LD-score regression (LDSC) intercept, is associated with stronger evidence of selection prior to ancestry regression. **(c)** Ancestry regression reduces the relationship between measures of selection and population stratification. See also Figure S6.

We conducted secondary analyses to investigate potential impacts of subtle population stratification because it was recently shown that SDS correlations can be highly influenced by this confounder^35, 36^. Exploratory analyses of GWAS data that were uncorrected for ancestry showed a clear relationship between LD-score regression intercept (a measure of the degree of population stratification) and the level of correlation between SDS and the GWAS Z-scores (cor = 0.467, *p*-value = 0.0046; **Figure 6B**). In contrast, for analyses of GWAS data that had undergone ancestry regression, there was no significant relationship between SDS and GWAS Z-scores (cor = 0.242, *p*-value = 0.161; **Figure 6C**), indicating that the ancestry regression procedure is effective for diminishing confounding effects of population stratification. We note that although the ancestry regression procedure attenuates signals of polygenic selection impacting SA, nevertheless several regions are robust to such adjustment (FDR < 0.05; **Figure 6A**). Finally, we show that SDS correlations within the UKBB European population alone, which is less susceptible to the impacts of subtle population stratification than meta-analysis of data from consortia, shows a highly consistent SDS relationship with the ancestry corrected meta-analysis results (cor = 0.588, *p*-value = 0.00020; **Figure S6**).

## Discussion

By integrating genomic annotations of primate evolutionary history with the largest available genome-wide association analysis of neuroanatomy in living populations^16^, we are able to map genetic variation shaping cortical SA across multiple time periods on the lineage that led to modern humans. Our analyses reveal that common variation found within human-gained enhancers that are active during fetal development has effects on cortical SA measured largely in adults. These findings implicate neural progenitor proliferation and differentiation as processes critical to evolutionary expansion of cortical SA on the human lineage. Such a relationship is consistent with the radial unit hypothesis^5^, which posits that cortical expansion is driven by an increase in the progenitor pool present during development. Moreover, we show that polygenic selection has influenced global SA over the past 2,000 years. Multiple lines of evidence in our study point to effects of polygenic selection on SA of parts of the inferior frontal gyrus, highlighting cortical regions known to be important for the production of spoken language. These results are interesting in light of a recent study that used paleoanthropology, speech biomechanics, ethnography, and historical linguistics to show that changes in human bite configuration and speech-sound inventories occurred after the Neolithic period, potentially due to advances in food-processing technologies^50^. Thus, it is plausible that the consequent increases in the diversity of sounds produced may have led to a subtle, but consistent, selection of alleles increasing cortical SA in brain regions with relevance for speech. If this hypothesis is confirmed, it would represent a novel example of gene-culture co-evolution on the human lineage^51^.

In addition, we identify specific genetic loci that have been subject to selection on our lineage and that impact human cortical SA globally across the brain and within gyrally-defined regions. These include human gained enhancers that map near the developmental transcription factor *HEY2*, and that are associated with interindividual variation in inferior frontal SA. Intriguingly, a rare single gene duplication of *HEY2* was identified in a child with cardiac and neurodevelopmental deficits, including disrupted speech development^52^. Such findings are consistent with common variant effects on the promoter region impacting cortical SA of inferior frontal regions, as these brain areas are important for speech and language functions.

Our study should be carefully interpreted in light of some limitations. First, in this study we were only able to assess a subset of genetic variation that is important for human cortical SA expansion and refinement through human evolution. Specifically, we assess alleles that are both common and polymorphic in current human populations, with a bias towards European ancestry. It is highly likely that derived alleles that are now fixed in modern human populations (and therefore not detectable in GWAS) also made substantial contributions to the shaping of cortical SA during hominid evolution. So far, relatively few of these variants are known^13^, but future studies, for example introducing fixed chimpanzee or Neanderthal alleles into human neural progenitor cells, will help to assess the impacts of this class of genetic variants^53^. Second, subtle population stratification can strongly influence inferences of polygenic selection impacting a trait^35, 36^. Prior to analyses of GWAS data, we implemented an ancestry regression procedure to correct for subtle population stratification^40^. We show that this procedure reduces the impact of population stratification, as evaluated by at least two independent methods (ancestry PC correlations and LDSC intercept). However, allele frequency differences across populations may not be independent of selective pressures, so our procedure may over-correct leading to diminished evidence of selective effects. Even using our conservative ancestry regression approach, robust signals of polygenic selection were detected for cortical surface area, giving us confidence in the results. Nevertheless, replication of our findings in future genetic association studies of brain structure in sufficiently large family-based populations that are less susceptible to impacts of population stratification^54, 55^ would allow further verification of the results presented here. Finally, future studies focusing on understanding genetic influences on behavioral and cognitive traits (language, motor skills)^56^ combined with GWAS of their neurobiological substrates (like this one) may provide a more complete picture of how shifts in genetic variation across time might yield changes in brain structure and behavior.

These findings provide new insights into a number of long-standing debates about the genetic basis for brain size and cortical SA expansion in modern humans. First, consistent with the idea that non-coding genetic variation is a large driver of human brain evolution^57^, we note that genomic annotations of evolutionary history in which cortical SA heritability enrichment was observed are not derived from protein-coding variations. Instead, these come largely from non-coding intergenic or specifically regulatory sequences. Second, our work refutes prior claims that an evolutionary change in just one gene (or perhaps a small handful of genes) can fully account for the distinctive nature of the modern human brain. For example, it was previously proposed that a single genetic variant of strong effect was sufficient to cause the expansion of human brains and cognitive abilities around 50 kya^58^. Here, we not only show that variation in multiple human-gained enhancers influences cortical SA in aggregate, but also find evidence of much more recent polygenic selection acting on these traits. Thus, multiple alleles each of small effect have contributed to the shaping of modern human cortical SA across different evolutionary timescales, even within the last 2,000 years, supporting the importance of gene-culture co-evolution in explaining our biology. In sum, selective pressures over the last 30 million years of human evolution appear to have shaped different aspects of modern human brain structure, from ancient effects on broad growth patterns through to very recent influences on a number of cortical regions, including those linked to our capacity for spoken language.

## Methods

### Genome-wide Association Summary Statistics

Summary statistics for 35 brain phenotypes (global SA and average bilateral SA for 34 regions) were obtained from a European ancestry discovery sample of the ENIGMA cortical SA meta-analysis^16^ including data from the UK Biobank (UKBB). These data correspond to the Biorxiv paper^16^ posted on Sept 9, 2018. Although GWAS of cortical thickness was also calculated in the ENIGMA GWAS, here we focused on surface area given its particular relevance to hominid evolution. Details of image segmentation, genotyping, imputation, association, and meta-analysis are found in the primary GWAS meta-analysis reference^16^. Briefly, magnetic resonance images (MRIs) of the brain were segmented with FreeSurfer^60^ using a gyrally-defined atlas^30^, and visually quality checked based on guidelines provided at the ENIGMA website (http://enigma.ini.usc.edu/research/gwasma-of-cortical-measures/). Imputation of genome-wide genotyping arrays was conducted to the 1000 Genomes phase 1 release v3 reference panel. When conducting associations of gyrally-defined regions, the global measure of SA was included as a covariate, in order to test for genetic influences that were specific to each region. The association models also included four multi-dimensional scaling (MDS) components to help control for ancestry, as well as linear and non-linear corrections for age and sex, diagnostic status and scanner. Fixed effects meta-analysis was used to combine across all sites contributing to the analysis.

In **Figure S6**, GWAS summary statistics were used from the UKBB EUR population only (N = 9,923), which is less susceptible to the impact of population stratification by combining across many sites as in the full ENIGMA analysis.

### Ancestry Regression

We first determined the impact of subtle population stratification on the 35 GWAS summary statistic datasets, in light of studies showing that such stratification can confound estimates of selection^35, 36^. First, all unrelated subjects (defined in^61^) were selected from 1000 Genomes Phase 3 data^62^. We then selected SNPs that had a minor allele frequency (MAF) > 5% in 1000 genomes, and that were not located in the MHC locus, the chromosome 8 inversion region, or regions of long LD. LD independent SNPs (r^2^ < 0.2) were selected via pruning using a window size 500 kb and slide of 100 kb (PLINK –indep-pairwise 500 100 0.2). PCA was performed in PLINK^63^ on the 264,339 remaining SNPs. In order to obtain SNP PC loadings for all SNPs in the 1000 genomes project (MAF < 0.05, MHC locus, the chromosome 8 inversion region, or regions of long LD removed), we performed linear regressions of the PC scores on the genotype allele count of each SNP (after controlling for sex) and used the resulting regression coefficients as the SNP PC loading estimates. This procedure followed that used in previous work^35^. For the first 20 PCs, the weighting of the PCs for each subject was used as a trait and tested for association with each subject’s genotype in PLINK. For each SNP, across all 20 PCs, we identified the degree of association of that SNP to population frequency differences along that principal axis of variation (Beta_PCs). After merging summary statistics of each SA GWAS without genomic control^64^ correction (Beta_strat) with Beta_PC values, flipping beta values to the same effect allele, and sorting based on chromosomal position, a block jackknife correlation with 1000 blocks approach was used to assess the correlation between Beta_strat and Beta_PCs, shown in **Figure 1A** and **Figure S1**.

We then implemented an ancestry regression procedure following previous work^40^. We used a regression model fitting each set of SA GWAS summary statistics without genomic control correction (Beta_strat) simultaneously to the 20 Beta_PC values calculated as described above using the lm() function in R (v3.2.3). The residuals of this model (Beta_r) were used as ancestry corrected effect sizes. Ancestry corrected standard errors and *p*-values were calculated following the same prior work^40^. The same block jackknife correlation method was used to assess the impact of subtle population stratification by correlating Beta_r with Beta_PC in **Figure 1B** and **Figure S2**.

We evaluated an additional measure of population stratification, the LD-score (LDSC) regression intercept^39^, before and after ancestry regression. The summary statistics (with or without ancestry correction, as above) were first written into a standard format using munge_sumstats.py. Then, pre-computed LD scores from 1000 Genomes Phase 3 (using only HapMap3 SNPs, excluding the MHC region) were downloaded from the LDSC website (https://github.com/bulik/ldsc) and implemented according to the guidelines given there.

### Partitioned Heritability

The contributions of each SNP set to the total SNP heritability of each trait were determined using partitioned heritability analyses as implemented in the LDSC software package^41^. The SNPs within HARs^65^, selective sweep regions^26^, Neanderthal introgressed SNPs^24^, and Neanderthal depleted regions^66^ were compared to a control set containing all 1000 Genomes Phase 3 SNPs. Fetal brain HGE SNPs^20^ were compared to a control set containing fetal brain active regulatory elements (E081) from the Epigenomics Roadmap resource, while adult brain HGE SNPs^21^ were compared to the Epigenomics Roadmap adult brain active regulatory elements (E073). Active regulatory elements were defined using chromHMM^67^ marks from the 15 state model including all the following annotations: 1_TssA, 2_TssAFlnk, 7_Enh, 6_EnhG.

### Polygenic Scores

Polygenic scores reflecting the aggregate effect of derived alleles within 7 PCW fetal brain HGEs were constructed from ancestry corrected summary statistics. We began by subsetting the summary statistics to those within the HGEs, followed by clumping using PLINK 1.9 (*r^2^* > 0.2). The clumped SNPs were further subset to those with *p*-values below a range of thresholds (0.5 to 0.0005, 0.005 is presented in the main text). For each set of SNPs (e.g., those with *p*-values less than 0.005), the ancestry corrected betas were aligned to the derived alleles from GRCh37, as reported in the Ensembl multi-species alignment (downloaded using the Bioconductor package ‘biomaRt’ version 2.38.0^68, 69^). At each threshold, the derived allele-aligned betas were divided by their standard errors, summed, and the total was divided by the number of SNPs passing the *p*-value threshold.

### Gene Annotations

Gene sets impacted by genetic variation within 7 PCW HGEs were derived separately for (1) global SA or (2) any of the 34 regional SA loci. We first identified all SNPs within 10,000 kb of a nominally significant (*p*-value < 5×10^−8^) GWAS locus with *r^2^* > 0.6 in the 1000G EUR population to the index SNP, using PLINK 1.9. With this extended list of SNPs in LD with the GWAS index SNP, we looked for overlaps with any of 7 PCW HGEs^20^. For those genome-wide significant loci that also overlapped with HGEs, we then recorded known functional impacts on gene expression using adult brain eQTL from the PsychENCODE dataset^45^, downloaded from http://adult.psychencode.org/ selecting the dataset thresholded by the following parameters (FDR < 0.05, expression > 0.1 FPKM in at least 10 samples). Gene biotype annotations (e.g. protein coding) were called using ENSEMBL via biomaRt.

Pathway enrichment was performed for each gene list using the *gost* function from the ‘gprofiler2’ package (version 0.1.3). Electronic GO annotations (evidence code IEA) were excluded, the sources were limited to GO, KEGG, and Reactome pathways, and FDR correction was applied with a significance threshold of 0.05.

### Qx Score Implementation

Allele frequencies in current human populations were downloaded from the Human Genome Diversity Project^70–72^ (http://plab-server.uchicago.edu). Background selection values were downloaded from previous work^73^. Cortical SA GWAS ancestry regressed summary statistics were first intersected to the set of SNPs present in the HGDP dataset and then clumped to a set of LD-independent SNPs (*r*^2^ < 0.2; 1000G Phase 3 EUR population) at a nominal significance threshold (*p*-value < 1×10^−4^) using PLINK(v1.9). The French population, a European population with similar allele frequencies to the cortical SA GWAS was used as the match population. Clumped SNPs with low minor allele frequency (MAF < 0.05) in the French population were removed from the dataset. Qx score functions were downloaded from https://github.com/jjberg2/PolygenicAdaptationCode and the parameters of PolygenicAdaptationFunction() were set to be identical to the example provided in the repository. The Qx *p*-value was recorded for global SA and each of the 34 gyrally defined regions. Benjamini Hochberg FDR correction was used to correct for multiple comparisons across each of the 35 GWASs used.

### Singleton Density Score Implementation

Singleton Density Scores^28^ for each SNP were downloaded from https://datadryad.org/resource/doi:10.5061/dryad.kd58f. Ancestry regressed summary statistics (without any significance thresholding) were merged with SDS scores by rsID and flipped such that the SDS value describes the trait increasing allele (tSDS). The ancestry regressed Z-score was calculated as ancestry regressed beta divided by the ancestry regressed standard error. Merged files were then sorted by chromosomal position and block jackknife spearman’s correlation with 100 blocks was used to determine the relationship between ancestry regressed Z-scores and tSDS values. Benjamini Hochberg FDR correction was used to correct for multiple comparisons across each of the 35 GWASs used. Results were plotted on a representative brain surface using the R/plotly package, where the correlation values were only shown for significant associations after FDR correction (FDR adjusted *p*-value < 0.05; see **Figure 6A**). These analyses were run in an identical manner in two additional ways: (1) without ancestry regression on the full ENIGMA SA GWASs in **Figures 6B**; (2) without ancestry regression in the UKBB dataset subset to only European individuals (**Figure S6**).

### Data Visualization

Genomic loci plots (Figures 5 and S5) were constructed using the R package ‘GViz’, with evolutionary annotation data sourced from the references given in Data Availability. Brain plots (Figures 3, 4, and 6) were made using the ‘plotly’ package. All other plots were made in R using ‘ggplot2’ and related packages, with final assembly and annotation performed using Inkscape.

### Data and Code Availability

Code used to perform analyses is available at https://bitbucket.org/jasonlouisstein/e3enigmaevol_scripts/src/master/. Genomic regions that underwent rapid change on the human lineage (human accelerated regions, HARs) were combined from several sources^74–78^. BED files listing fetal brain enhancer elements not found in macaques or mice were obtained from Reilly, *et al.*, 2015. Adult brain enhancer elements arising since our last common ancestor with the macaque or chimpanzee were obtained from ^21^. A refined list of SNPs gained through introgression with Neanderthals was obtained from Simonti, *et al*., 2016. Genomic regions depleted of introgressed Neanderthal DNA were obtained from Supplemental Table 8 in Vernot, *et al*., 2016. Ancient selective sweep regions identified using extended lineage sorting were obtained from Peyregne, *et al*., 2017.

## ACKNOWLEDGEMENTS

This work was supported in part by the Foundation of Hope, the Brain Research Foundation, the National Institutes of Health (R01 MH118349, R00 MH102357, U54 EB020403), and the National Science Foundation (ACI-16449916). SEF is supported by the Max Planck Society. We thank Leo Zsembik and Shana Hall for initial work on polygenic selection analyses.

## AUTHOR CONTRIBUTIONS

SEF and JLS originated the project and oversaw the work. AKT, SEF, and JLS drafted the manuscript. AKT performed partitioned heritability analyses, polygenic score analyses, and gene ontology enrichment analyses. JLS implemented ancestry regression. SL, SMB, and JLS implemented recent polygenic selection analyses. EAK, BES, LKD provided data for performing ancestry regression and independently implemented analyses. KG, NJ, JP, LCC, JB, DPH, PAL, PMT, SEM, and JLS calculated SA GWAS summary statistics. All authors edited the manuscript.

## FINANCIAL DISCLOSURES

DPH is a full-time employee of Genentech, Inc.

## SUPPLEMENTAL INFORMATION

**Table S1, related to Figure 1:**
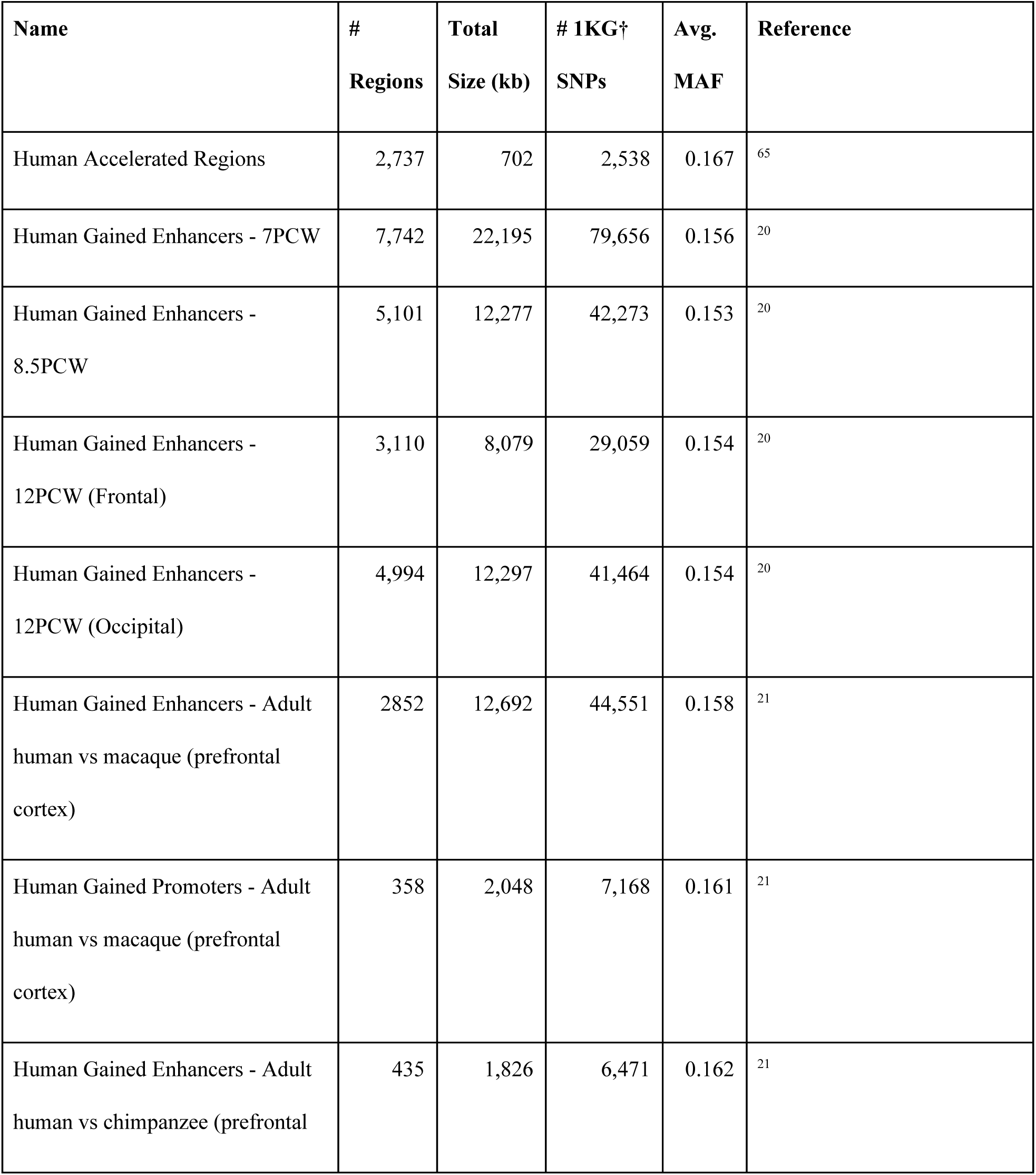
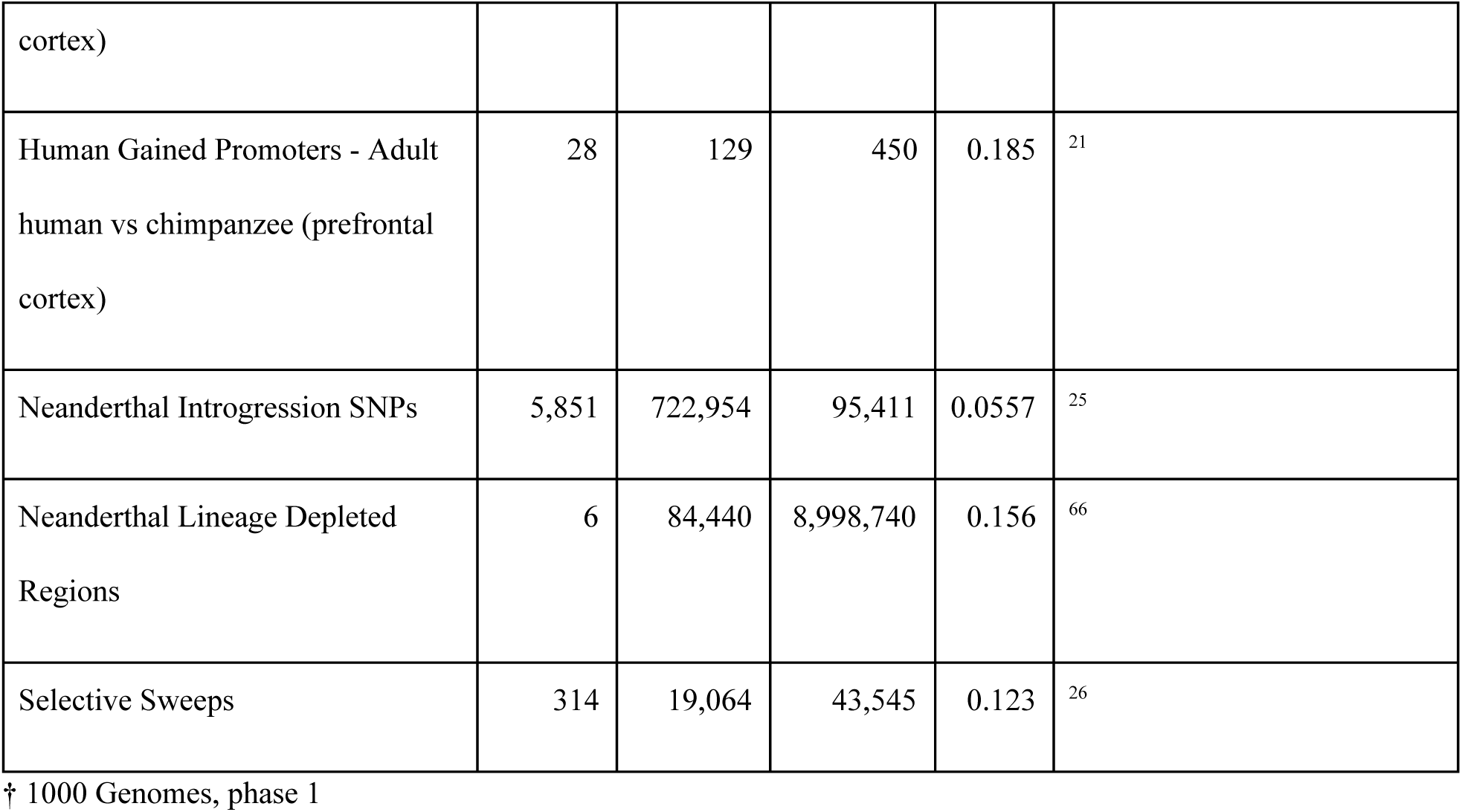
Characteristics of human-evolution focused genome annotations in 1000 Genomes Phase 1 (EUR)

**Table S2, related to Figure 5.** eGenes impacted by loci within HGEs that are also associated with Regional or Global SA.

**Table S3, related to Figure 6.** Qx score results are available as an Excel file.

**Figure S1, related to Figure 2.**
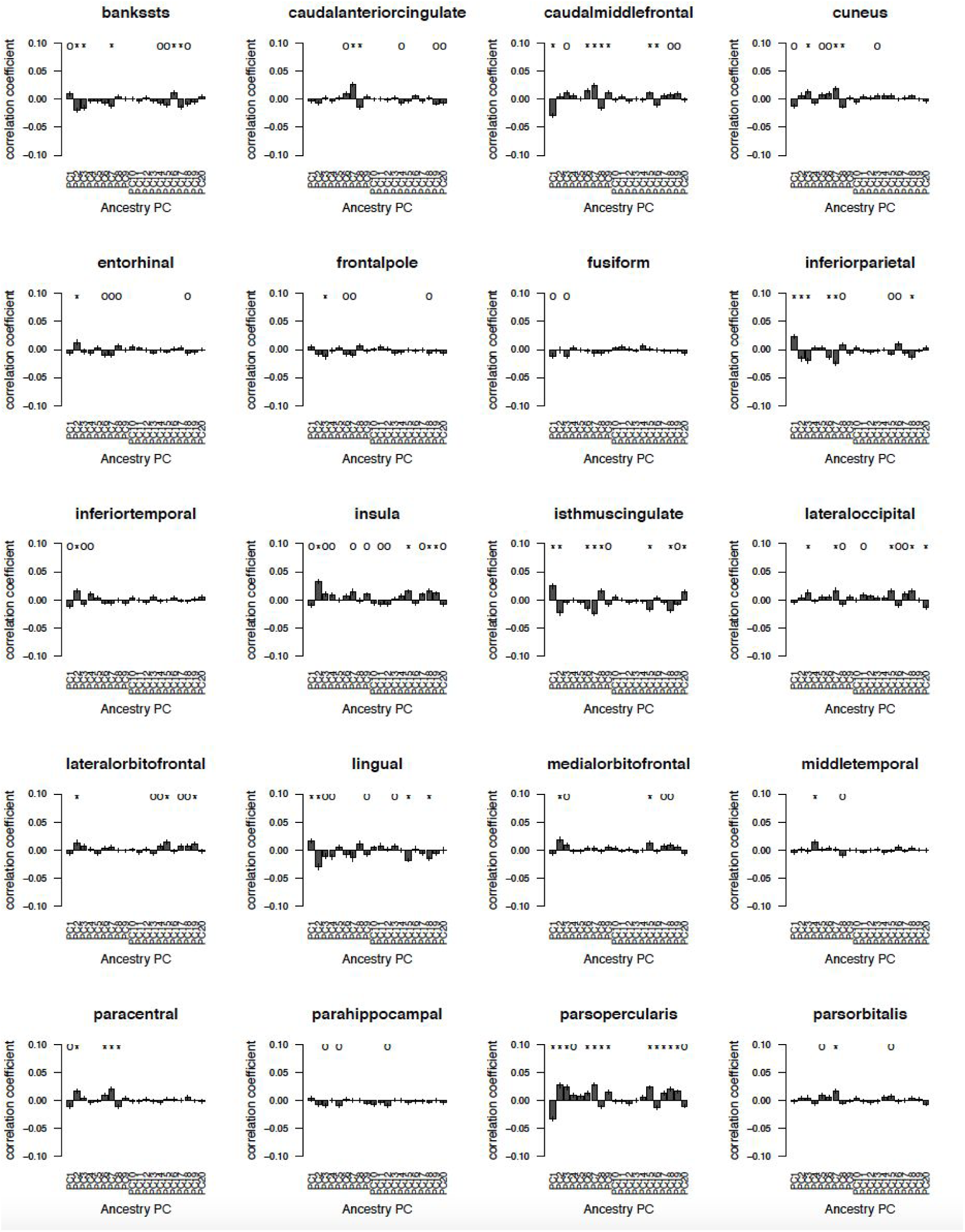
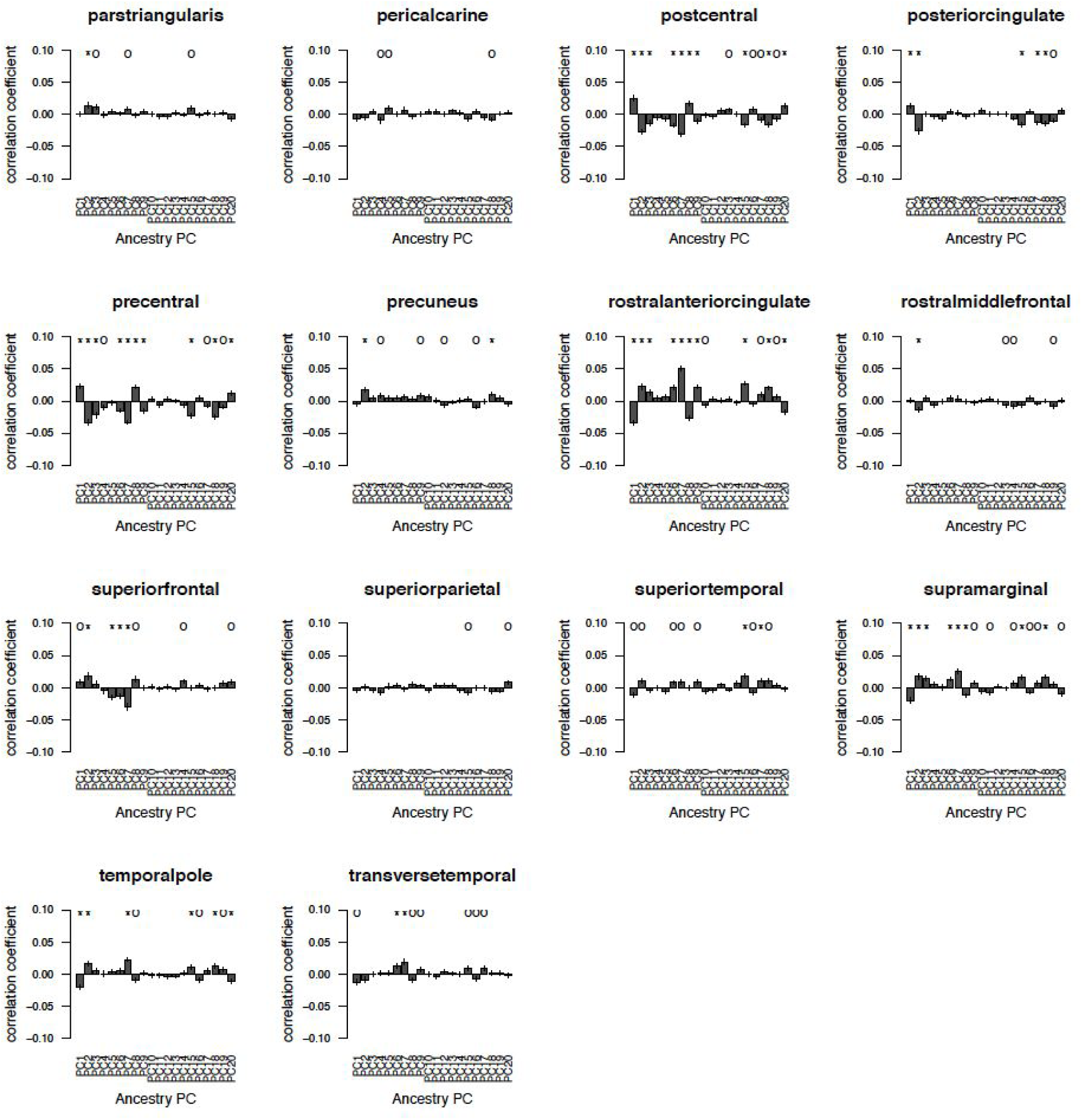
Detecting subtle ancestry regression in cortical SA regional GWASs. Error bars are standard errors.

**Figure S2, related to Figure 2.**
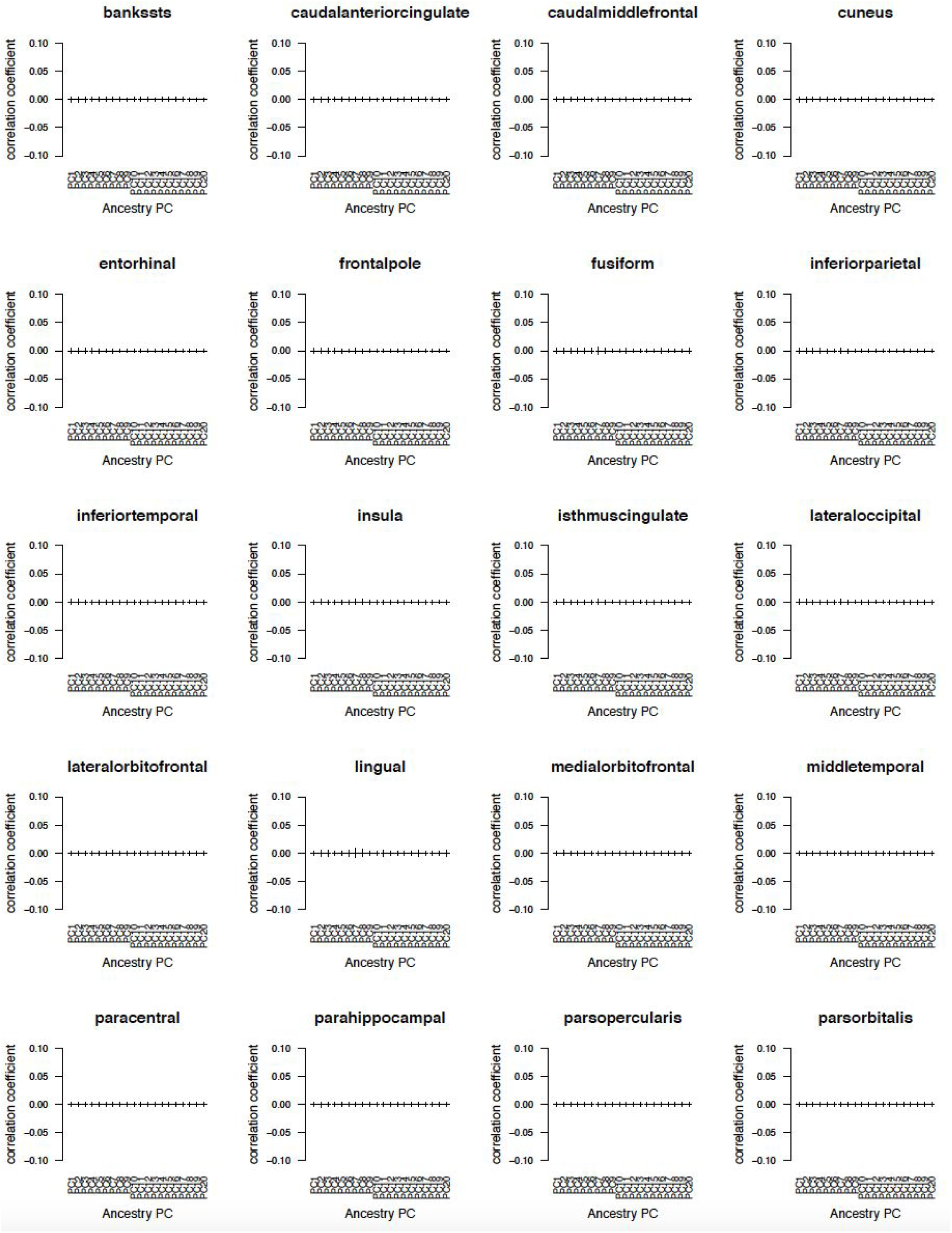
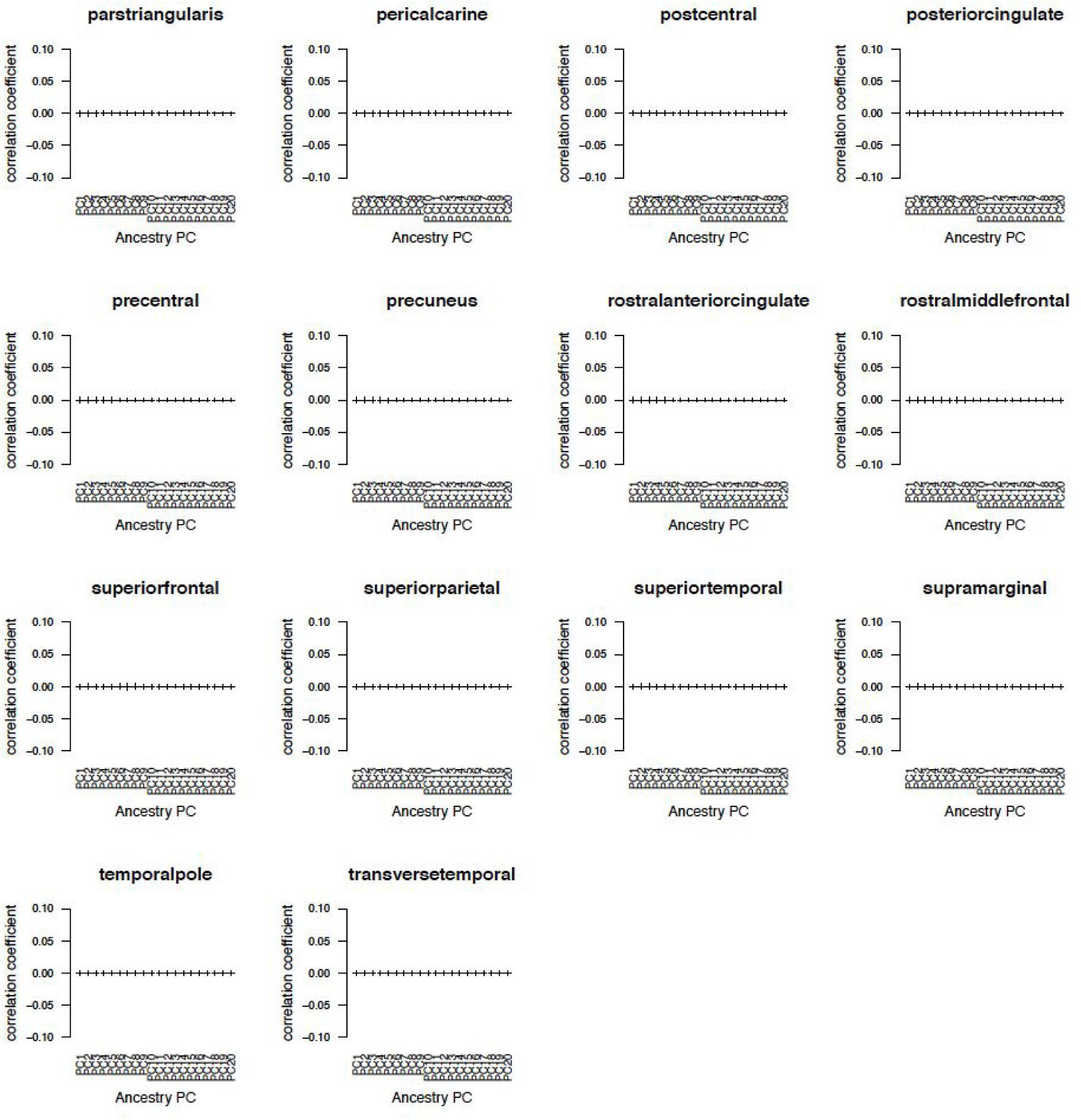
Ancestry regressed cortical SA regional GWASs show diminished effects of subtle population stratification. Error bars are standard errors.

**Figure S3, related to Figure 4.**
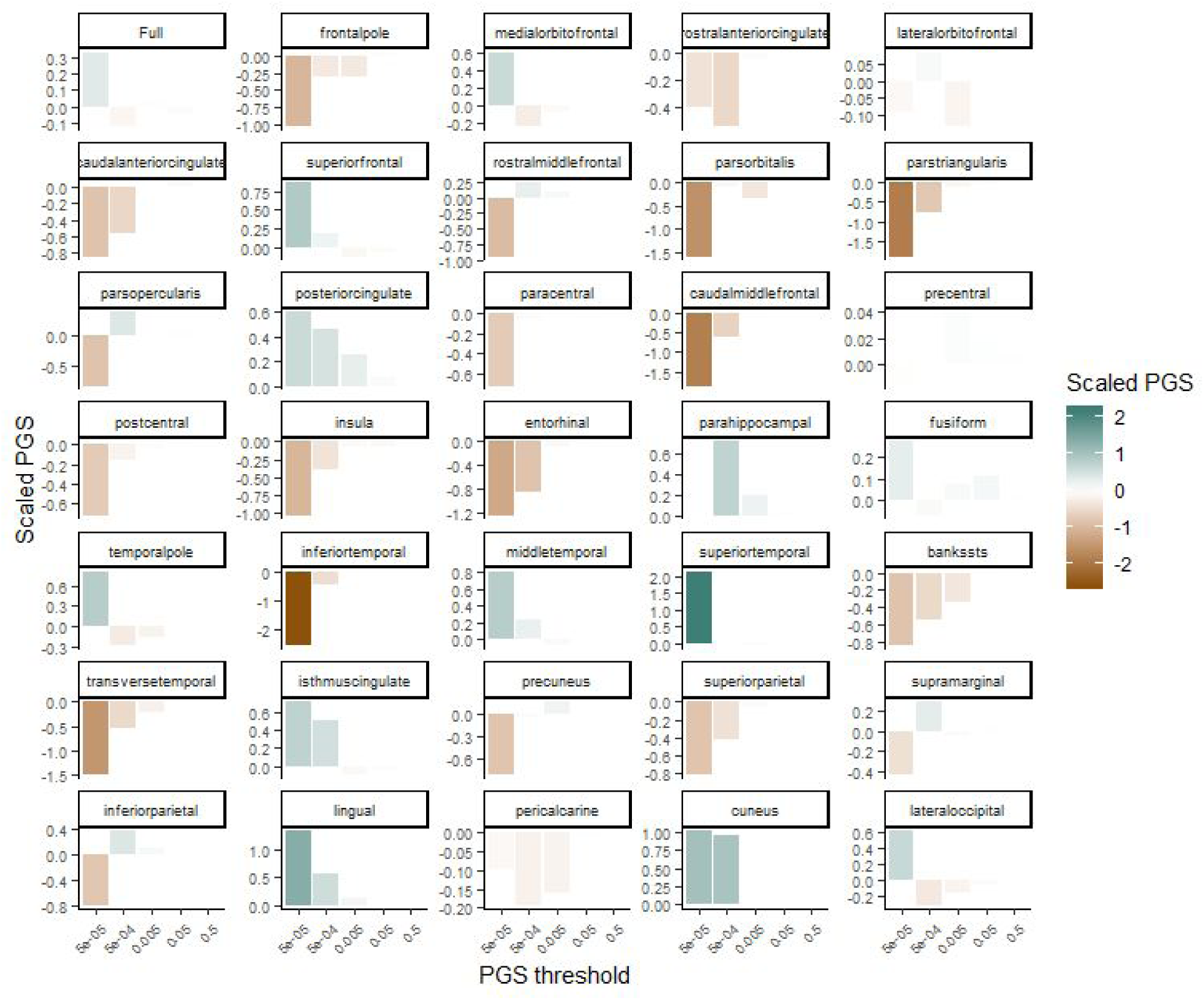
Derived allele-based polygenic scores (PGS) across *p*-value thresholds ranging from 5e-05 to 0.5.

**Figure S4, related to Figure 3.**
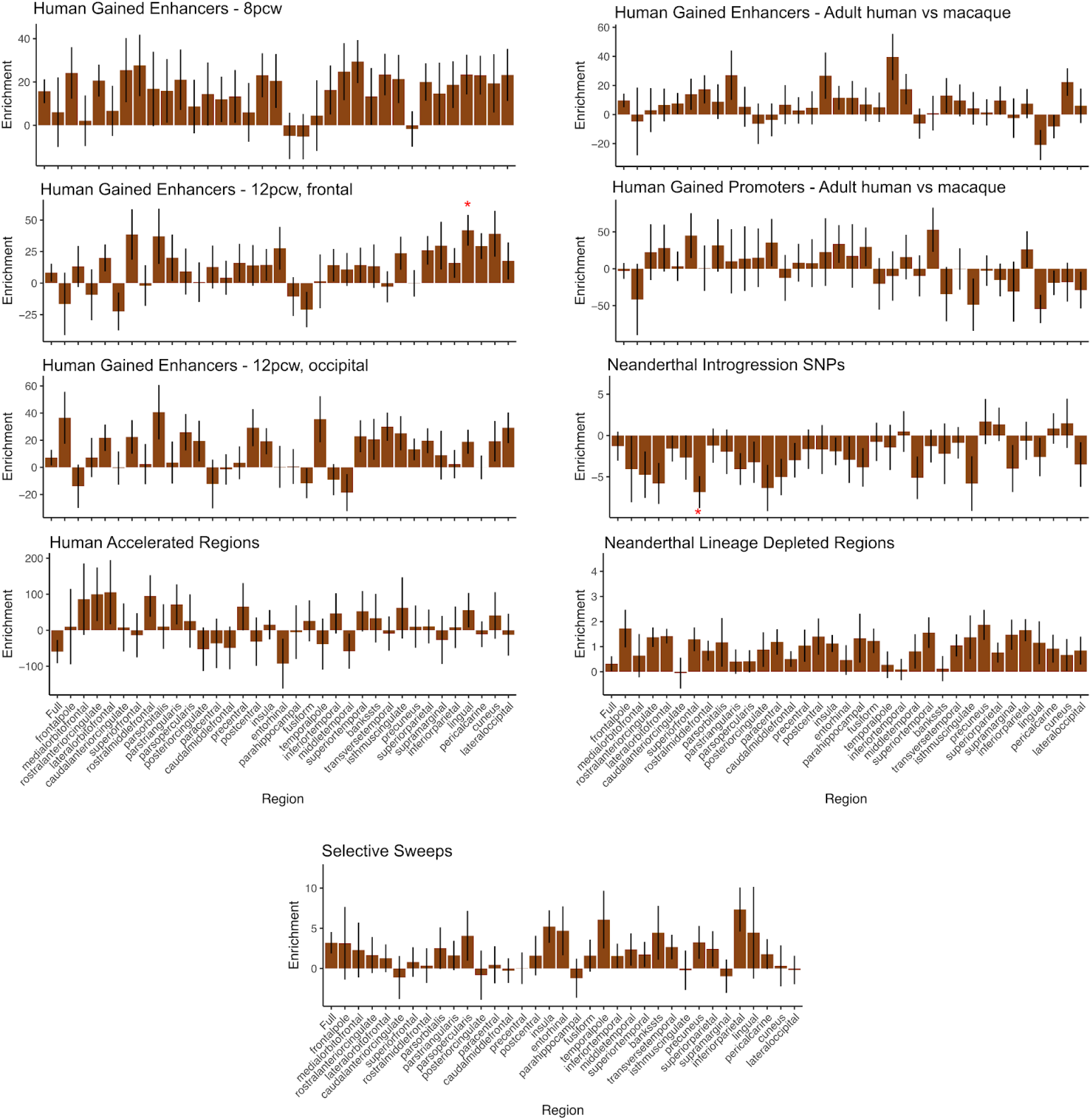
Cortical surface area enrichment scores for Human Accelerated Regions (HARs), fetal and adult HGEs, selective sweeps, Neanderthal introgressed regions, and Neanderthal depleted regions. Error bars represent standard errors, stars indicate FDR corrected p-values less than 0.05.

**Figure S5, related to Figure 5.** GViz plots for 60 GWAS loci are available as a separate PDF.

**Figure S6, related to Figure 6:**
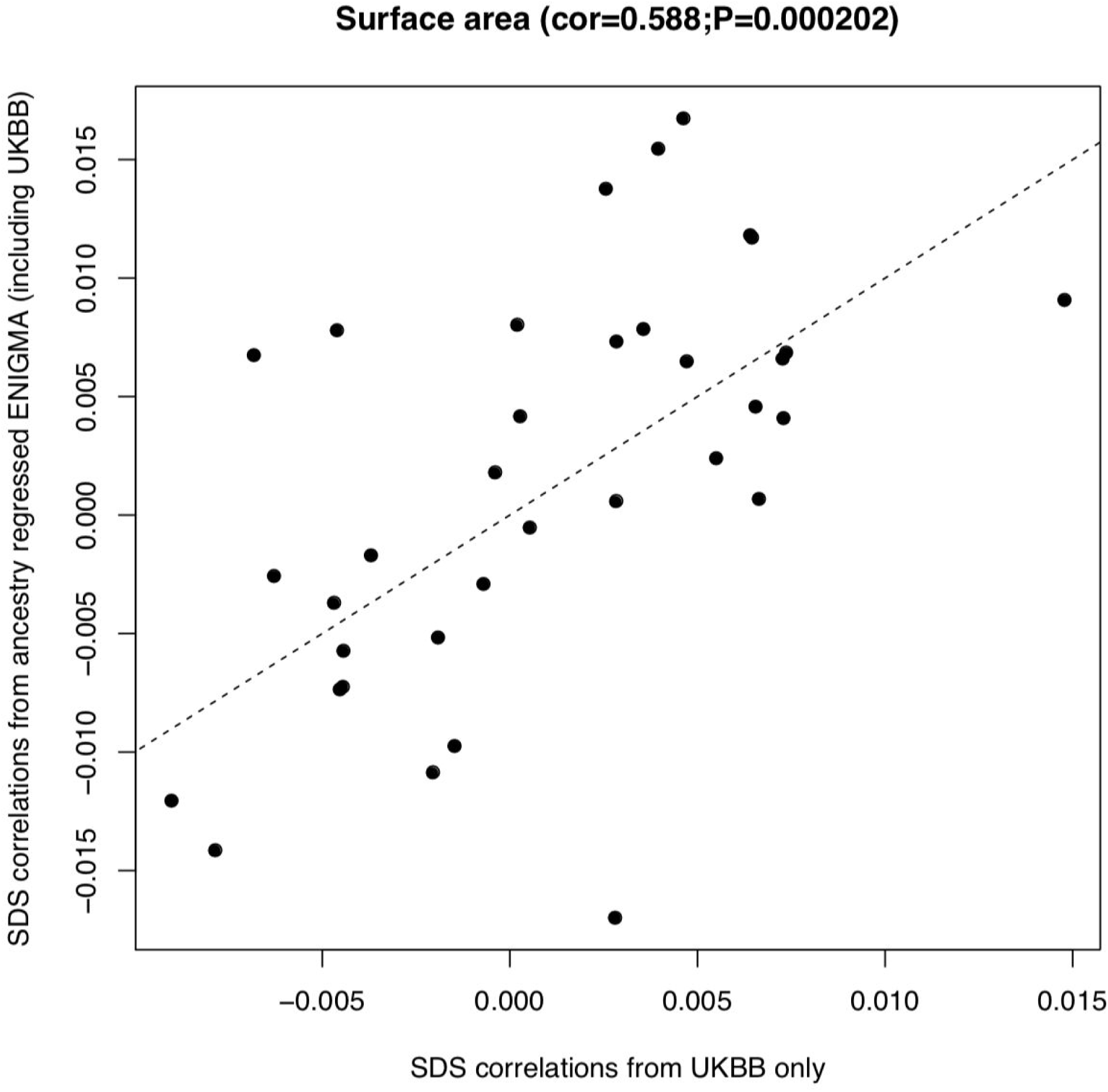
SDS correlations to cortical surface area in a population less susceptible to subtle population stratification (UKBB EUR) and the ancestry regressed ENIGMA data. The y=x line is shown.

