## Supplementary material for "The Evolutionary History of Common Genetic Variants Influencing Human Cortical Surface Area": Figure S5

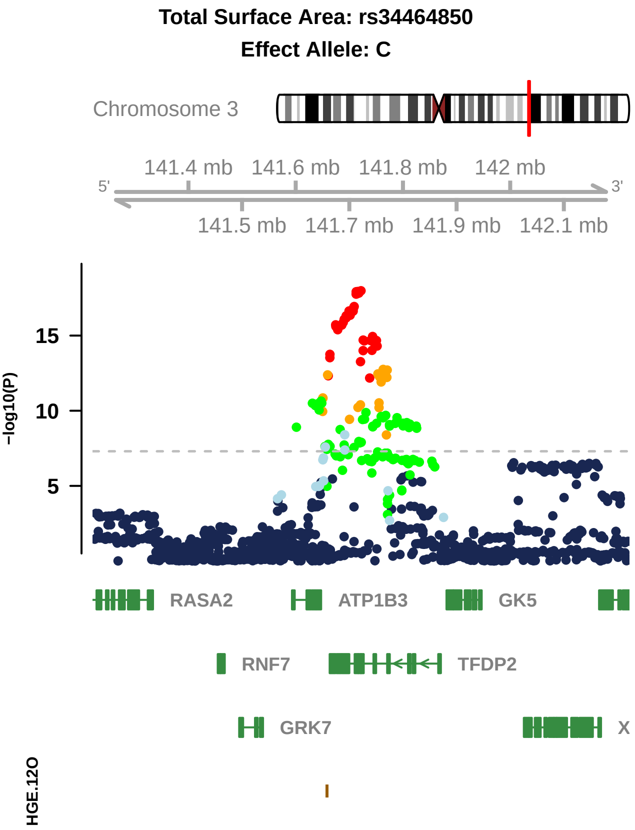

Total Surface Area: rs34464850
Effect Allele: C
Chromosome 3
141.4 mb
141.6 mb
141.8 mb
142 mb
5'
3'
141.5 mb
141.7 mb
141.9 mb
142.1 mb
15
10
5
-log10(P)
RASA2
ATP1B3
GK5
RNF7
TFDP2
GRK7
X
HGE.12O

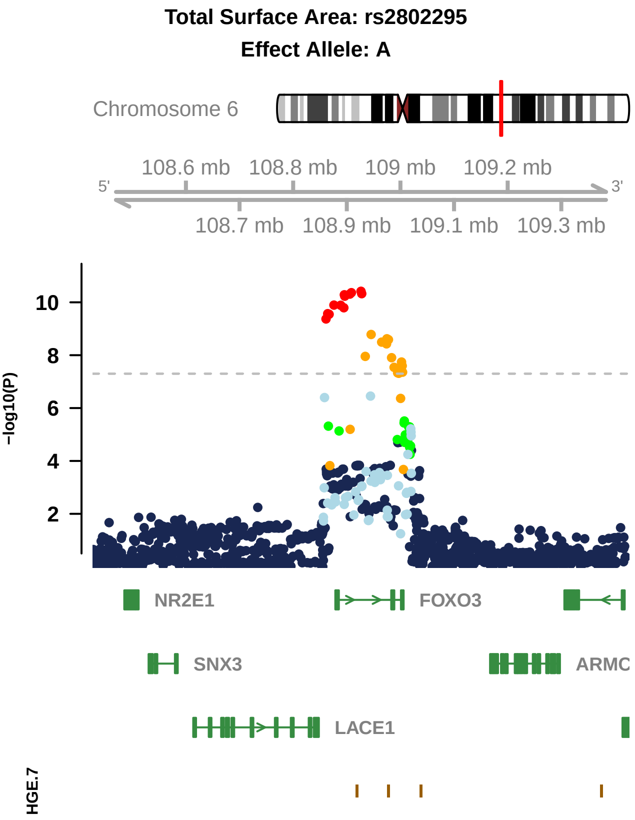
Total Surface Area: rs2802295
Effect Allele: A
Chromosome 6
108.6 mb
108.8 mb
109 mb
109.2 mb
5'
3'
108.7 mb
108.9 mb
109.1 mb
109.3 mb
−log10(P)
10
8
6
4
2
NR2E1
FOXO3
SNX3
ARMC
LACE1
HGE.7

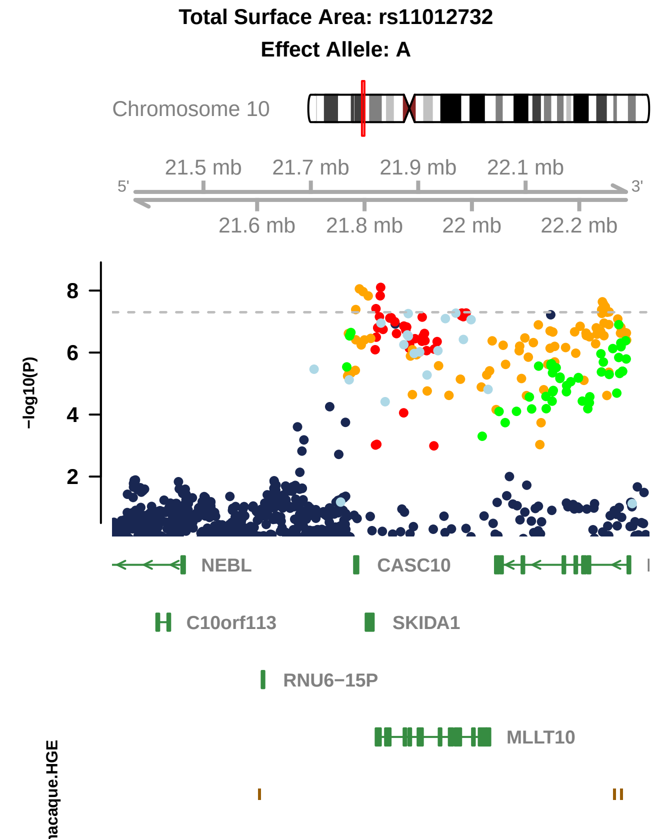

Total Surface Area: rs11012732
Effect Allele: A
Chromosome 10
21.5 mb
21.7 mb
21.9 mb
22.1 mb
5'
3'
21.6 mb
21.8 mb
22 mb
22.2 mb
-log10(P)
8
6
4
2
NEBL
CASC10
C10orf113
SKIDA1
RNU6-15P
MLLT10

### Total Surface Area: rs11171739

### Effect Allele: T

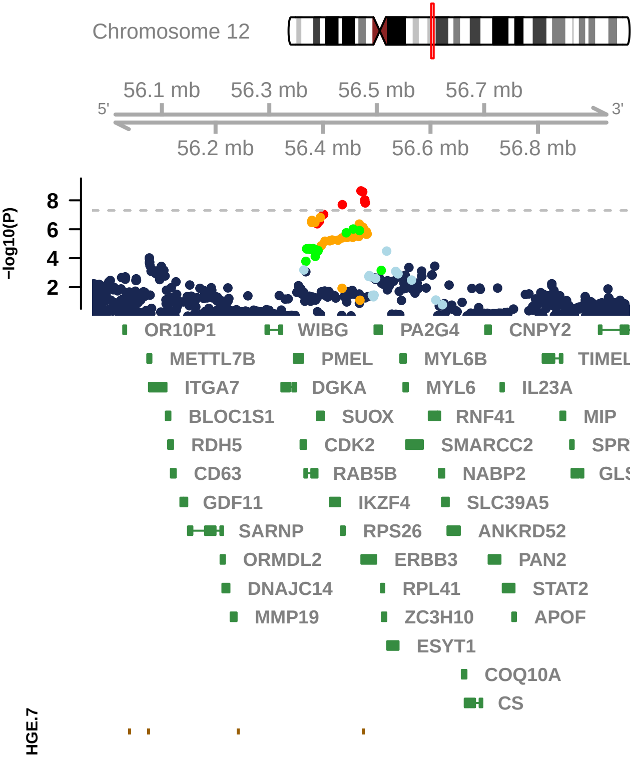

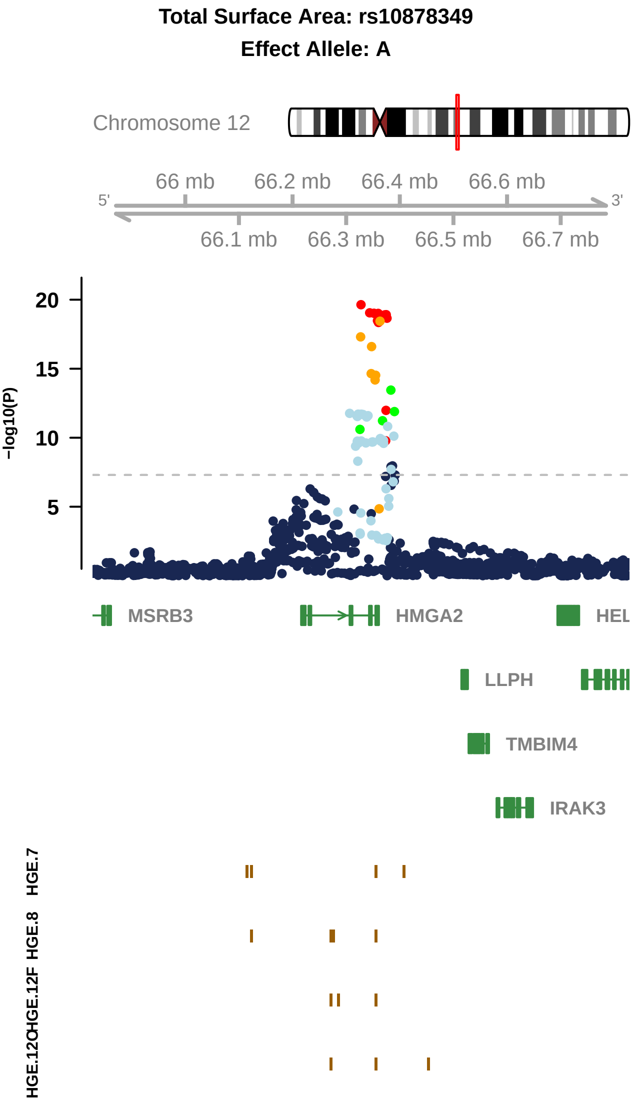

Total Surface Area: rs10878349
Effect Allele: A
Chromosome 12
66 mb
66.2 mb
66.4 mb
66.6 mb
5'
3'
66.1 mb
66.3 mb
66.5 mb
66.7 mb
20
15
10
5
-log10(P)
MSRB3
HMGA2
HEL
LLPH
TMBIM4
IRAK3
HGE.7
HGE.8
HGE.12F
HGE.12O

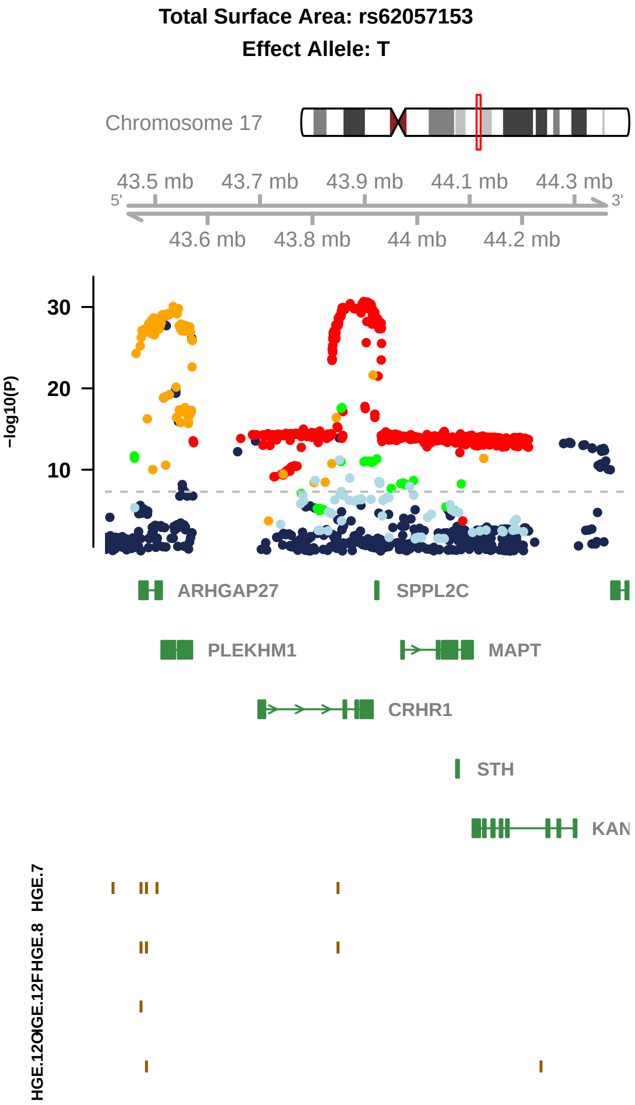

Total Surface Area: rs62057153
Effect Allele: T
Chromosome 17
43.5 mb
43.7 mb
43.9 mb
44.1 mb
44.3 mb
5'
3'
43.6 mb
43.8 mb
44 mb
44.2 mb
30
20
10
-log10(P)
ARHGAP27
SPPL2C
PLEKHM1
MAPT
CRHR1
STH
KAN
HGE.7
HGE.8
HGE.12F
HGE.12OI

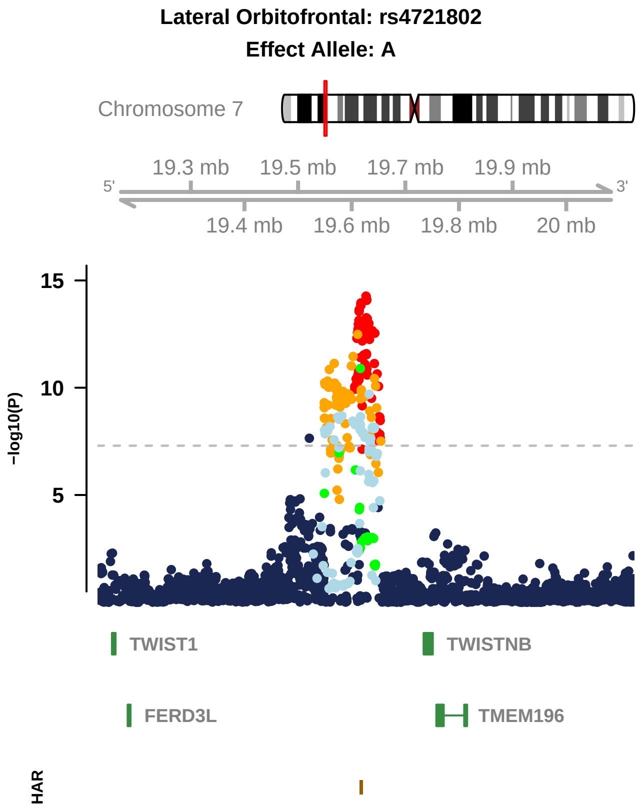

Lateral Orbitofrontal: rs4721802
Effect Allele: A
Chromosome 7
19.3 mb
19.5 mb
19.7 mb
19.9 mb
5'
3'
19.4 mb
19.6 mb
19.8 mb
20 mb
15
10
5
−log10(P)
TWIST1
TWISTNB
FERD3L
TMEM196
HAR

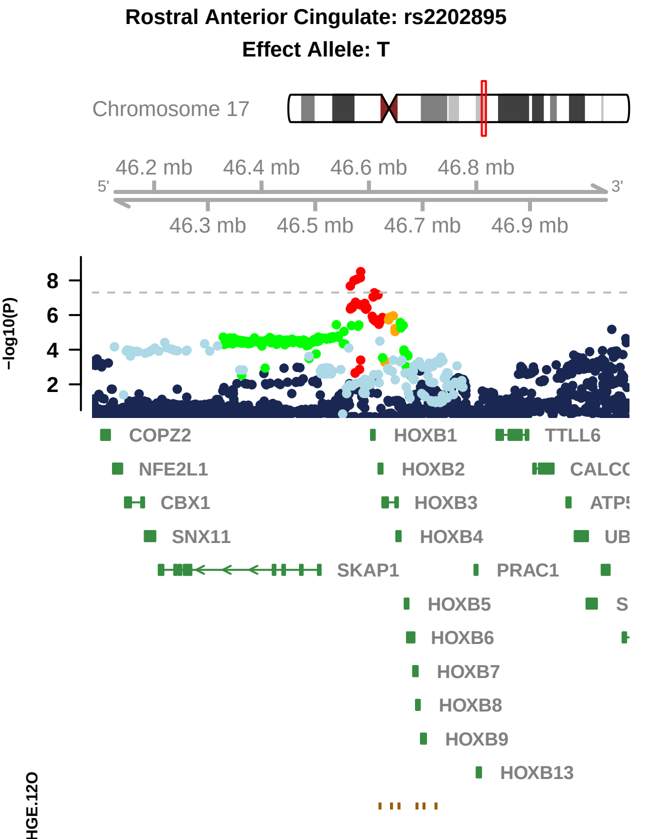

Rostral Anterior Cingulate: rs2202895
Effect Allele: T
Chromosome 17
46.2 mb
46.4 mb
46.6 mb
46.8 mb
5'
3'
46.3 mb
46.5 mb
46.7 mb
46.9 mb
-log10(P)
8
6
4
2
COPZ2
NFE2L1
CBX1
SNX11
SKAP1
HOXB1
HOXB2
HOXB3
HOXB4
PRAC1
HOXB5
HOXB6
HOXB7
HOXB8
HOXB9
HOXB13
TTLL6
HGE.12O

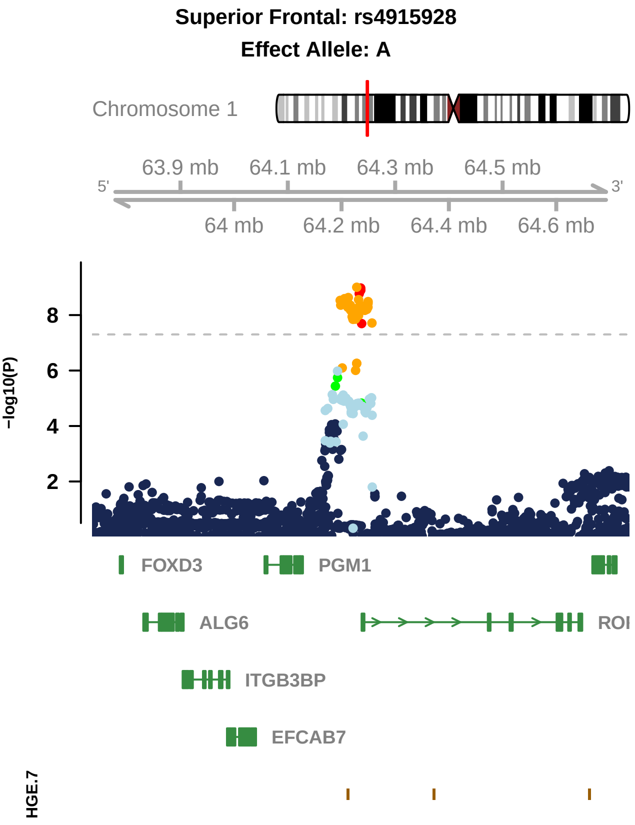

Superior Frontal: rs4915928
Effect Allele: A
Chromosome 1
63.9 mb
64.1 mb
64.3 mb
64.5 mb
5'
3'
64 mb
64.2 mb
64.4 mb
64.6 mb
8
6
4
2
-log10(P)
FOXD3
PGM1
ALG6
ROI
ITGB3BP
EFCAB7
HGE.7

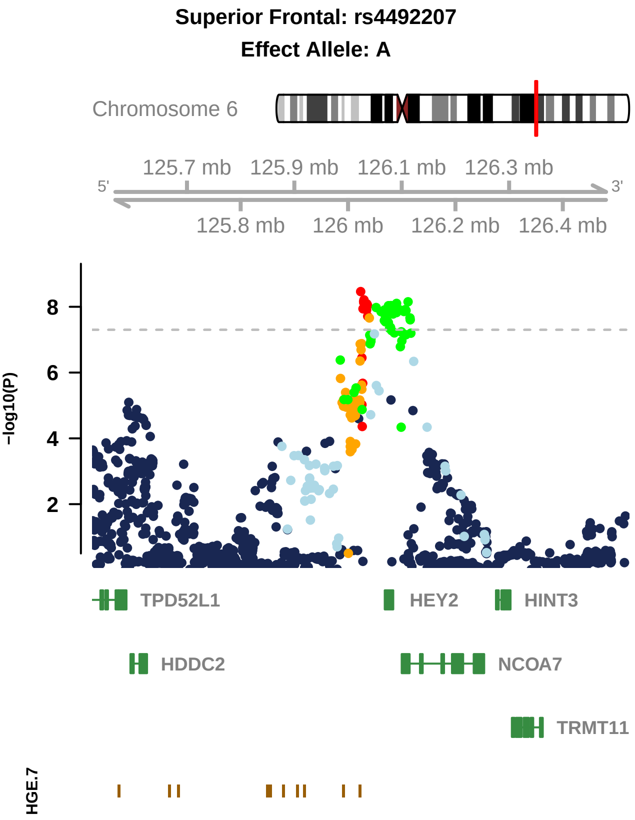
Superior Frontal: rs4492207
Effect Allele: A
Chromosome 6
125.7 mb
125.9 mb
126.1 mb
126.3 mb
5'
3'
125.8 mb
126 mb
126.2 mb
126.4 mb
8
6
4
2
−log10(P)
TPD52L1
HEY2
HINT3
HDDC2
NCOA7
TRMT11
HGE.7

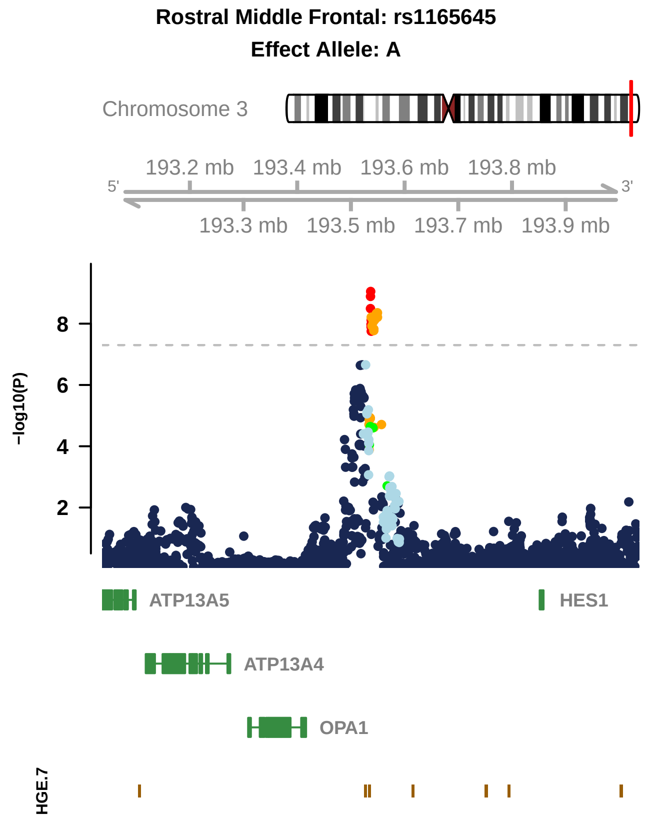

Rostral Middle Frontal: rs1165645
Effect Allele: A
Chromosome 3
193.2 mb
193.4 mb
193.6 mb
193.8 mb
5'
3'
193.3 mb
193.5 mb
193.7 mb
193.9 mb
8
6
4
2
−log10(P)
ATP13A5
HES1
ATP13A4
OPA1
HGE.7

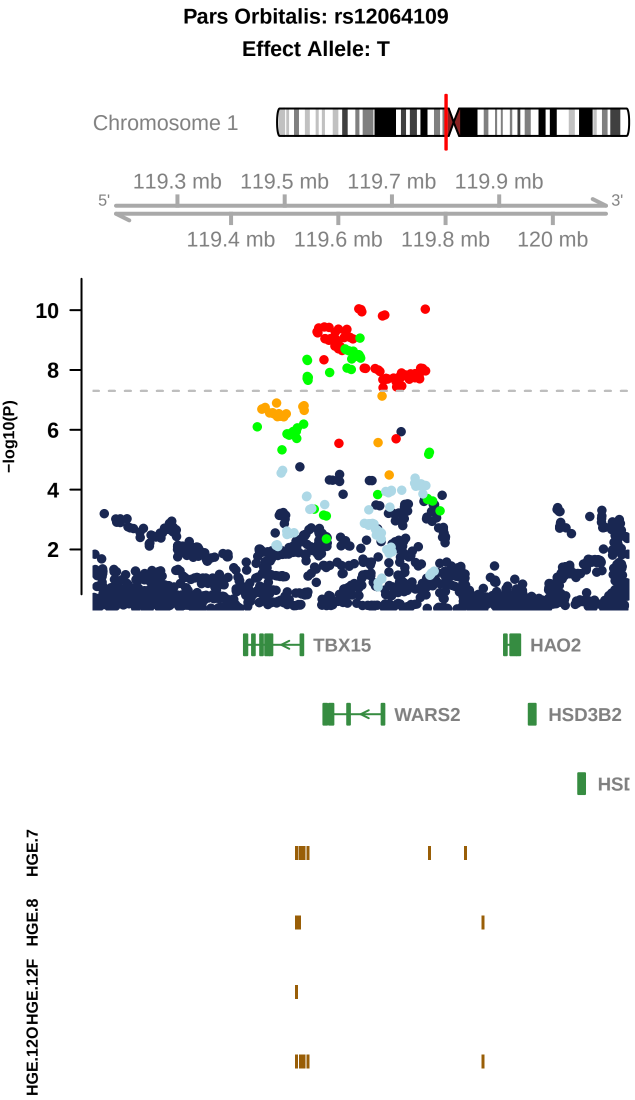

Pars Orbitalis: rs12064109
Effect Allele: T
Chromosome 1
119.3 mb
119.5 mb
119.7 mb
119.9 mb
5'
3'
119.4 mb
119.6 mb
119.8 mb
120 mb
−log10(P)
10
8
6
4
2
TBX15
HAO2
WARS2
HSD3B2
HSD
HGE.7
HGE.8
HGE.12F
HGE.12O

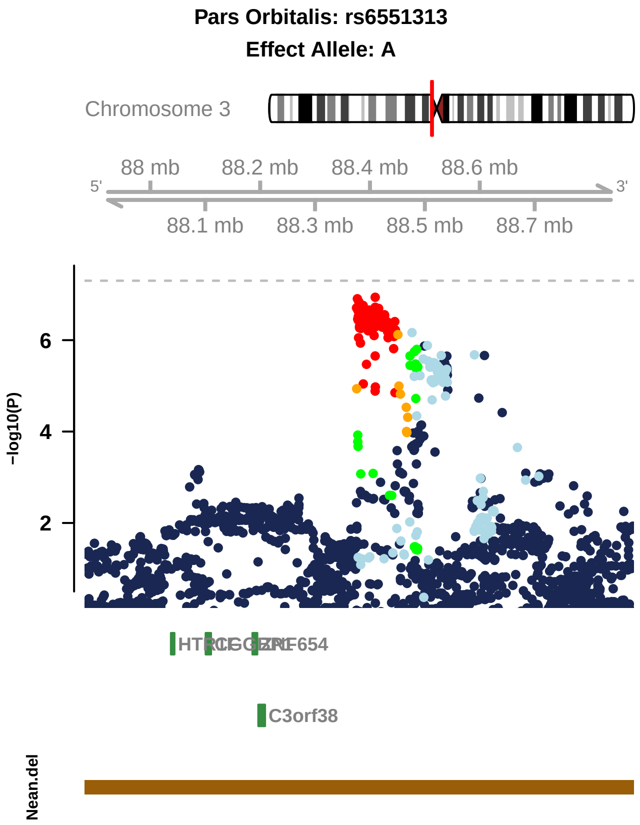

Pars Orbitalis: rs6551313
Effect Allele: A
Chromosome 3
88 mb
88.2 mb
88.4 mb
88.6 mb
5'
3'
88.1 mb
88.3 mb
88.5 mb
88.7 mb
6
4
2
−log10(P)
C3orf38
Nean.del

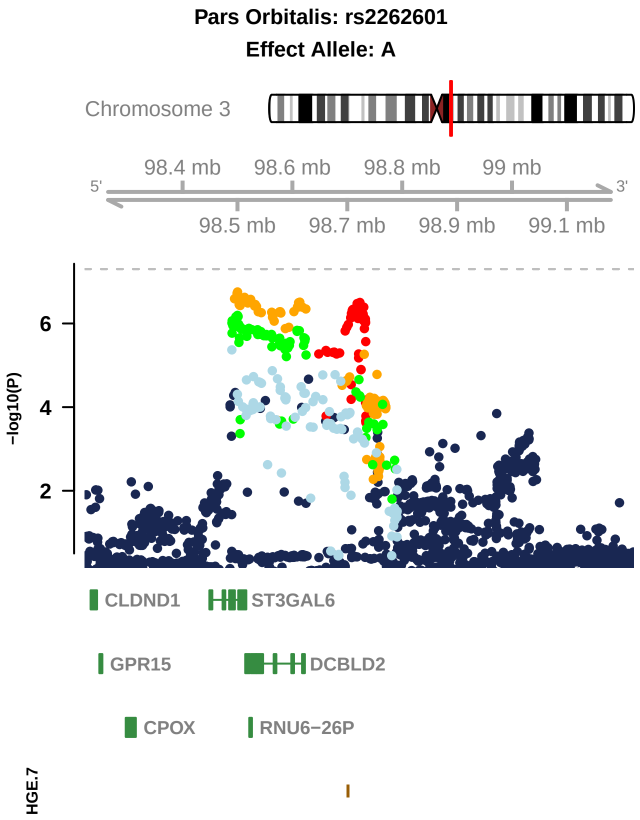

Pars Orbitalis: rs2262601
Effect Allele: A
Chromosome 3
98.4 mb
98.6 mb
98.8 mb
99 mb
5'
3'
98.5 mb
98.7 mb
98.9 mb
99.1 mb
6
4
2
−log10(P)
CLDND1
ST3GAL6
GPR15
DCBLD2
CPOX
RNU6−26P
HGE.7

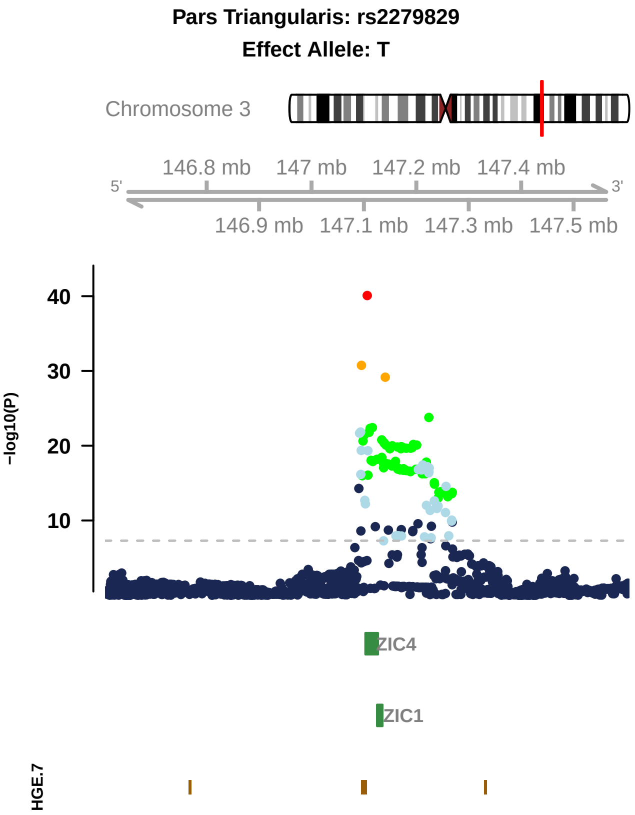

Pars Triangularis: rs2279829
Effect Allele: T
Chromosome 3
146.8 mb
147 mb
147.2 mb
147.4 mb
5'
3'
146.9 mb
147.1 mb
147.3 mb
147.5 mb
40
30
20
10
−log10(P)
ZIC4
ZIC1
HGE.7

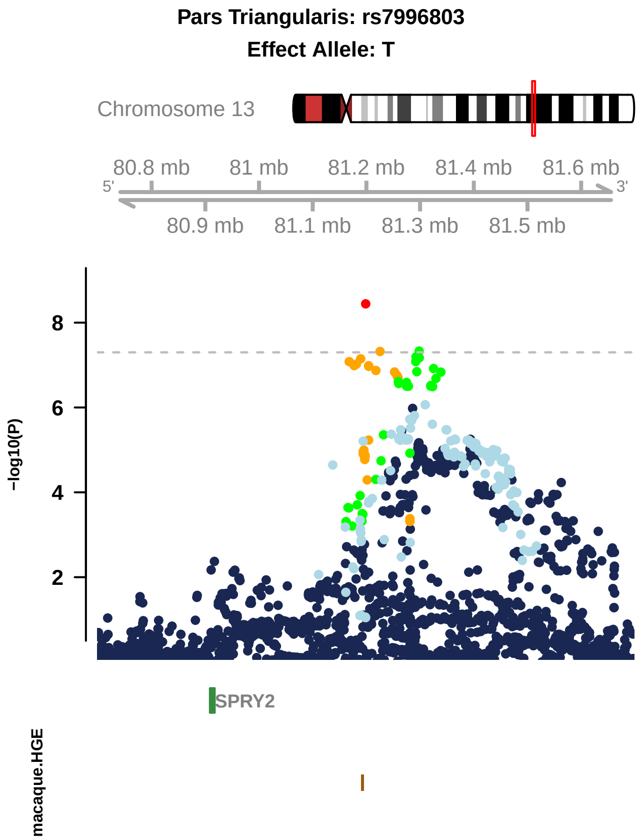

Pars Triangularis: rs7996803
Effect Allele: T
Chromosome 13
80.8 mb
81 mb
81.2 mb
81.4 mb
81.6 mb
5'
3'
80.9 mb
81.1 mb
81.3 mb
81.5 mb
-log10(P)
8
6
4
2
SPRY2
macaque.HGE

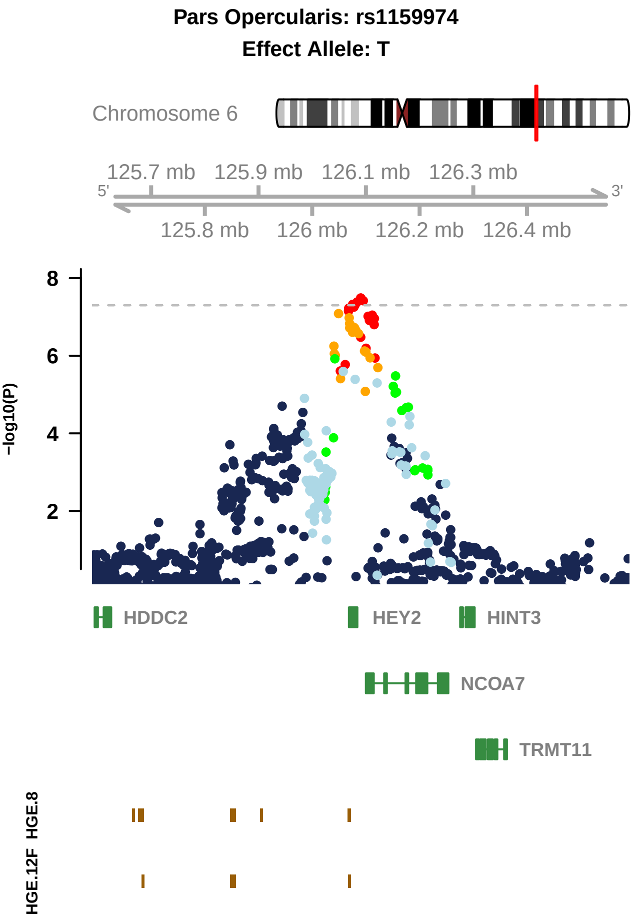

Pars Opercularis: rs1159974
Effect Allele: T
Chromosome 6
125.7 mb
125.9 mb
126.1 mb
126.3 mb
5'
3'
125.8 mb
126 mb
126.2 mb
126.4 mb
8
6
4
2
−log10(P)
HDDC2
HEY2
HINT3
NCOA7
TRMT11
HGE.8
HGE.12F

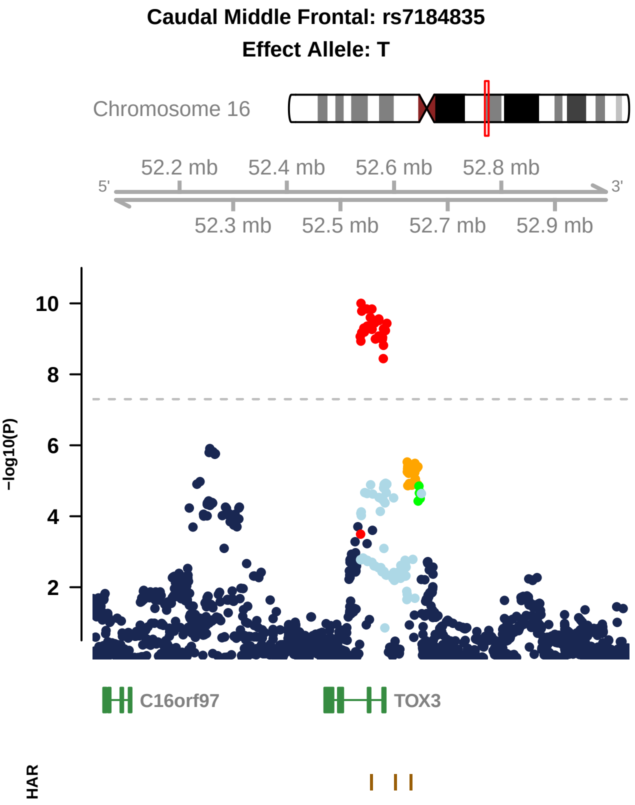

Caudal Middle Frontal: rs7184835
Effect Allele: T
Chromosome 16
52.2 mb
52.4 mb
52.6 mb
52.8 mb
5'
3'
52.3 mb
52.5 mb
52.7 mb
52.9 mb
-log10(P)
10
8
6
4
2
C16orf97
TOX3
HAR

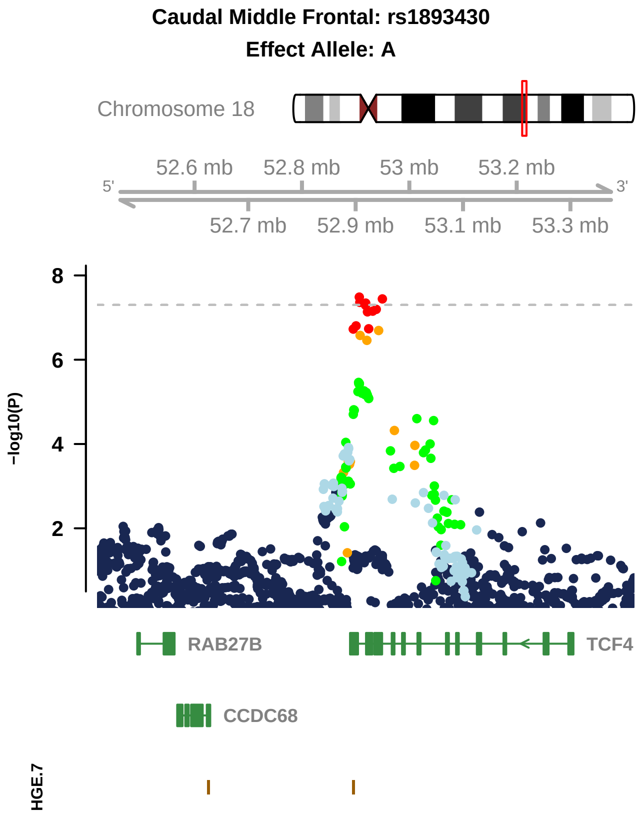
Caudal Middle Frontal: rs1893430
Effect Allele: A
Chromosome 18
52.6 mb
52.8 mb
53 mb
53.2 mb
5'
3'
52.7 mb
52.9 mb
53.1 mb
53.3 mb
8
6
4
2
-log10(P)
RAB27B
TCF4
CCDC68
HGE.7

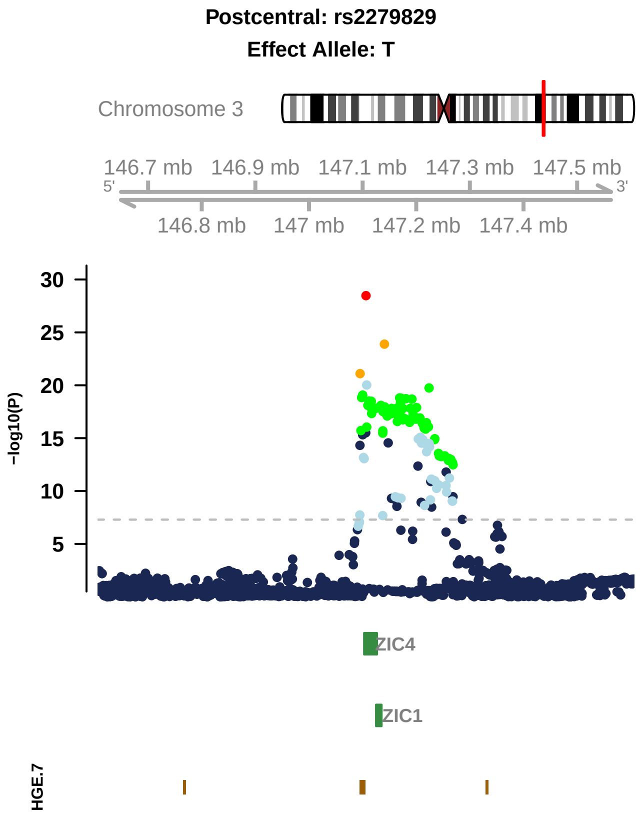

Postcentral: rs2279829
Effect Allele: T
Chromosome 3
146.7 mb
146.9 mb
147.1 mb
147.3 mb
147.5 mb
5'
3'
146.8 mb
147 mb
147.2 mb
147.4 mb
30
25
20
15
10
5
−log10(P)
ZIC4
ZIC1
HGE.7

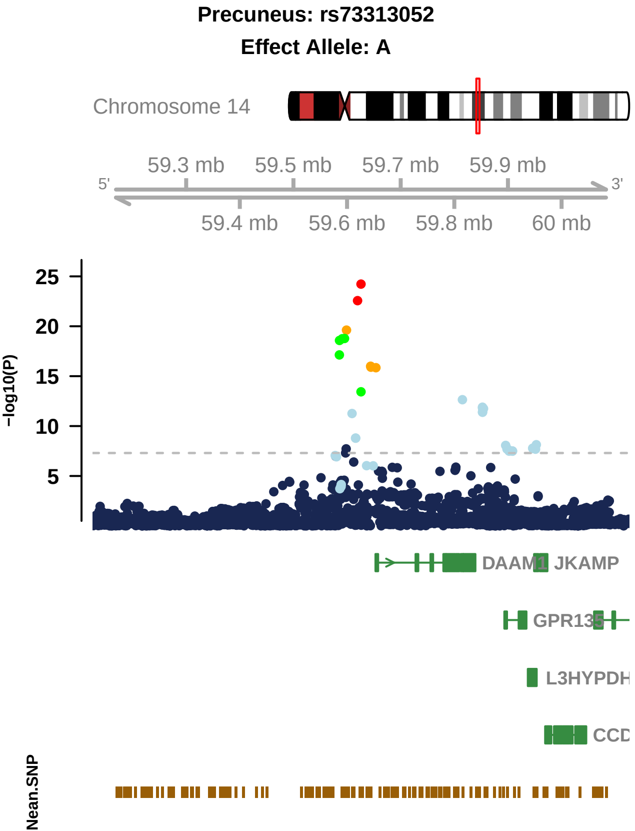

Precuneus: rs73313052
Effect Allele: A
Chromosome 14
59.3 mb
59.5 mb
59.7 mb
59.9 mb
5'
3'
59.4 mb
59.6 mb
59.8 mb
60 mb
−log10(P)
25
20
15
10
5
DAAM1
JKAMP
GPR135
L3HYPDH
Nean.SNP

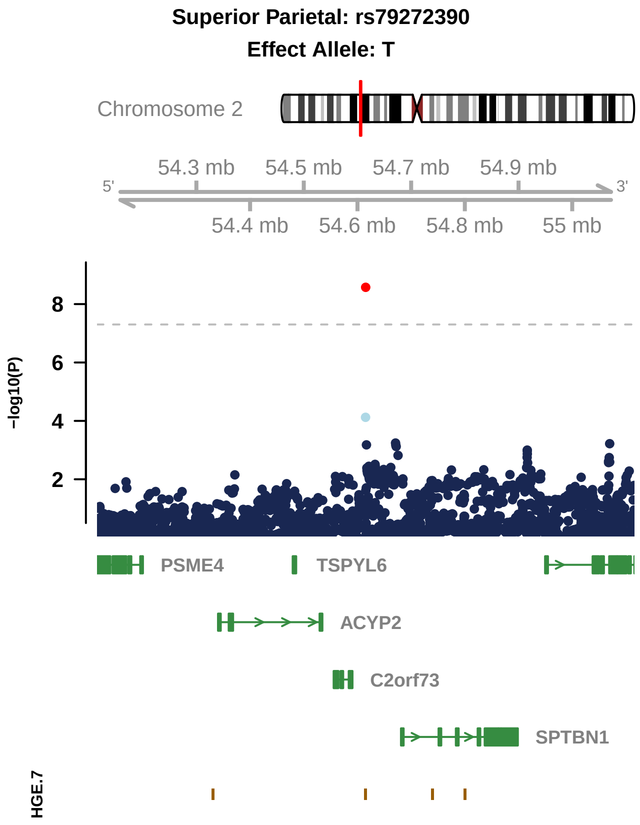

Superior Parietal: rs79272390
Effect Allele: T
Chromosome 2
54.3 mb
54.5 mb
54.7 mb
54.9 mb
5'
3'
54.4 mb
54.6 mb
54.8 mb
55 mb
-log10(P)
8
6
4
2
PSME4
TSPYL6
ACYP2
C2orf73
SPTBN1
HGE.7

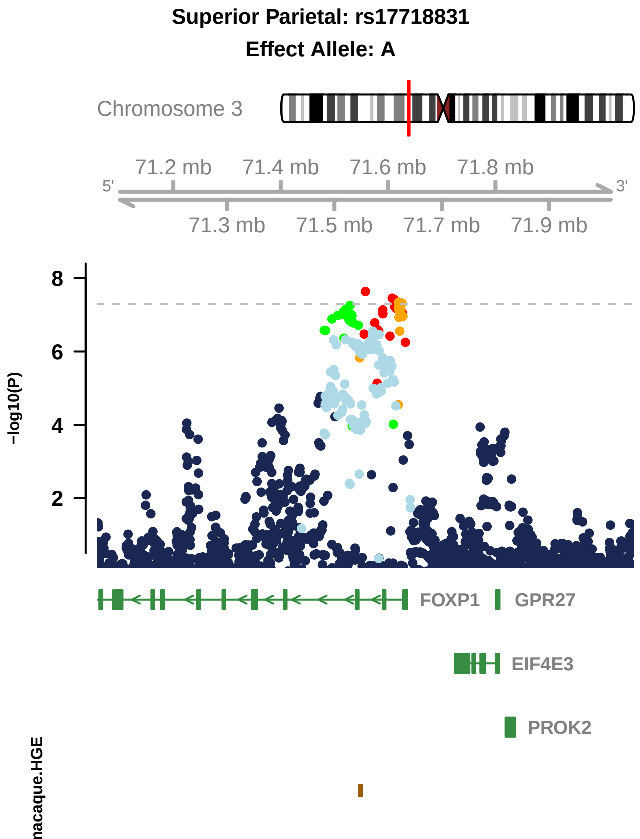

Superior Parietal: rs17718831
Effect Allele: A
Chromosome 3
71.2 mb
71.4 mb
71.6 mb
71.8 mb
5'
3'
71.3 mb
71.5 mb
71.7 mb
71.9 mb
8
6
4
2
−log10(P)
FOXP1
GPR27
EIF4E3
PROK2
macaque.HGE

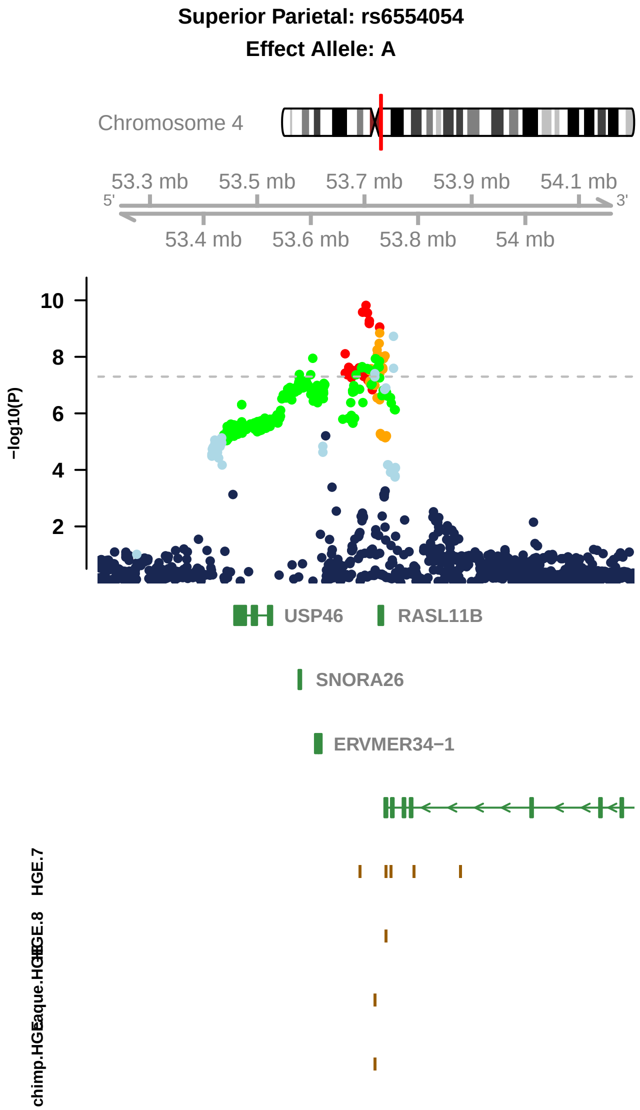

Superior Parietal: rs6554054
Effect Allele: A
Chromosome 4
53.3 mb
53.5 mb
53.7 mb
53.9 mb
54.1 mb
5'
3'
53.4 mb
53.6 mb
53.8 mb
54 mb
−log10(P)
10
8
6
4
2
USP46
RASL11B
SNORA26
ERVMER34−1
HGE.7
HGE.8
macaque.HGE
chimp.HGE

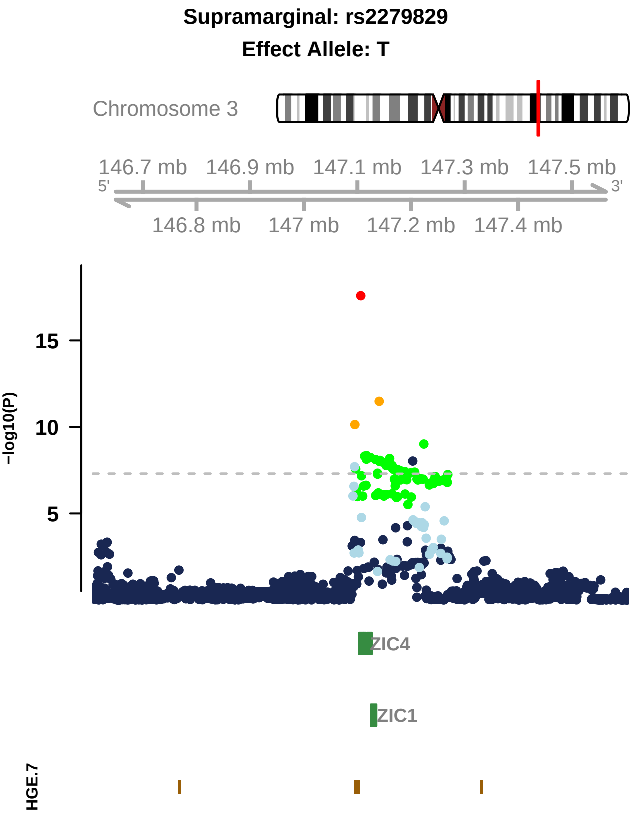

Supramarginal: rs2279829
Effect Allele: T
Chromosome 3
146.7 mb
146.9 mb
147.1 mb
147.3 mb
147.5 mb
5'
3'
146.8 mb
147 mb
147.2 mb
147.4 mb
−log10(P)
15
10
5
ZIC4
ZIC1
HGE.7

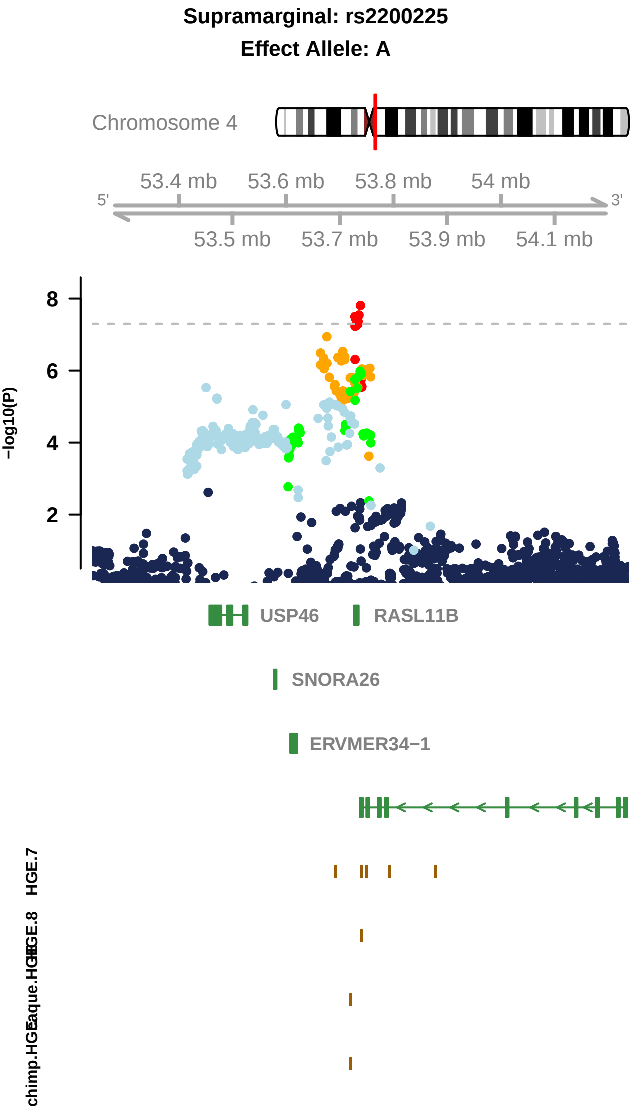

Supramarginal: rs2200225
Effect Allele: A
Chromosome 4
53.4 mb
53.6 mb
53.8 mb
54 mb
5'
3'
53.5 mb
53.7 mb
53.9 mb
54.1 mb
8
6
4
2
−log10(P)
USP46
RASL11B
SNORA26
ERVMER34−1
HGE.7
HGE.8

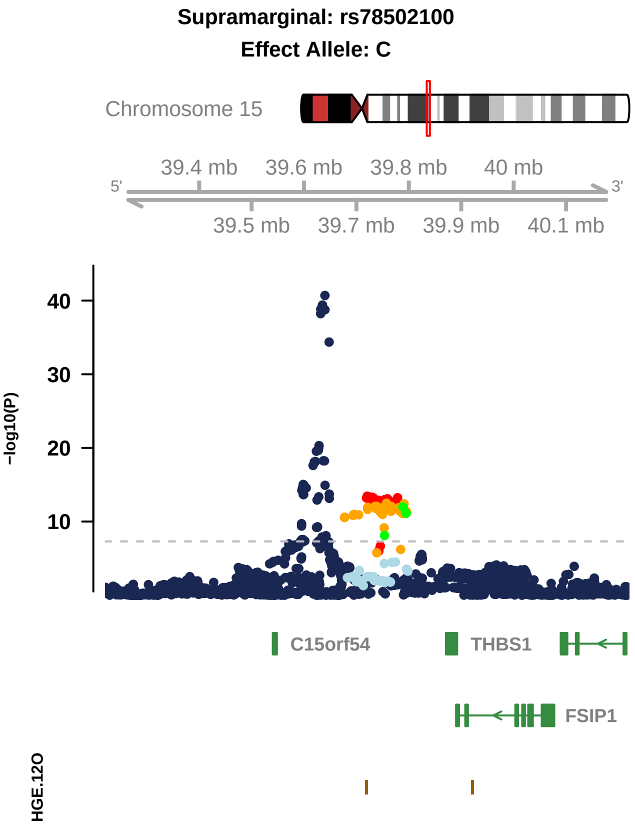

Supramarginal: rs78502100
Effect Allele: C
Chromosome 15
39.4 mb
39.6 mb
39.8 mb
40 mb
5'
3'
39.5 mb
39.7 mb
39.9 mb
40.1 mb
40
30
20
10
−log10(P)
C15orf54
THBS1
FSIP1
HGE.12O

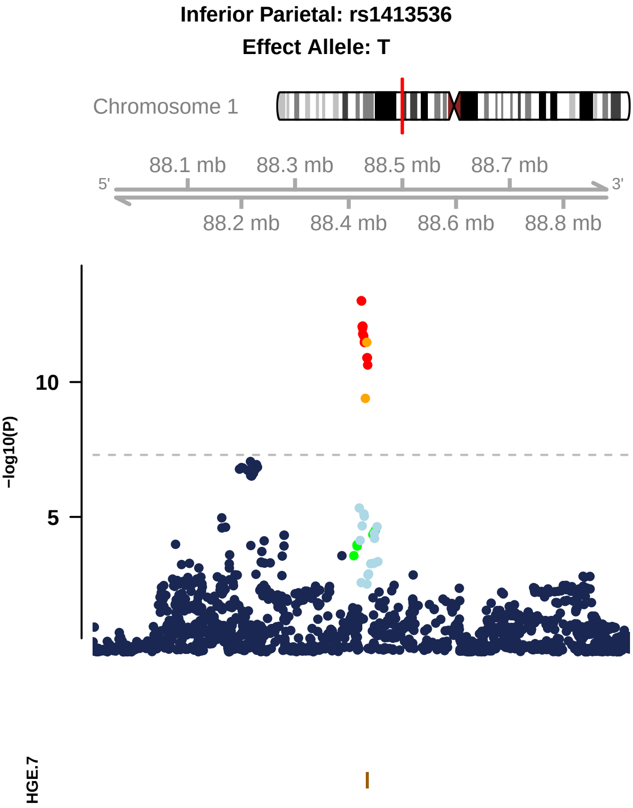

Inferior Parietal: rs1413536
Effect Allele: T
Chromosome 1
88.1 mb
88.3 mb
88.5 mb
88.7 mb
5'
3'
88.2 mb
88.4 mb
88.6 mb
88.8 mb
10
5
-log10(P)
HGE.7

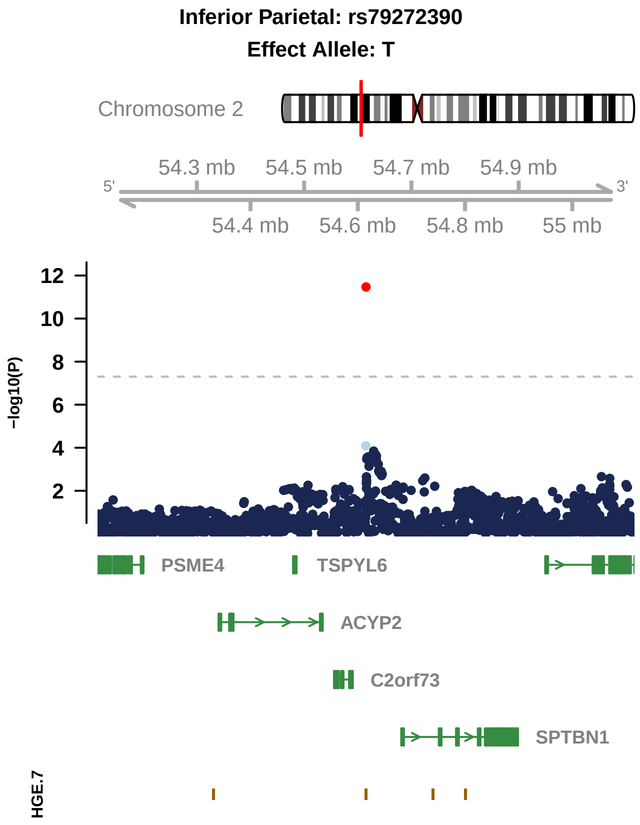

Inferior Parietal: rs79272390
Effect Allele: T
Chromosome 2
54.3 mb
54.5 mb
54.7 mb
54.9 mb
5'
3'
54.4 mb
54.6 mb
54.8 mb
55 mb
−log10(P)
12
10
8
6
4
2
PSME4
TSPYL6
ACYP2
C2orf73
SPTBN1
HGE.7

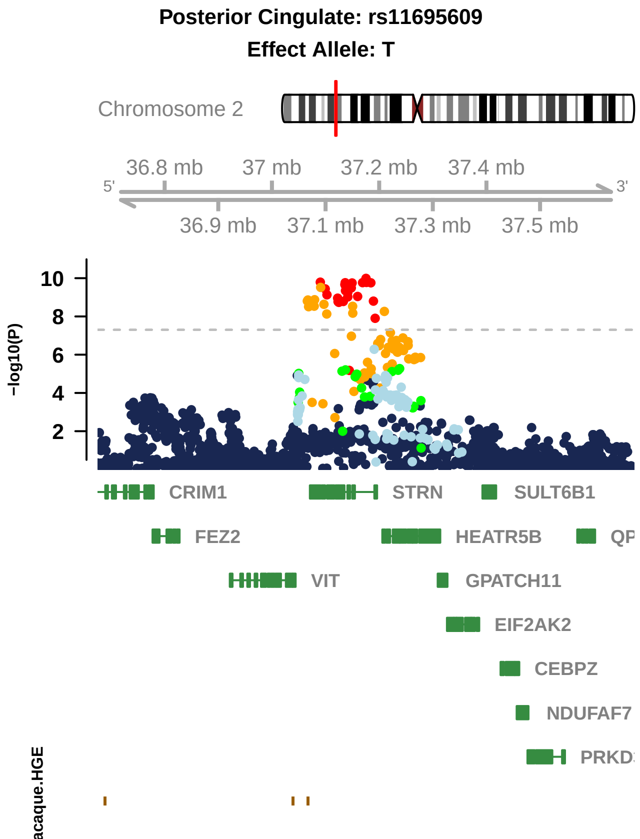

Posterior Cingulate: rs11695609
Effect Allele: T
Chromosome 2
36.8 mb
37 mb
37.2 mb
37.4 mb
5'
3'
36.9 mb
37.1 mb
37.3 mb
37.5 mb
10
8
6
4
2
−log10(P)
CRIM1
STRN
SULT6B1
FEZ2
HEATR5B
QP
VIT
GPATCH11
EIF2AK2
CEBPZ
NDUFAF7
PRKD
acaque.HGE

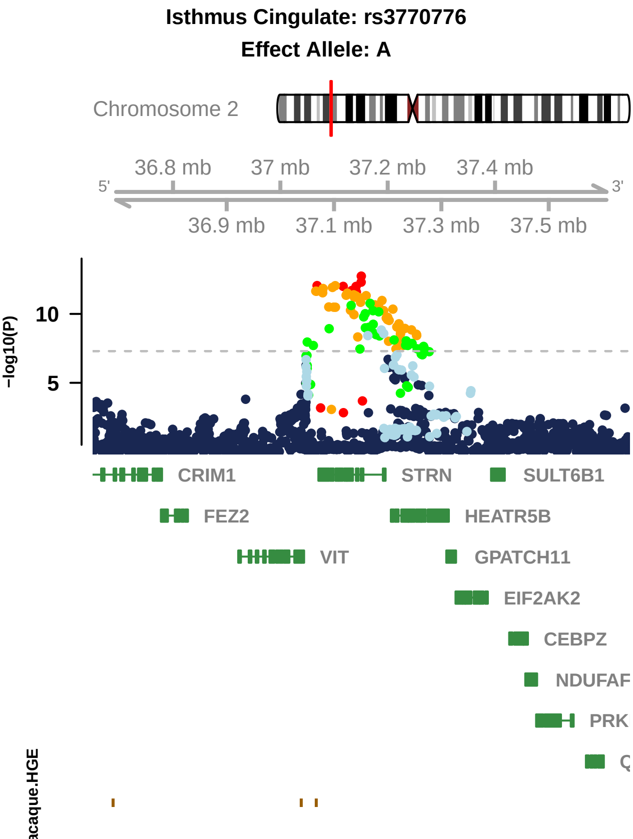

Isthmus Cingulate: rs3770776
Effect Allele: A
Chromosome 2
36.8 mb
37 mb
37.2 mb
37.4 mb
5'
3'
36.9 mb
37.1 mb
37.3 mb
37.5 mb
−log10(P)
10
5
CRIM1
STRN
SULT6B1
FEZ2
HEATR5B
VIT
GPATCH11
EIF2AK2
CEBPZ
NDUFAF
PRK
acaque.HGE

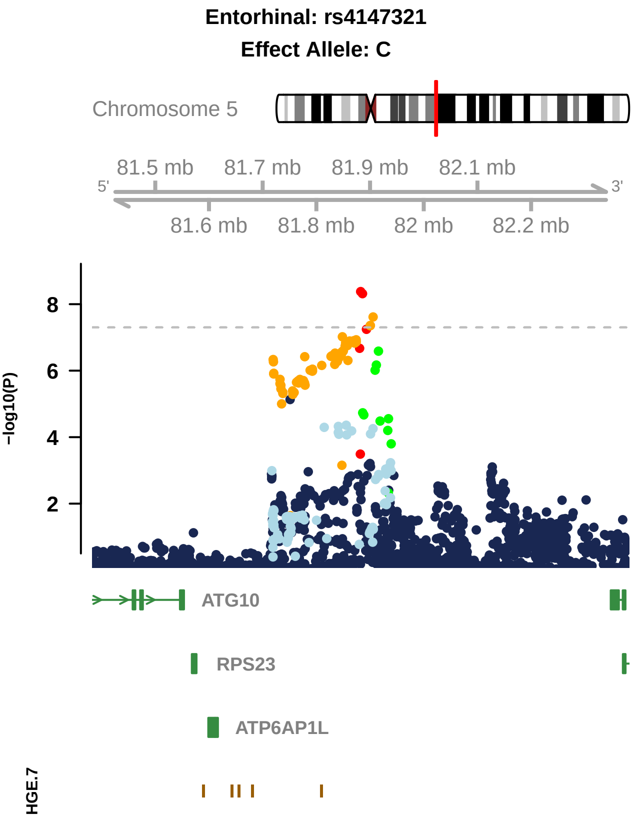

Entorhinal: rs4147321
Effect Allele: C
Chromosome 5
81.5 mb
81.7 mb
81.9 mb
82.1 mb
5'
3'
81.6 mb
81.8 mb
82 mb
82.2 mb
8
6
4
2
−log10(P)
ATG10
RPS23
ATP6AP1L
HGE.7

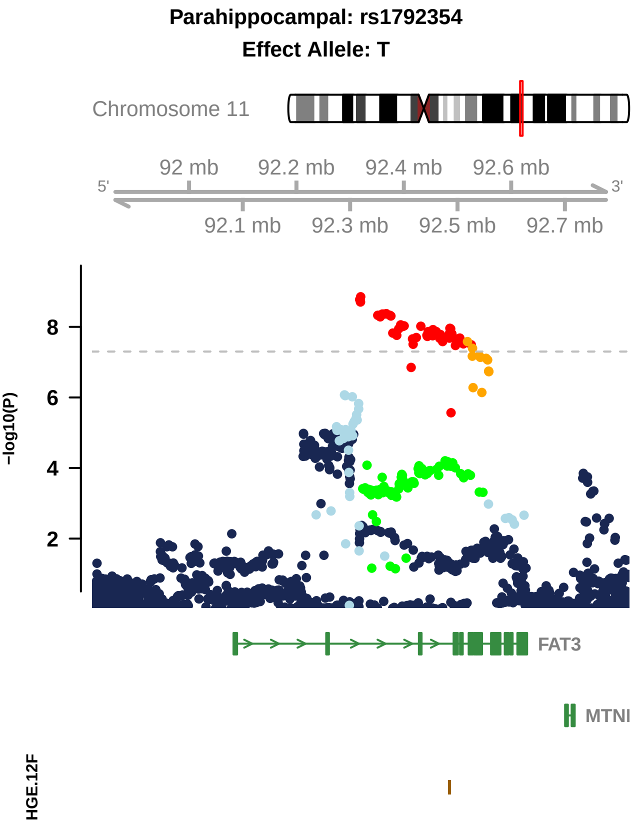

Parahippocampal: rs1792354
Effect Allele: T
Chromosome 11
92 mb
92.2 mb
92.4 mb
92.6 mb
5'
3'
92.1 mb
92.3 mb
92.5 mb
92.7 mb
8
6
4
2
−log10(P)
FAT3
MTNI
HGE.12F

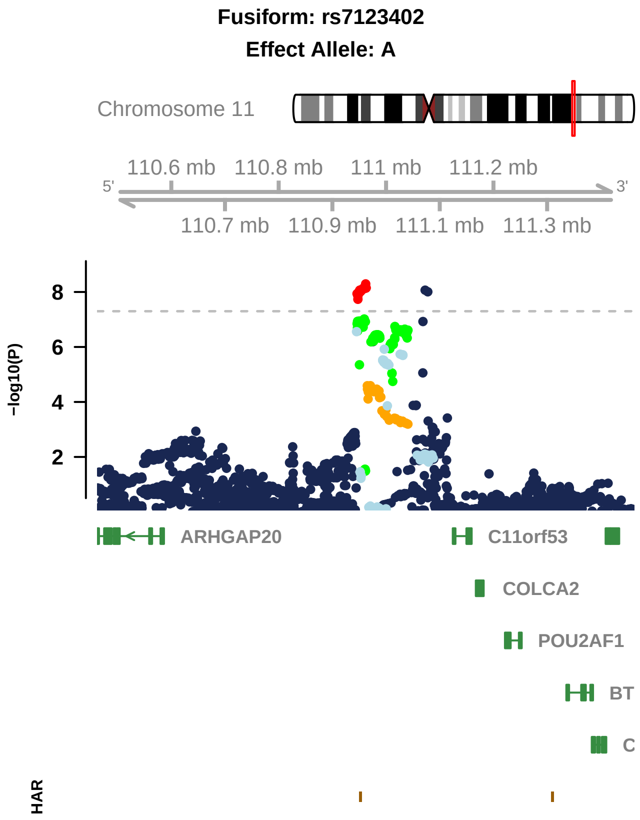

Fusiform: rs7123402
Effect Allele: A
Chromosome 11
110.6 mb
110.8 mb
111 mb
111.2 mb
5'
3'
110.7 mb
110.9 mb
111.1 mb
111.3 mb
−log10(P)
8
6
4
2
ARHGAP20
C11orf53
COLCA2
POU2AF1
BT
C
HAR

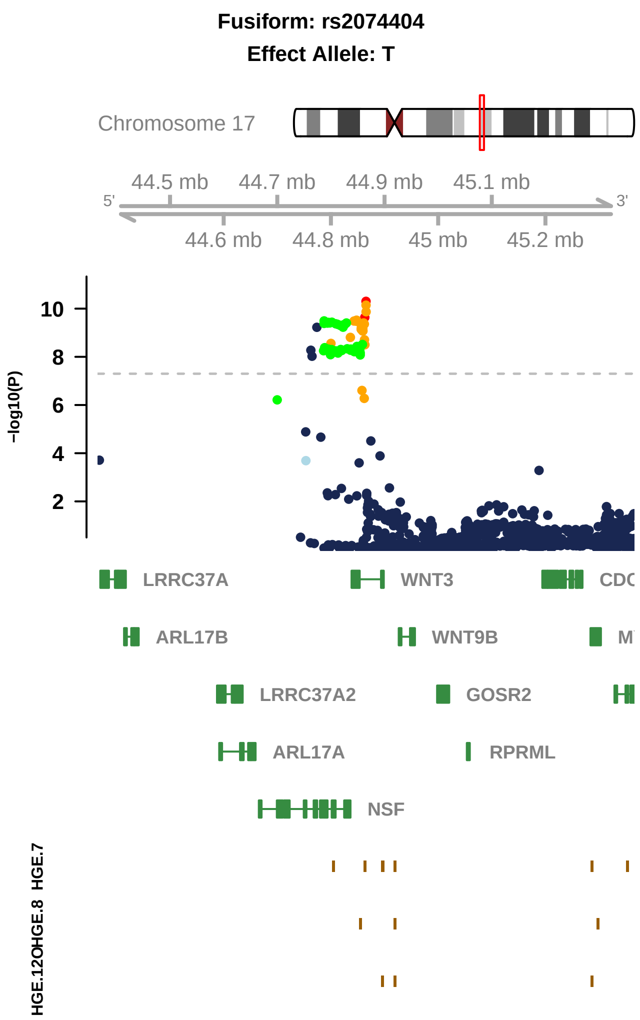
Fusiform: rs2074404
Effect Allele: T
Chromosome 17
44.5 mb
44.7 mb
44.9 mb
45.1 mb
5'
3'
44.6 mb
44.8 mb
45 mb
45.2 mb
−log10(P)
10
8
6
4
2
LRRC37A
WNT3
ARL17B
WNT9B
LRRC37A2
GOSR2
ARL17A
RPRML
NSF
HGE.7
HGE.8
HGE.12OH

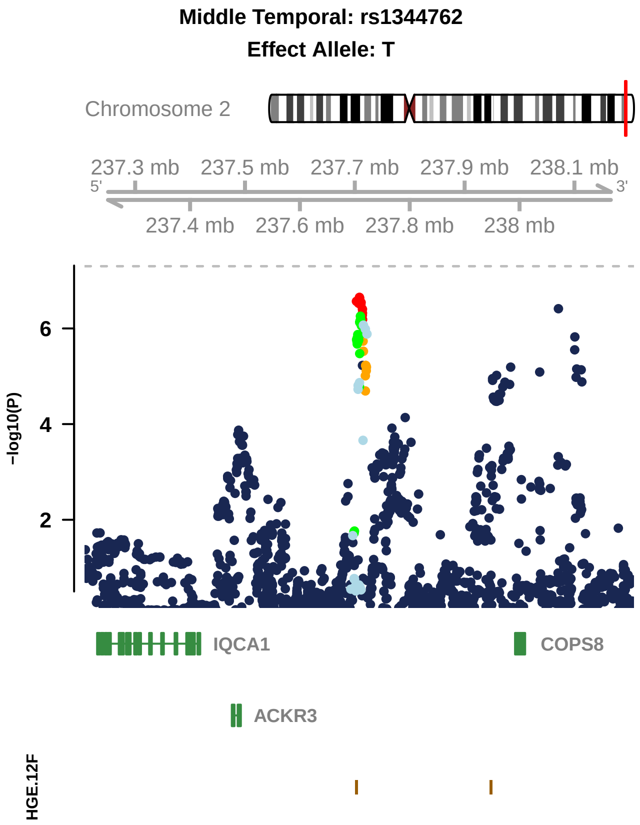

Middle Temporal: rs1344762
Effect Allele: T
Chromosome 2
237.3 mb
237.5 mb
237.7 mb
237.9 mb
238.1 mb
5'
3'
237.4 mb
237.6 mb
237.8 mb
238 mb
6
4
2
−log10(P)
IQCA1
COPS8
ACKR3
HGE.12F

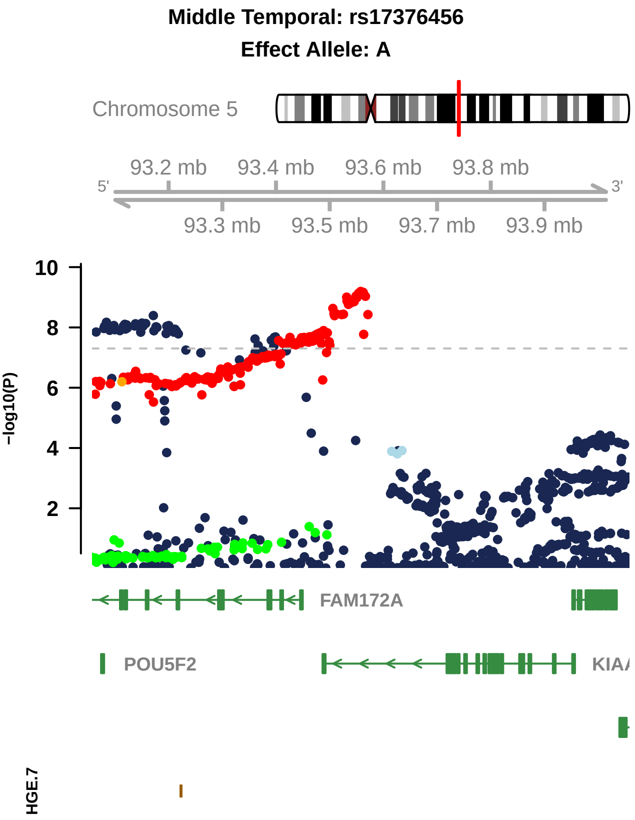
Middle Temporal: rs17376456
Effect Allele: A
Chromosome 5
93.2 mb
93.4 mb
93.6 mb
93.8 mb
5'
3'
93.3 mb
93.5 mb
93.7 mb
93.9 mb
10
8
6
4
2
−log10(P)
FAM172A
POU5F2
KIAA
HGE.7

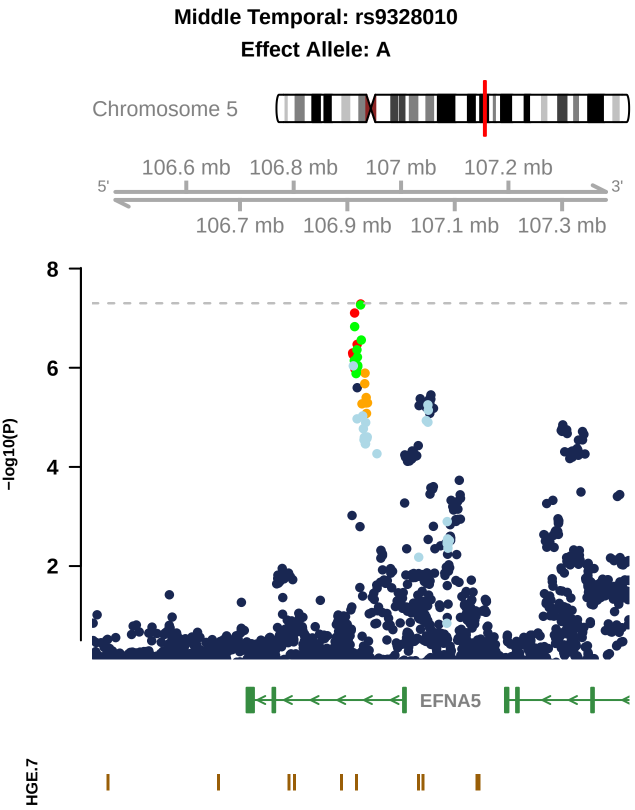

Middle Temporal: rs9328010
Effect Allele: A
Chromosome 5
106.6 mb
106.8 mb
107 mb
107.2 mb
5'
3'
106.7 mb
106.9 mb
107.1 mb
107.3 mb
8
6
4
2
-log10(P)
EFNA5
HGE.7

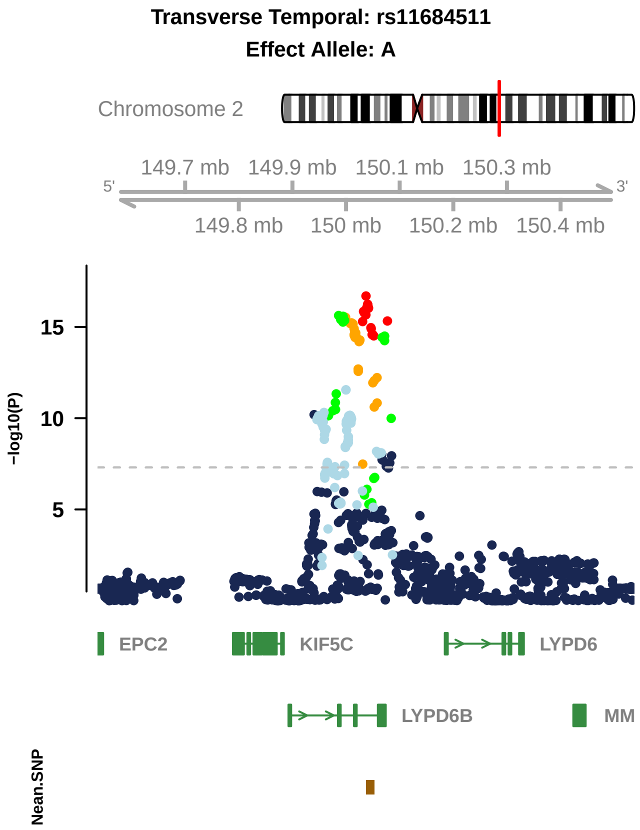
Transverse Temporal: rs11684511
Effect Allele: A
Chromosome 2
149.7 mb
149.9 mb
150.1 mb
150.3 mb
5'
3'
149.8 mb
150 mb
150.2 mb
150.4 mb
-log10(P)
15
10
5
EPC2
KIF5C
LYPD6
LYPD6B
MM
Nean.SNP

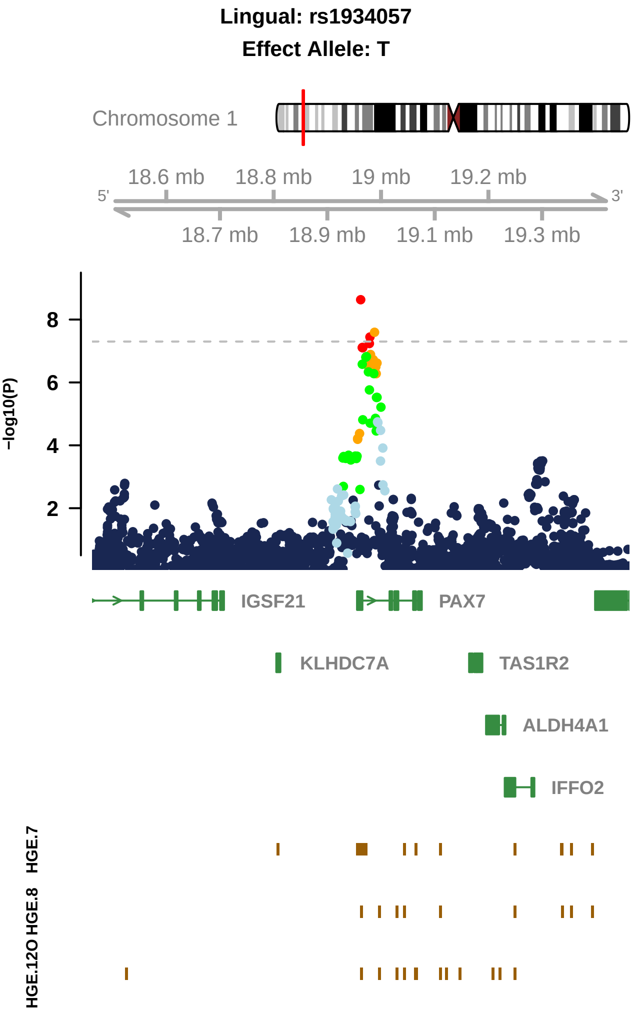

Lingual: rs1934057
Effect Allele: T
Chromosome 1
18.6 mb
18.8 mb
19 mb
19.2 mb
5'
3'
18.7 mb
18.9 mb
19.1 mb
19.3 mb
−log10(P)
8
6
4
2
IGSF21
PAX7
KLHDC7A
TAS1R2
ALDH4A1
IFFO2
HGE.7
HGE.8
HGE.12O

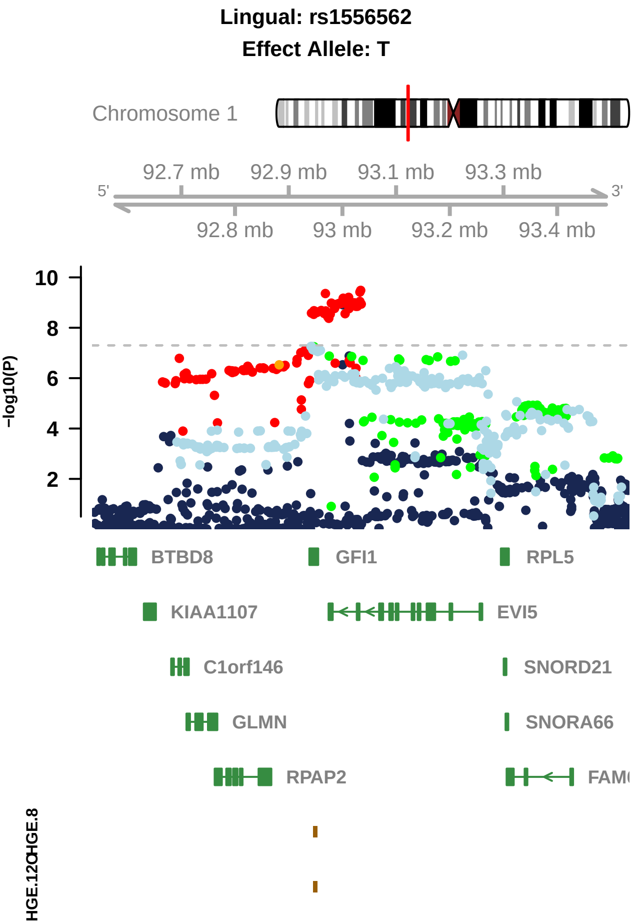

Lingual: rs1556562
Effect Allele: T
Chromosome 1
92.7 mb
92.9 mb
93.1 mb
93.3 mb
5'
3'
92.8 mb
93 mb
93.2 mb
93.4 mb
10
8
6
4
2
−log10(P)
BTBD8
GFI1
RPL5
KIAA1107
EVI5
C1orf146
SNORD21
GLMN
SNORA66
RPAP2
HGE.8

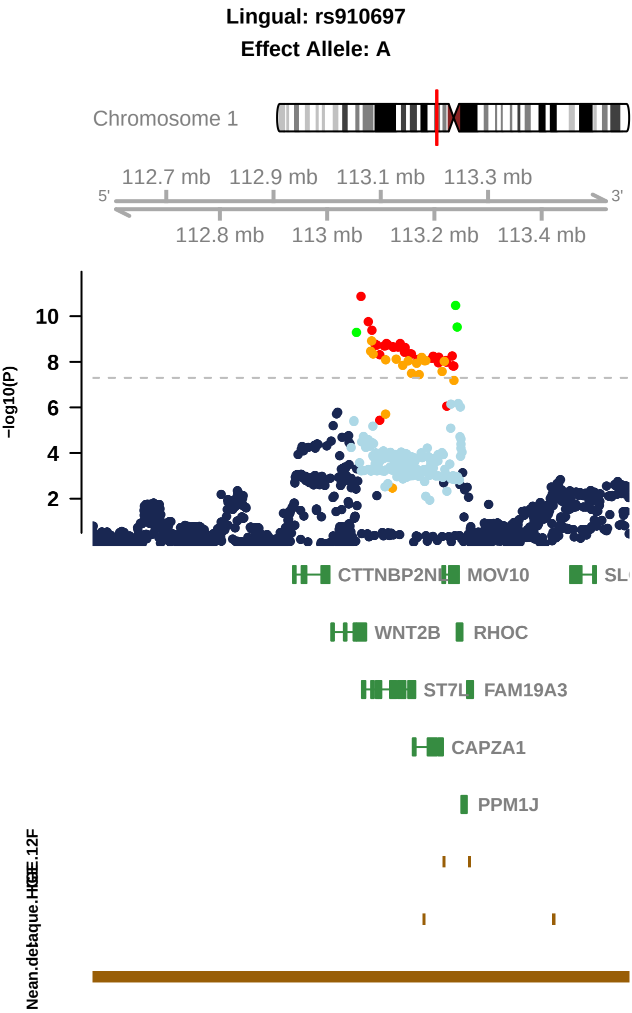
Lingual: rs910697
Effect Allele: A
Chromosome 1
112.7 mb
112.9 mb
113.1 mb
113.3 mb
5'
3'
112.8 mb
113 mb
113.2 mb
113.4 mb
-log10(P)
10
8
6
4
2
CTTNBP2NL
MOV10
WNT2B
RHOC
ST7L
FAM19A3
CAPZA1
PPM1J

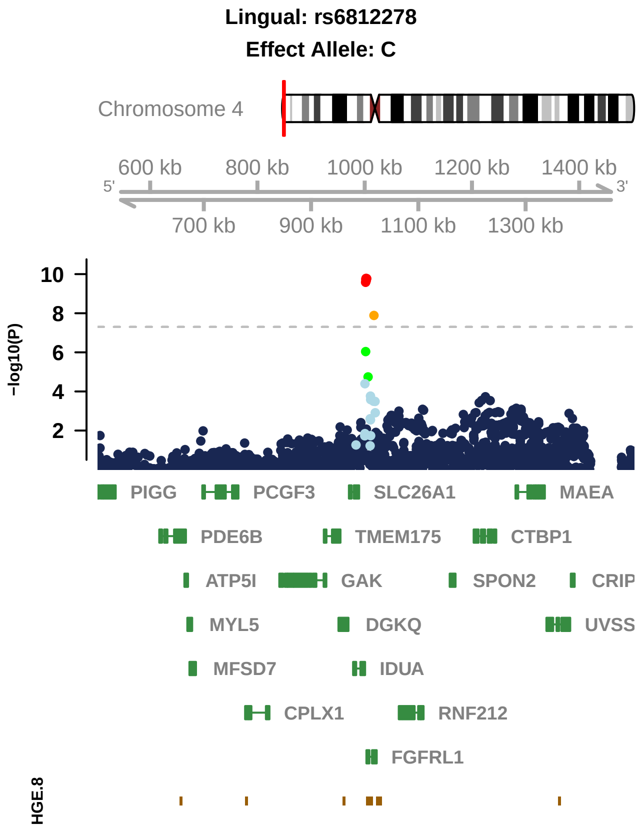

Lingual: rs6812278
Effect Allele: C
Chromosome 4
600 kb
800 kb
1000 kb
1200 kb
1400 kb
5'
3'
700 kb
900 kb
1100 kb
1300 kb
10
8
6
4
2
−log10(P)
PIGG
PCGF3
SLC26A1
MAEA
PDE6B
TMEM175
CTBP1
ATP5I
GAK
SPON2
CRIP
MYL5
DGKQ
UVSS
MFSD7
IDUA
CPLX1
RNF212
FGFRL1
HGE.8

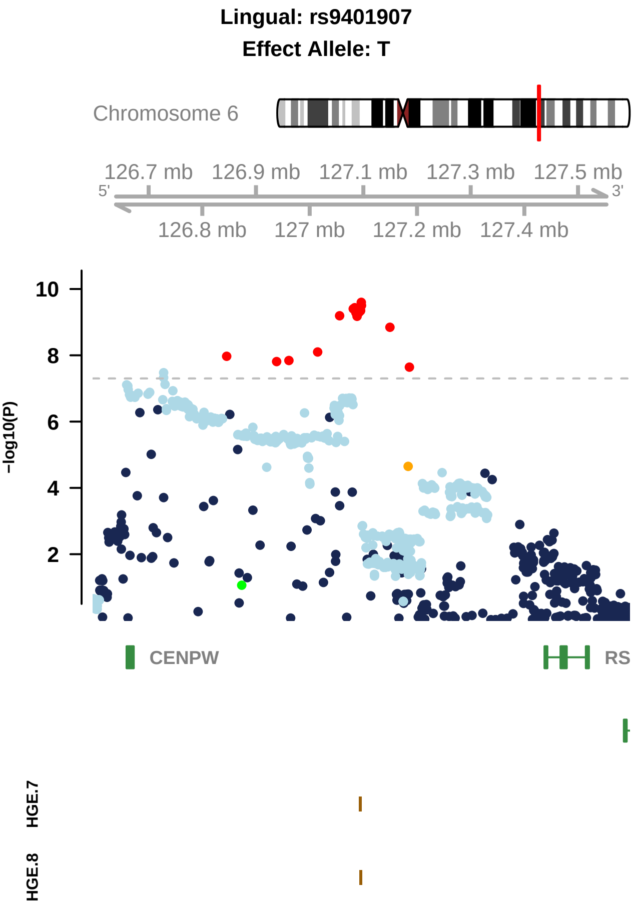

Lingual: rs9401907
Effect Allele: T
Chromosome 6
126.7 mb
126.9 mb
127.1 mb
127.3 mb
127.5 mb
5'
3'
126.8 mb
127 mb
127.2 mb
127.4 mb
−log10(P)
10
8
6
4
2
CENPW
RS
HGE.7
HGE.8

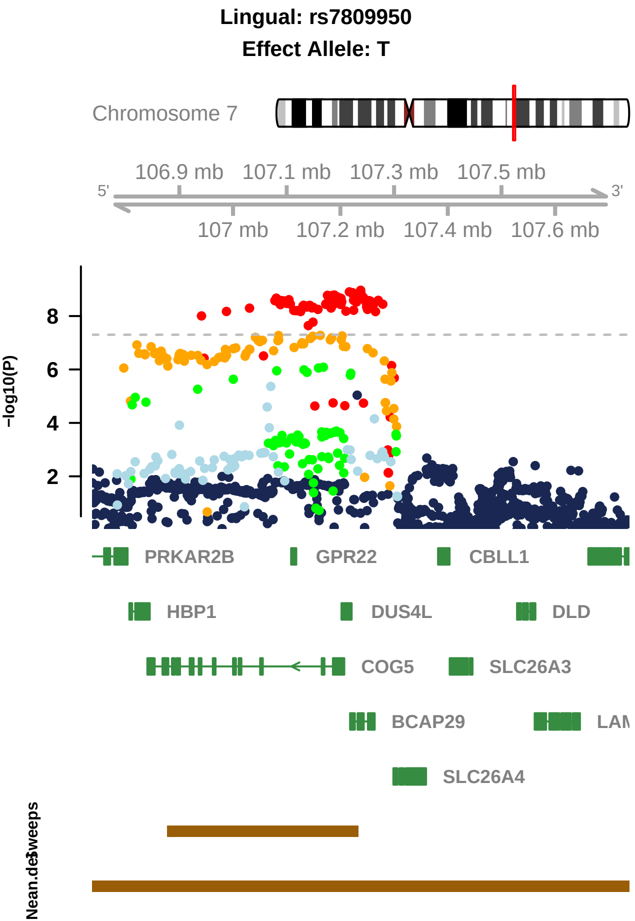
Lingual: rs7809950
Effect Allele: T
Chromosome 7
106.9 mb
107.1 mb
107.3 mb
107.5 mb
5'
3'
107 mb
107.2 mb
107.4 mb
107.6 mb
−log10(P)
8
6
4
2
PRKAR2B
GPR22
CBLL1
HBP1
DUS4L
DLD
COG5
SLC26A3
BCAP29
SLC26A4
Nean.des
Sweeps

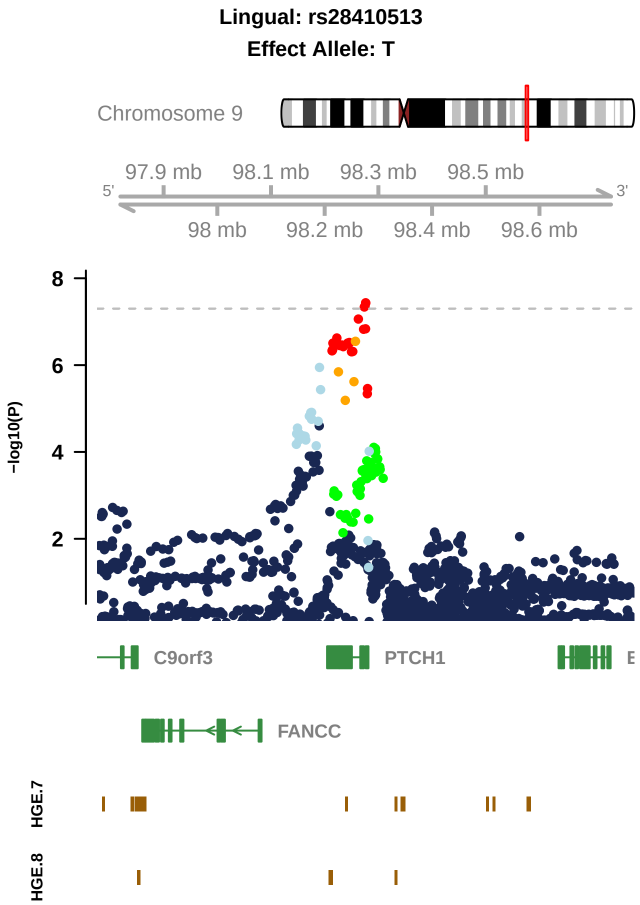
Lingual: rs28410513
Effect Allele: T
Chromosome 9
97.9 mb
98.1 mb
98.3 mb
98.5 mb
5'
3'
98 mb
98.2 mb
98.4 mb
98.6 mb
8
6
4
2
-log10(P)
C9orf3
PTCH1
FANCC
HGE.7
HGE.8

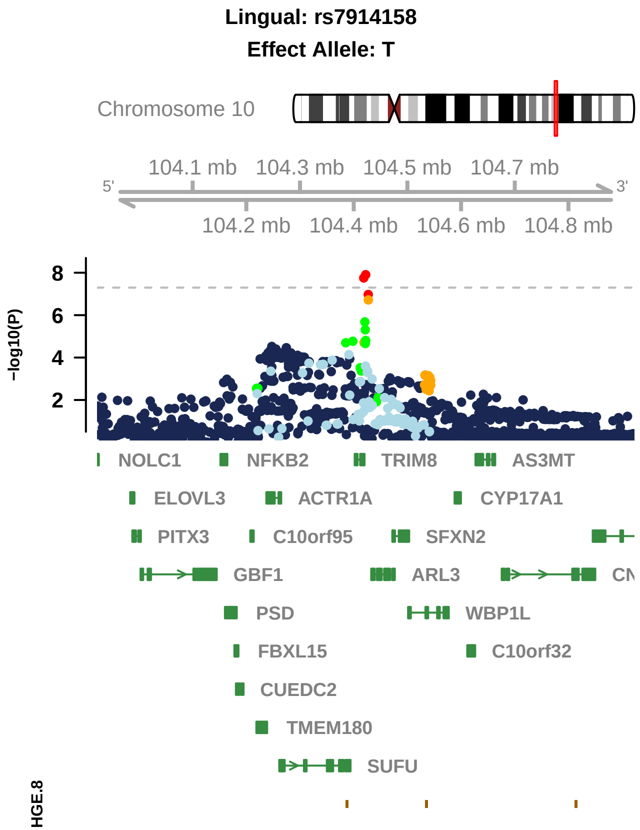
Lingual: rs7914158
Effect Allele: T
Chromosome 10
104.1 mb
104.3 mb
104.5 mb
104.7 mb
5'
3'
104.2 mb
104.4 mb
104.6 mb
104.8 mb
-log10(P)
8
6
4
2
NOLC1
NFKB2
TRIM8
AS3MT
ELOVL3
ACTR1A
CYP17A1
PITX3
C10orf95
SFXN2
GBF1
ARL3
PSD
WBP1L
FBXL15
C10orf32
CUEDC2
TMEM180
SUFU
HGE.8

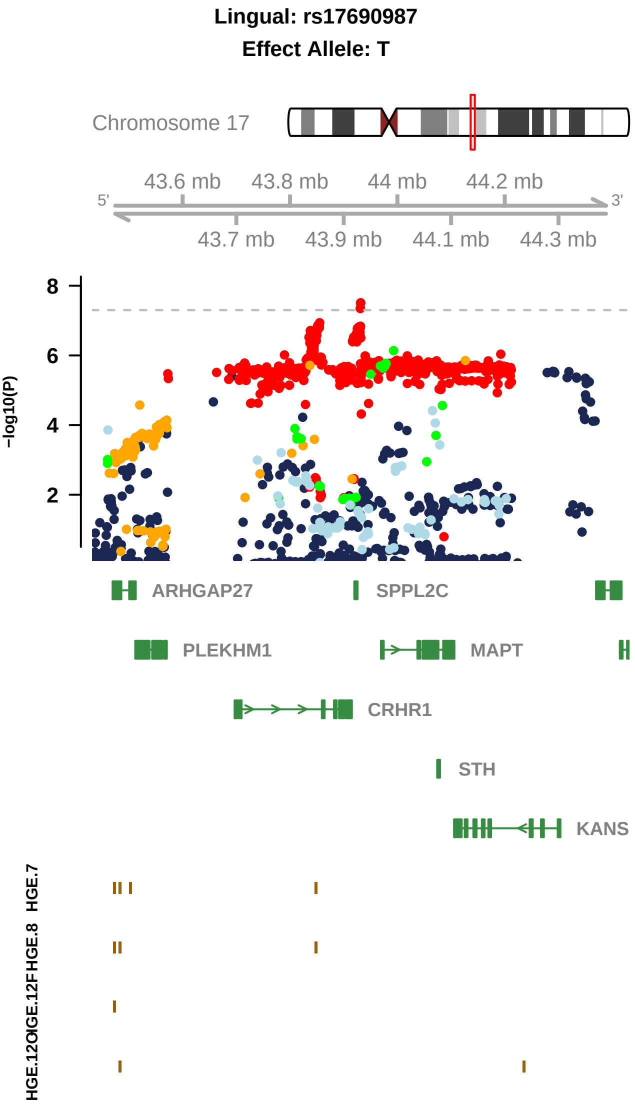

Lingual: rs17690987
Effect Allele: T
Chromosome 17
43.6 mb
43.8 mb
44 mb
44.2 mb
5'
3'
43.7 mb
43.9 mb
44.1 mb
44.3 mb
-log10(P)
8
6
4
2
ARHGAP27
SPPL2C
PLEKHM1
MAPT
CRHR1
STH
KANS
HGE.7
HGE.8
HGE.12F

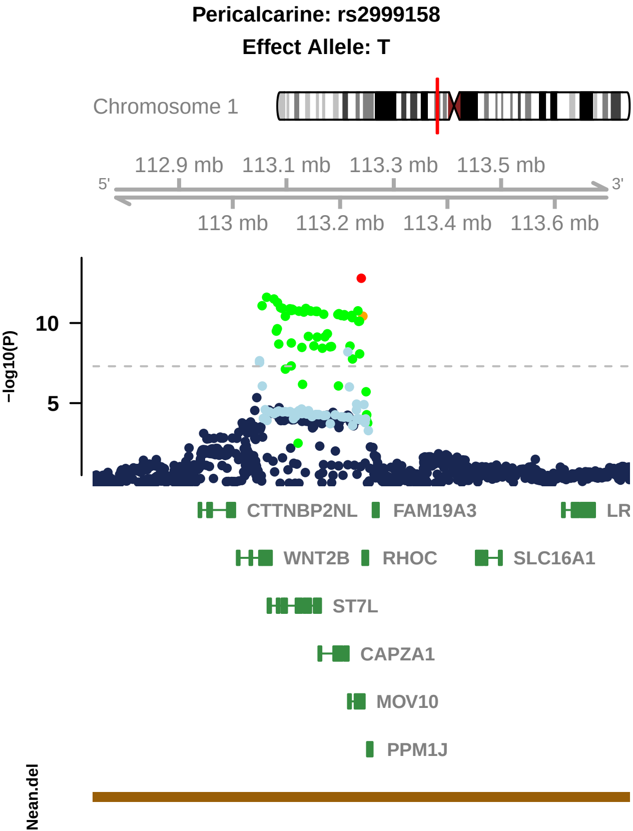

Pericalcarine: rs2999158
Effect Allele: T
Chromosome 1
112.9 mb
113.1 mb
113.3 mb
113.5 mb
5'
3'
113 mb
113.2 mb
113.4 mb
113.6 mb
−log10(P)
10
5
CTTNBP2NL
FAM19A3
LR
WNT2B
RHOC
SLC16A1
ST7L
CAPZA1
MOV10
PPM1J
Nean.del

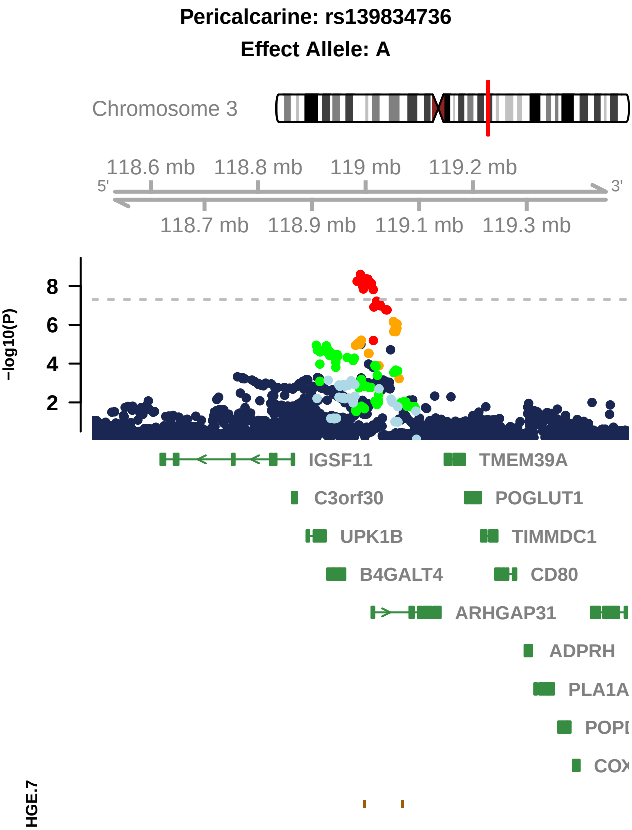

Pericalcarine: rs139834736
Effect Allele: A
Chromosome 3
118.6 mb
118.8 mb
119 mb
119.2 mb
5'
3'
118.7 mb
118.9 mb
119.1 mb
119.3 mb
−log10(P)
8
6
4
2
IGSF11
TMEM39A
C3orf30
POGLUT1
UPK1B
TIMMDC1
B4GALT4
CD80
ARHGAP31
ADPRH
PLA1A
HGE.7

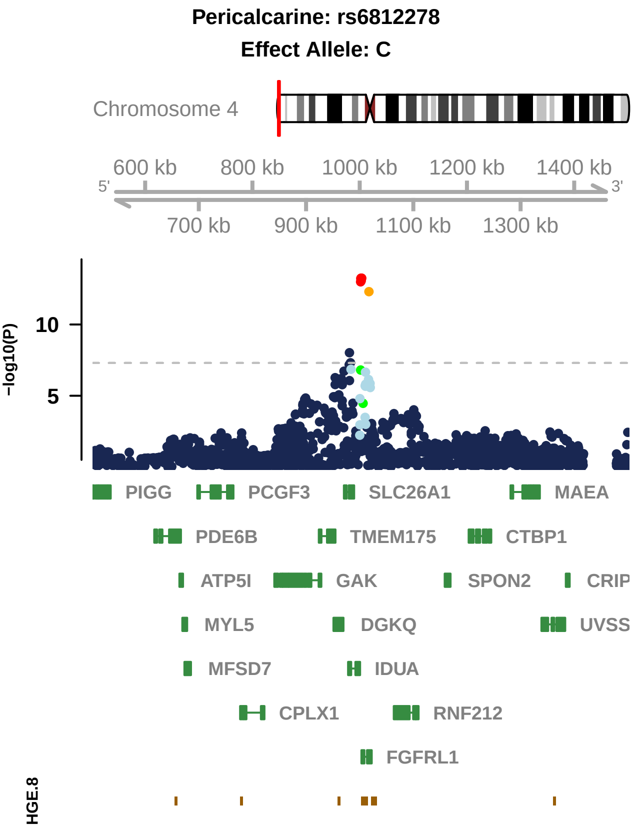
Pericalcarine: rs6812278
Effect Allele: C
Chromosome 4
600 kb
800 kb
1000 kb
1200 kb
1400 kb
5'
3'
700 kb
900 kb
1100 kb
1300 kb
−log10(P)
10
5
PIGG
PCGF3
SLC26A1
MAEA
PDE6B
TMEM175
CTBP1
ATP5I
GAK
SPON2
CRIP
MYL5
DGKQ
UVSS
MFSD7
IDUA
CPLX1
RNF212
FGFRL1
HGE.8

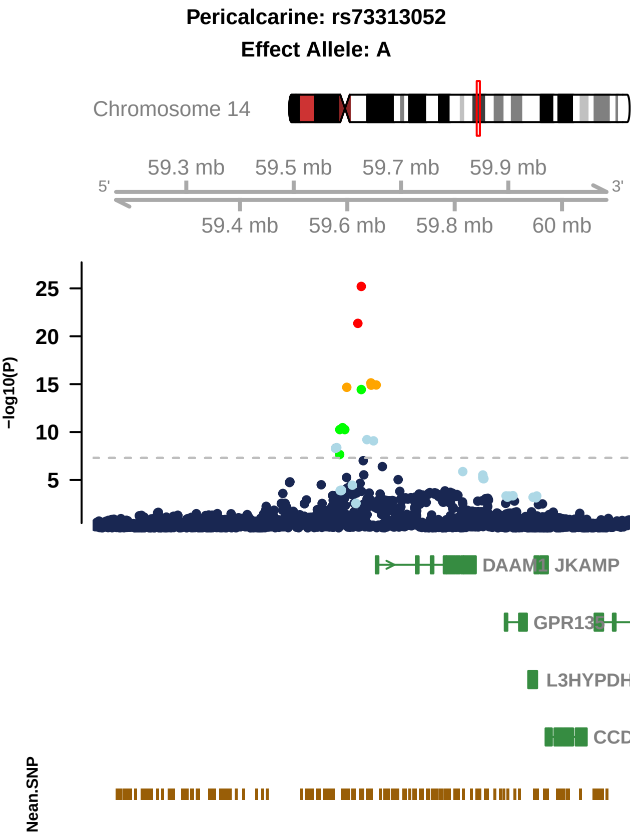
Pericalcarine: rs73313052
Effect Allele: A
Chromosome 14
59.3 mb
59.5 mb
59.7 mb
59.9 mb
5'
3'
59.4 mb
59.6 mb
59.8 mb
60 mb
−log10(P)
25
20
15
10
5
DAAM1
JKAMP
GPR135
L3HYPDH
CCD
Nean.SNP

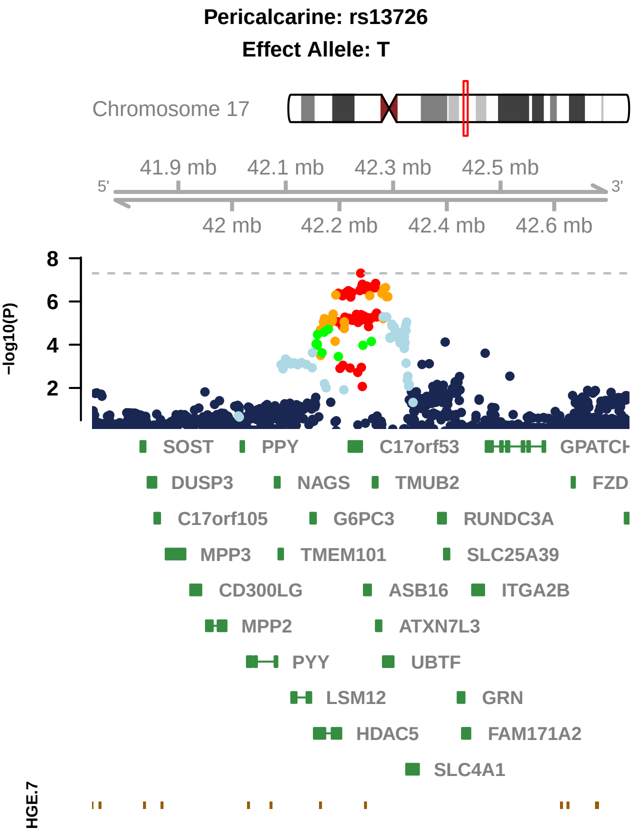

Pericalcarine: rs13726
Effect Allele: T
Chromosome 17
41.9 mb
42.1 mb
42.3 mb
42.5 mb
5'
3'
42 mb
42.2 mb
42.4 mb
42.6 mb
-log10(P)
8
6
4
2
SOST
PPY
C17orf53
DUSP3
NAGS
TMUB2
C17orf105
G6PC3
RUNDC3A
MPP3
TMEM101
SLC25A39
CD300LG
ASB16
ITGA2B
MPP2
ATXN7L3
PYY
UBTF
LSM12
GRN
HDAC5
FAM171A2
SLC4A1
HGE.7

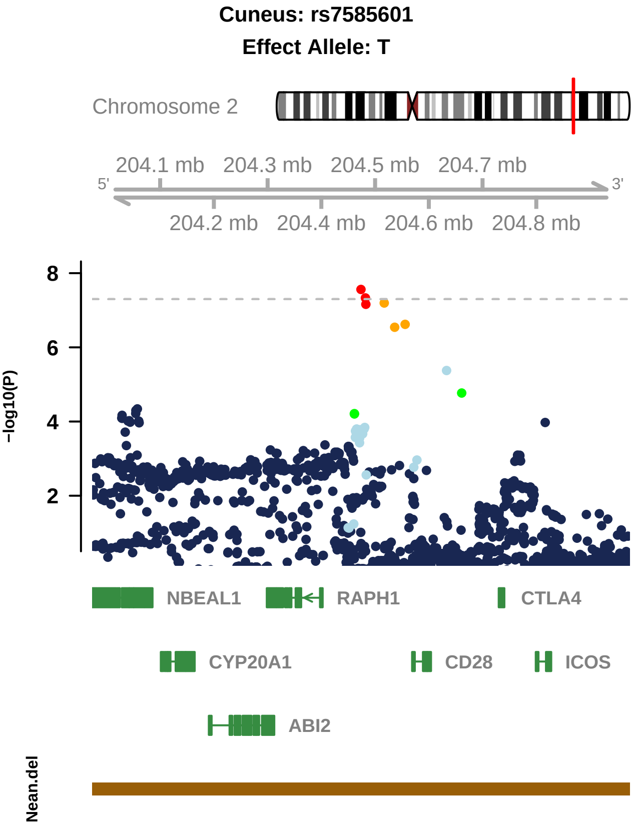

Cuneus: rs7585601
Effect Allele: T
Chromosome 2
204.1 mb
204.3 mb
204.5 mb
204.7 mb
5'
3'
204.2 mb
204.4 mb
204.6 mb
204.8 mb
8
6
4
2
−log10(P)
NBEAL1
RAPH1
CTLA4
CYP20A1
CD28
ICOS
ABI2
Nean.del

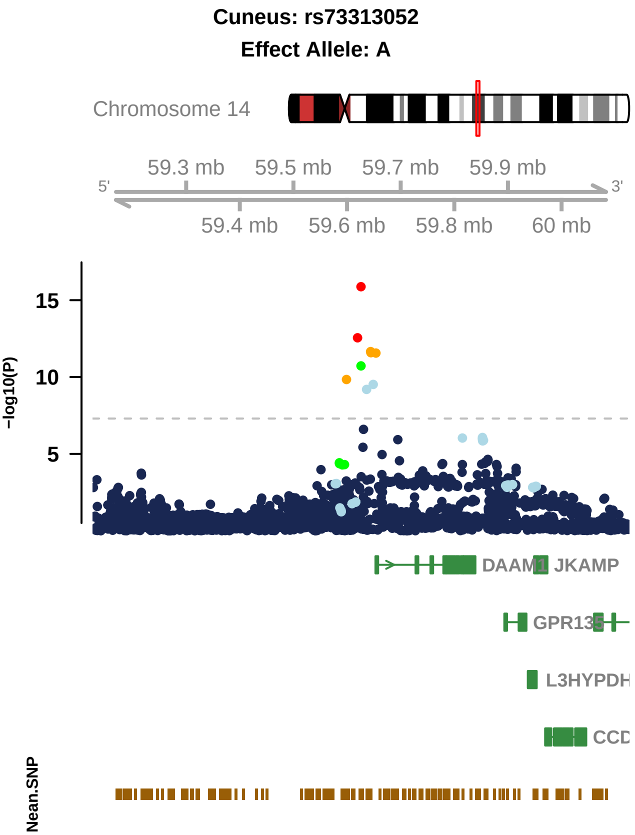

Cuneus: rs73313052
Effect Allele: A
Chromosome 14
59.3 mb
59.5 mb
59.7 mb
59.9 mb
5'
3'
59.4 mb
59.6 mb
59.8 mb
60 mb
15
10
5
-log10(P)
DAAM1
JKAMP
GPR135
L3HYPDH
CCD
Nean.SNP

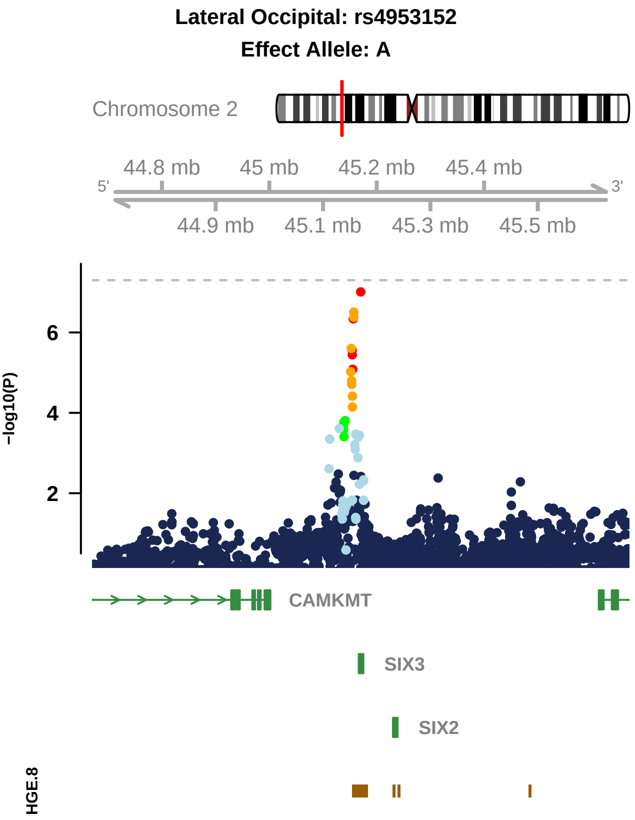
Lateral Occipital: rs4953152
Effect Allele: A
Chromosome 2
44.8 mb
45 mb
45.2 mb
45.4 mb
5'
3'
44.9 mb
45.1 mb
45.3 mb
45.5 mb
6
4
2
−log10(P)
CAMKMT
SIX3
SIX2
HGE.8

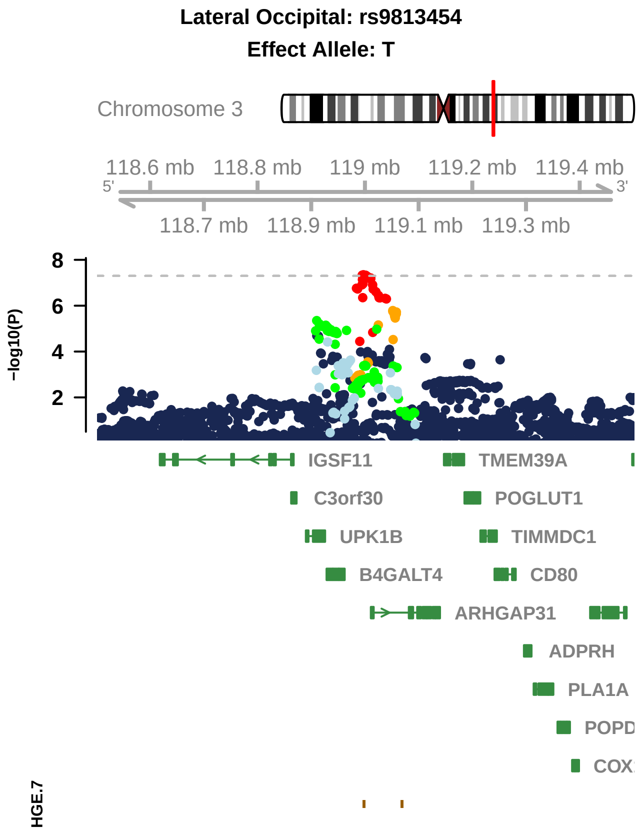

Lateral Occipital: rs9813454
Effect Allele: T
Chromosome 3
118.6 mb
118.8 mb
119 mb
119.2 mb
119.4 mb
5'
3'
118.7 mb
118.9 mb
119.1 mb
119.3 mb
−log10(P)
8
6
4
2
IGSF11
TMEM39A
C3orf30
POGLUT1
UPK1B
TIMMDC1
B4GALT4
CD80
ARHGAP31
ADPRH
PLA1A
HGE.7

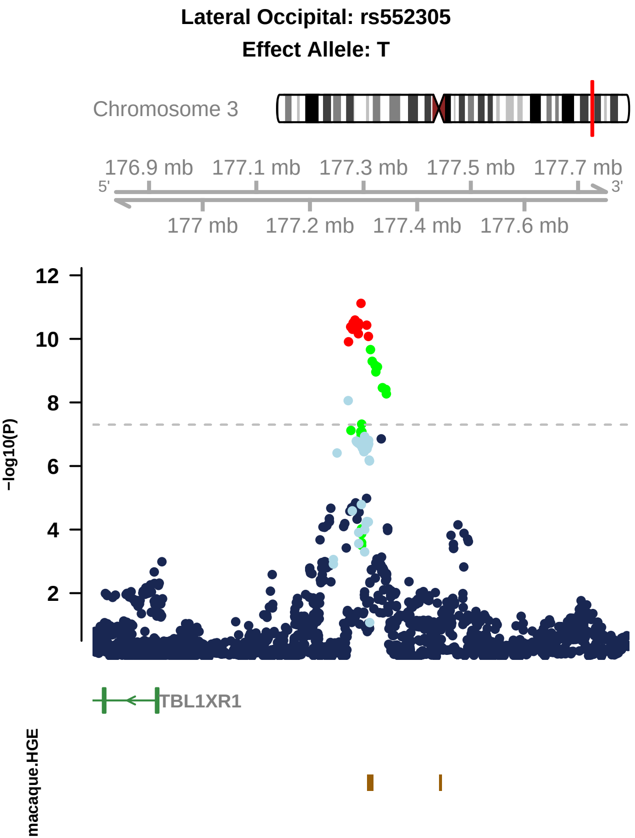

Lateral Occipital: rs552305
Effect Allele: T
Chromosome 3
176.9 mb
177.1 mb
177.3 mb
177.5 mb
177.7 mb
5'
3'
177 mb
177.2 mb
177.4 mb
177.6 mb
12
10
8
6
4
2
-log10(P)
TBL1XR1
macaque.HGE

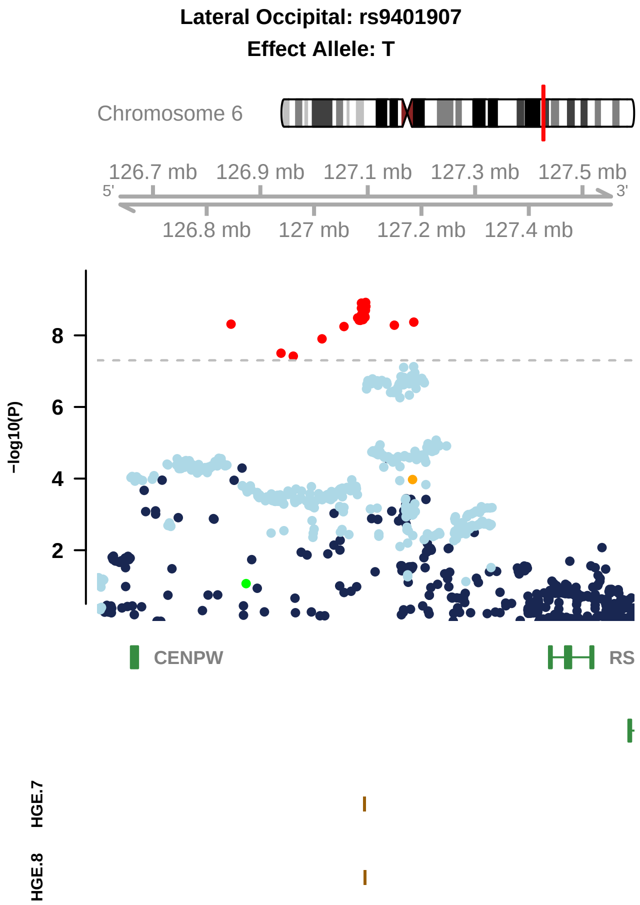
Lateral Occipital: rs9401907
Effect Allele: T
Chromosome 6
126.7 mb
126.9 mb
127.1 mb
127.3 mb
127.5 mb
5'
3'
126.8 mb
127 mb
127.2 mb
127.4 mb
-log10(P)
8
6
4
2
CENPW
RS
HGE.7
HGE.8

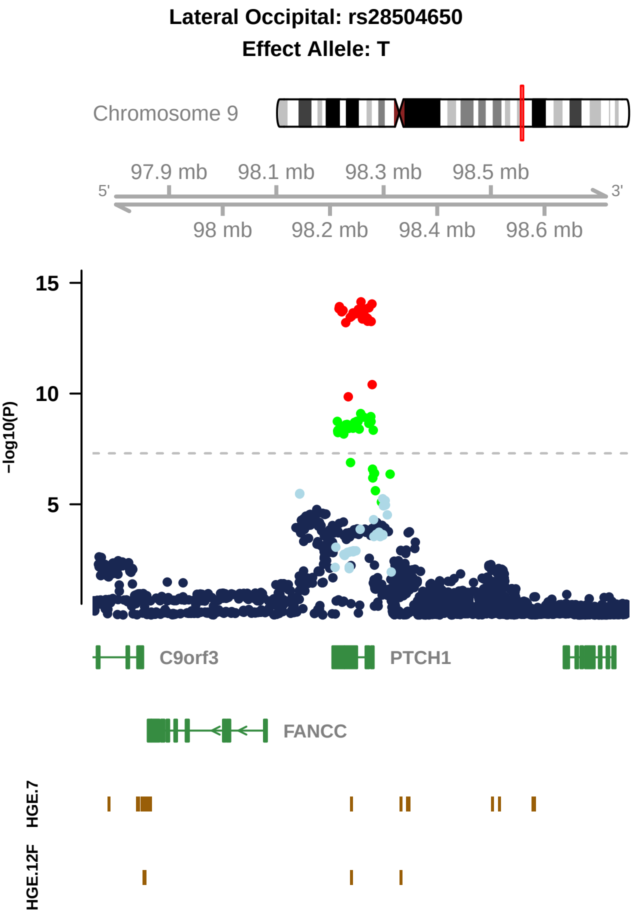

Lateral Occipital: rs28504650
Effect Allele: T
Chromosome 9
97.9 mb
98.1 mb
98.3 mb
98.5 mb
5'
3'
98 mb
98.2 mb
98.4 mb
98.6 mb
15
10
5
−log10(P)
C9orf3
PTCH1
FANCC
HGE.7
HGE.12F
